# A gene program association study (GPAS) delineates novel patterns of lineage plasticity and aggressiveness in non-small cell lung cancer

**DOI:** 10.1101/2025.05.06.652317

**Authors:** Rui Hong, Pascal Klöckner, Nageen Zahra, Sarah Mazzilli, Avrum E. Spira, Jennifer Beane, Masanao Yajima, Joshua D. Campbell

## Abstract

Combinations of molecular programs can be altered in cancer cells and contribute to tumor initiation and aggressiveness. We developed a novel framework called a Gene Program Association Study (GPAS) to create an atlas of gene co-expression modules in cancer cells from 127 primary non-small cell lung cancers (NSCLCs) and determine their association with clinical and histopathological features. Broadly expressed housekeeping modules were explained by study-specific effects while other categories of modules were better explained with other biological or clinical factors. Lung adenocarcinoma (LUAD) displayed extensive lineage infidelity characterized by heterogeneity in expression of modules from different alveolar cell types. Modules associated with late-stage LUAD were found in cells from early-stage tumors suggesting that aggressive phenotypes can be observed before clinical progression. Basal-and squamous-like cells were identified as possible intermediates in the formation of invasive mucinous LUAD. In lung squamous cell carcinoma (LUSC), a novel subtype was identified with lower levels of canonical squamous modules and higher levels of fibrinogen. LUSC-associated modules were also observed in normal basal cells exposed to cigarette smoke suggesting that this cell state is the more proximal “cell-state-of-origin” than unexposed basal cells. Overall, this atlas elucidated the combinations of molecular programs in cancer cells that contribute to heterogeneity and aggressiveness in NSCLC.

**Teaser:** We created an atlas of gene modules in NSCLC cells that defined late-stage disease, lineage plasticity, and novel subtypes.

## INTRODUCTION

Lung cancer remains one of the leading causes of death from cancer around the world. Despite advances in diagnosis and treatment, the 5-year survival rate remains around 20%^1^. Cancer stage at diagnosis remains the most important prognostic factor in determining survival and is used to determine the appropriate treatment strategy for each patient. Early-stage patients with localized disease have a five-year survival rate over 60% while patients with distant metastatic lung cancer have a much lower rate at 7%^1^. Patients within each stage also experience a wide variety of clinical outcomes with 30-50% of stage I patients ultimately recurring^2^. Another major challenge in the clinical management of lung cancer is the lack of standardized diagnostic and treatment strategies for subsolid nodules, including ground-glass opacities (GGOs)^3^, focal hazy opacifications in which the underlying lung structures remain visible. GGOs are now commonly detected with the increased use of low dose computed tomography (LDCT) for screening and may contain pathologically benign, premalignant, or invasive components^4,5^. Non-small cell lung cancer (NSCLC) is the most common type of lung cancer and is largely comprised of two major histological categories: 1) lung adenocarcinoma (LUAD) which is found more often in the peripheral alveolar space and thought to arise from alveolar type II cells and 2) lung squamous cell carcinoma (LUSC) which likely originates from basal cells along the central airways^6^. While some histological, mutational, and expression subtypes based on bulk assays have been described for both premalignant and invasive LUADs and LUSCs^7–12^, the spectrum of molecular programs in cancer cells that drive progression from premalignant to invasive to advanced disease or that distinguish between histological subtypes have not been characterized in a large-scale fashion.

Advances in single cell RNA-seq (scRNA-seq) assays have enabled the transcriptome profiling of individual cells from complex tissues including NSCLCs^11–23^. Despite the detailed resolution, the high cost and cumbersome nature of experimental protocols have prohibited the use of scRNA-seq assays in a large-scale fashion. The published scRNA-seq studies containing NSCLC only have a median of 7 samples, which decreases the ability to obtain statistically robust conclusions regarding associations with patient-level characteristics such as stage. A major computational hurdle in characterizing the diversity of molecular states in NSCLC is the lack of systematic methods to analyze the transcriptome of cancer cells across patients and studies. Each cancer cell can have a unique combination of genetic alterations, epigenetic changes and cell-cell interactions from the tumor microenvironment (TME) that influence its transcriptome. Standard clustering approaches for single cell data will often result in many patient-specific clusters of cancer cells without a clear strategy for defining what is shared or unique across clusters. Existing integration approaches for single-cell data generally assume that the same cell types or cell states exist across samples and datasets and will attempt to globally align cells in a common reduced dimensional space. While these approaches may be useful for analyzing shared immune and stromal cell states, they fail to capture the diversity and combinations of gene programs that are active in cancer cells within and across tumors. Non-negative matrix factorization (NMF) has been used to discover gene expression programs in single-cell RNA-seq data^24^. However, the factorization itself does not quantify how much of a program’s variability arises from technical versus biological sources, and associations with patient characteristics must be tested by a separate downstream analysis. To address the substantial technical and biological variation across samples and studies in cancer, recent approaches derive programs within each sample and then reconcile recurrent programs into consensus meta-programs^25,26^. Because factorization is performed on within-sample relative expression, this strategy restricts the recoverable programs to axes of intra-tumoral variation, requires that a program recur in multiple samples to be retained, and defines program number, membership, and granularity through user-specified similarity thresholds. Furthermore, this approach yields programs without an estimate of how much each contributes to technical versus biological variation.

To overcome these challenges, we developed a novel framework to perform a Gene Program Association Study (GPAS) and applied it to cancer cells from 127 NSCLCs obtained across 12 scRNA-seq studies. The major components of the GPAS framework include: 1) the identification of interpretable gene co-expression modules using a Bayesian co-clustering factorization tool called Celda, and 2) a systematic dissection of variability from technical and biological sources at the module level using linear mixed effect (LME) modeling. This strategy allowed for the identification of modules associated with histology and stage, detailed description of lineage infidelity and plasticity in LUAD, characterization of novel subtypes and the more proximal “cell-state-of-origin” in LUSC, and elucidation of which classes of modules are driven by genetic alterations.

## RESULTS

### Detection and characterization of gene co-expression modules in NSCLC

We collected data from 12 studies that applied scRNA-seq to tumor tissue collected from biopsies or resections from patients with NSCLC (**Supplementary Table 1**). This included 91 LUADs, 29 LUSC, and 7 NSCLCs without a more specific histological classification. Epithelial cells were identified using the published annotations or by assigning labels from reference atlases. Cancer cells were identified by identifying cells with detectable copy number alterations (**Methods**). After the initial annotation, two groups of cells with putative CNVs had high expression of either B-cell markers or ciliary markers. On further inspection, these cells were likely normal cells with false positive CNV calls due to co-localization of cell type specific genes on specific chromosomes (**Supplementary Figure 1**). After excluding these cells, a total of 114,069 malignant cells from 127 samples were used for module detection. Celda is a factorization tool that co-clusters cells into subpopulations, genes into co-expression modules, and estimates the levels of each module in each cell and cell population^27^. Unlike standard non-negative matrix factorization (NMF), Celda clusters each gene into a single module thus ensuring that all genes in each module have the same pattern of expression across cells (i.e. co-expression), which aids in interpretability and enhances identification of genes that are co-regulated. In total, we identified 175 gene co-expression modules and 80 cell clusters (**Supplementary Figure 2A**, **Supplementary Tables 2-4**). Module size varied from 1 to 892 genes per module (median = 18, mean = 79.2; **Supplementary Figure 2B**). Each module contained genes with the same expression pattern across cells and each module displayed a unique pattern of expression across cells compared to other modules (**Figure 1A**). For example, module L12 contained NKX2-1, a known marker of alveolar lineages and was predominantly expressed in LUADs. In contrast, the module L14 contained SOX2, a marker for squamous cells and was predominantly expressed in LUSCs. Other modules displayed heterogeneity within and across tumor types. For example, module L65 contained many MHC class II genes and tended to be more highly expressed in early stage LUADs; the detoxification module L35 was enriched for genes involved in xenobiotic metabolism and expressed in a subset of LUSCs; and module L43 was enriched for cell cycle genes and expressed in a subset of cells within tumors across LUAD and LUSC.

**Figure 1.**
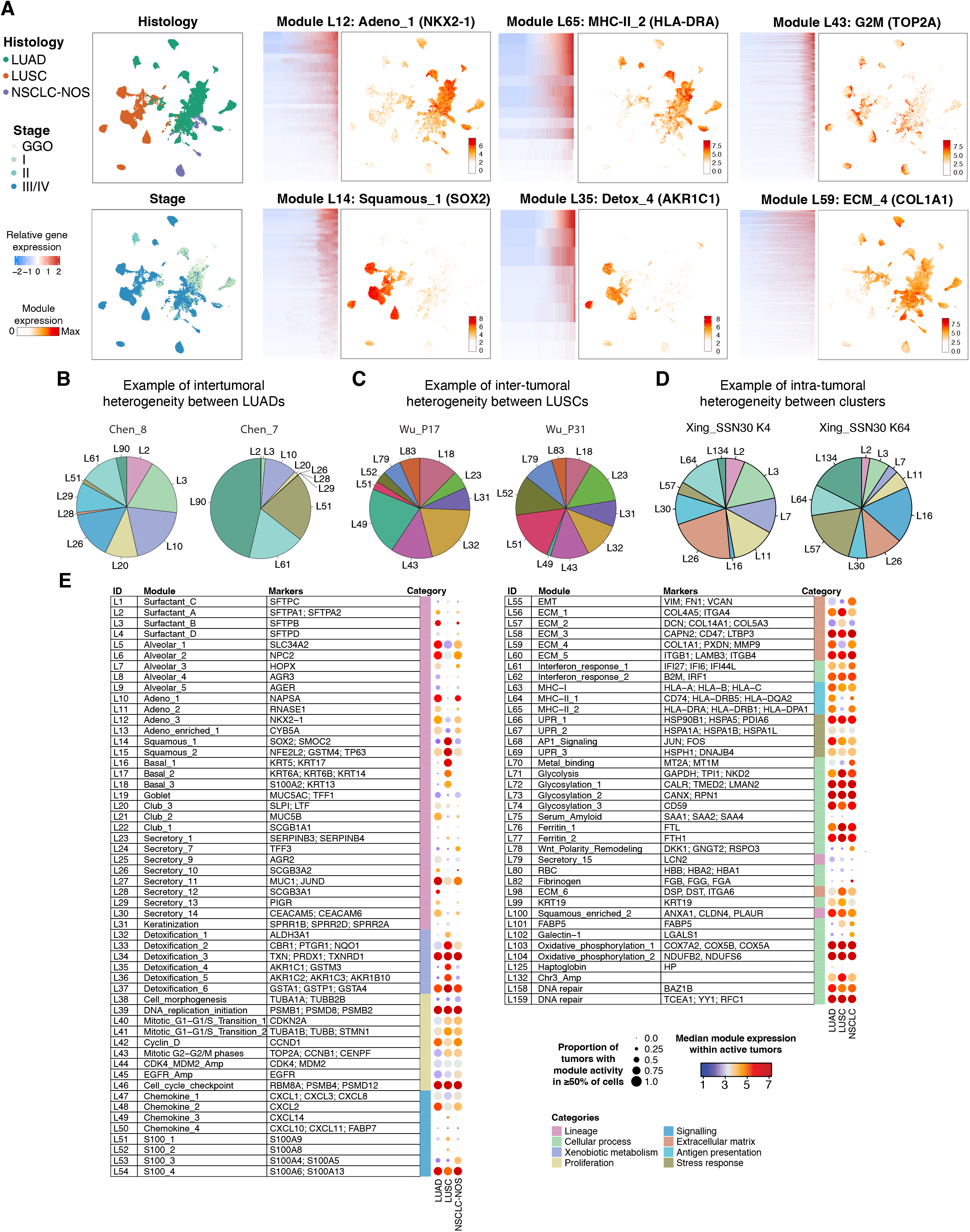
Detection of co-expression modules across tumors and studies. 114,069 cancer cells were identified from 127 NSCLCs across 12 studies by identifying copy number alterations. Celda was used for deconvolution of tumors to identify 175 co-expression modules across 80 cell clusters. Modules were further analyzed to understand variability from different sources, characterize patterns of lineage infidelity, and determine underlying genetic drivers of module expression. **(A)** Left: UMAPs are colored by tumor type and stage. Right: Six examples of co-expression modules are shown. Each heatmap shows the co-expression pattern of genes assigned to that module. Cells in the heatmap are ordered by those with the lowest to the highest module probability. Each UMAP shows a unique pattern of log2 normalized module expression across cells. **(B, C)** Each tumor is a unique composition of modules. The percentages of the top 10 most variable modules are shown for 2 LUADs and 2 LUSCs. **(D)** Additional heterogeneity can be observed across clusters within each tumor. The percentage of the top 10 most variable modules are shown for Xing_SSN30 across two cell clusters, K4 and K64. **(E)** The table lists the curated modules with corresponding key genes used for annotation. A module was considered “detected” in an individual cell if it had a log2 normalized expression value of at least 1. A module was considered active in an individual tumor if it was detected in at least 50% of cells. The size of the dot for each module indicates the proportion of activated tumors within LUAD, LUSC, or unspecified NSCLCs. The color of the dot indicates the median expression of the module within the activated tumors.

The primary reason for tumor-specific clusters is that each tumor expresses a unique composition of modules. To illustrate this heterogeneity, we examined the proportions of the top 10 most highly variable modules across the two major histological subtypes. In LUAD, Chen_8 and Chen_7 expressed high levels of adeno module L10 and interferon module L61. However, Chen_7 had higher levels of signaling module L51 and L90 containing SERPINA1 and lower levels of alveolar modules L2 and L3 (**Figure 1B**). In LUSC, Wu_P17 had higher proportions of L32:Detoxification_2 and L49:Chemokine_3 while Wu_P31 had higher proportions of the S100 signaling modules L51 and L52 (**Figure 1C**). This heterogeneity was also evident between different subpopulations within a tumor. For example, cells from Xing_SSN30 fell into two predominant clusters, K4 and K64 (**Figure 1D**). Subpopulation K4 expressed higher levels of alveolar/AT2 modules L3, L7, and L11, while subpopulation K64 displayed higher levels of the ECM module L57. Overall, these results show how the heterogeneity between tumors can be deconstructed by defining the combinations of co-expression modules.

The majority of modules were classified into one of 9 broad biological categories if they contained known marker genes for specific processes or cell types, or if they were enriched in specific functional categories (**Supplementary Table 5**). Categories included lineage, cell cycle and proliferation, extracellular matrix production (ECM) or epithelial-to-mesenchymal transition (EMT), antigen presentation, xenobiotic metabolism, stress response, chemokine signaling, housekeeping/ribosomal, and other general cellular processes (**Figure 1E**; **Supplementary Figures 3-5**). Several modules were enriched for genes related to cell cycle and proliferation which is a canonical hallmark of cancer^28^.

Some of these modules were highly expressed in a small number of tumors and are likely the result of focal amplification, such as L42 containing CCND1, L44 containing CDK4 and MDM2, and L45 containing EGFR and its neighboring gene SEC61G. To systematically identify modules that may reflect copy number alterations rather than co-regulated programs, we tested each module for enrichment of genes located within the same chromosomal cytoband (**Methods**; **Supplementary Table 6**). L44 was strongly enriched for genes from 12q14–q15 (FDR = 8 x 10^-13^), and L42 was modestly enriched for genes from 11q13 (FDR = 0.04), consistent with these modules capturing the CDK4/MDM2 and CCND1 amplicons, respectively. Other modules were broadly detected across the majority of tumors but were only highly expressed in subsets of cells within each tumor. These were reflective of the current state of the mitotic cycle for each cell and include L40 (CDKN2A), L41 (TUBA1B; TUBB; STMN1) and L43 (TOP2A, CCNB1, CENPF, MKI67). L39 and L46 were highly expressed across all cells and contained genes involved in DNA replication initiation and cell cycle checkpoint, respectively. Modules containing genes involved in ECM included L55 which contained markers for EMT (VIM, FN1, VCAN) as well as modules L56-L60 which contained different combinations of collagens, integrins, matrix metalloproteinases (MMPs), laminins, and decorin. The stress response module L68 was enriched in genes downstream of the AP-1 signaling pathway (JUN, FOS) while modules L66, L67, and L69 included many heatshock proteins (HSPs) from the unfolded protein response pathway (UPR). Modules enriched for signaling proteins included L47-L50 which contained various chemokine signaling genes (CXCL1, CXCL3, CXCL8, CXCL2, CXCL14, CXCL10 and CXCL11) as well as modules L51-L54 which contained many S100 calcium-binding proteins (S100A9, S100A8, S100A4, S100A5, S100A6, S100A13). Antigen presentation modules included L63 with MHC class I genes (HLA-A, HLA-B, HLA-C) and L64 and L65 which contained MHC class II genes (CD74, HLA-DRA, HLA-DRB1). Modules L32-L37 were enriched in functional categories such as xenobiotic metabolism, NFE2L2/KEAP1 pathway, and the glutathione pathway. Lastly, several modules were enriched for ribosomal genes or contained a large number of housekeeping genes.

### Dissecting biological and technical variability across samples and studies

A major statistical challenge when analyzing the transcriptomes of cells from multiple tumor samples and studies is how to best disentangle the variation driven by technical or biological factors. We devised an approach using linear mixed effect (LME) modeling to measure technical differences between studies as well as complex biological heterogeneity that arises due to inter-and intra-tumor variation (**Figure 2A**; **Supplementary Table 7**). This approach also accounts for the fact that cells from the same tumor are not independent from one another. Each cell is treated as a repeated measure in fully nested random effects, which include terms for Study, Sample (i.e. each tumor), and Cluster (i.e. different subpopulations within each tumor). The model also includes fixed effects for histology (LUAD vs. LUSC) and Stage (Subsolid/GGO and Stage I vs. Stage II-IV). The samples in the Subsolid/GGO group were mostly derived from Stage I patients except for 2 GGO samples adjacent to the invasive tumor that were taken from a Stage II patient in Yanagawa *et al* (Case4). When examining the variability explained by each variable, we found that 24 modules had over 20% of the variance explained by the Study covariate (**Figure 2B**). 163 modules had less than 30% explained by Study including 92 modules with at least 5% of variability explained by the fixed effects. Different sources of variability were more influential for different types of biological processes (**Figure 2C**; **Supplementary Figure 6**). For example, lineage, detoxification, and antigen presentation modules had a higher percentage of variability explained by the Histology and Stage fixed effects compared to the Study effect. In contrast, the broadly expressed housekeeping, ribosomal, and some unannotated modules had the highest percentage of variability explained by the Study term.

**Figure 2.**
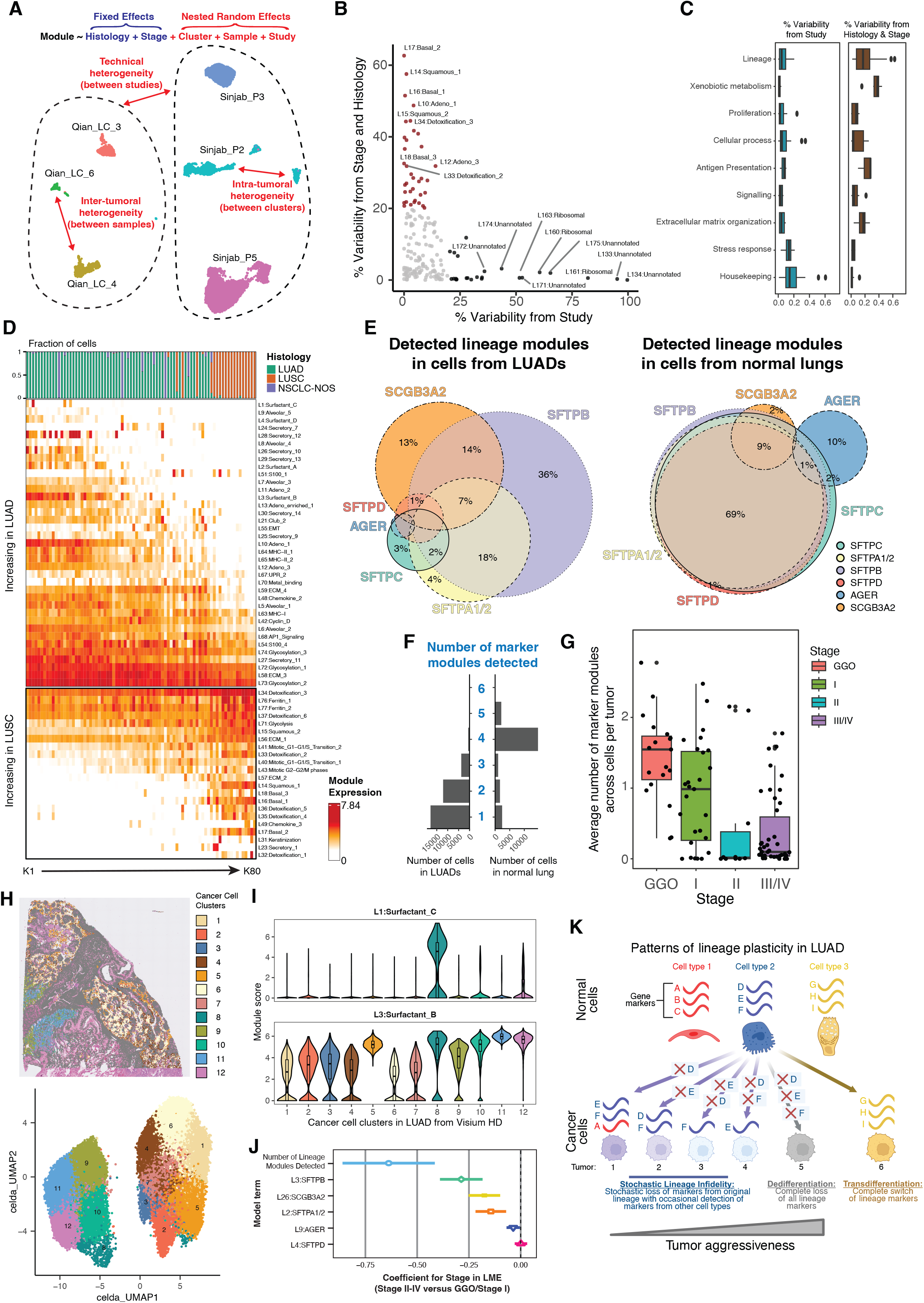
Dissecting variability from biological and technical sources enables characterization of lineage infidelity across samples and studies. **(A)** Linear mixed effect (LME) modeling was used to quantify the variability and association of each module with biological and technical covariates. Each cell was treated as a repeated measure using Study, Sample, and Cluster as fully nested random effects while Histology and Stage were included as fixed effects. **(B)** The percentage of variability explained by the Study effect is compared to the Histology and Stage effects for each module. **(C)** Large unannotated modules or modules enriched for ribosomal genes were largely driven by Study. In contrast, modules from other biological categories such as lineage, xenobiotic metabolism, and antigen presentation were more strongly explained by Histology and Stage. **(D)** LMEs identified modules significantly different between LUADs and LUSCs (FDR q-value < 0.05; |Log2 Fold Change| > 0.6). Each row in the heatmap corresponds to a module and each column corresponds to a cell cluster. The stacked barplot shows the fraction of cells with each histology within each cluster. **(E)** Left: Euler diagram showing the overlap in the modules detected in LUAD cells. Modules containing known AT2 (i.e. surfactants) or AT1 markers (i.e. AGER) were selected. Nearly every combination of detected modules could be found in a subset of cells when examining the surfactant modules (L1-L4) and the SCGB3A2 module (L26). Right: Euler diagram showing the overlap in modules for AT2 and AT1 cells from a normal lung dataset by Basil *et al*. In contrast to cancer cells, most AT2 cells expressed all AT2-related surfactant modules while most AT1 cells expressed the module containing the known AT1 markers. **(F)** The number of modules with AT2 or AT1 markers was quantified in each cell from LUADs and compared to alveolar cells from normal lung tissue. In LUAD, the majority of cells had detectable levels of one or two AT2 modules. In contrast, the majority of AT2 cells from normal lung tissue expressed four AT2 modules. **(G)** The average number of detected marker modules was calculated across cells per tumor. The number of detected marker modules was significantly lower in LUADs from late-stage patients (GGO and Stage I versus Stage II-IV; p = 2.07e-08; LME). (**H**) Cancer cells in a spatial transcriptomics dataset of a LUAD profiled with Visium HD were identified with inferCNV and grouped into 12 clusters. Top: Cancer cell clusters are shown at their location within the tissue section, with non-cancer cells shown in gray. Bottom: UMAP of cancer cells colored by cluster. (**I**) Violin plots of module expression for L1:Surfactant_C (SFTPC) and L3:Surfactant_B (SFTPB) across the 12 cancer cell clusters. L3 was highly expressed in several clusters and detected at intermediate to low levels in the others, whereas L1 was largely restricted to a single cluster. **(J)** Coefficients for stage along with 95% confidence intervals are shown for different LMEs with different response variables. The model with the number of detected modules as the response had a stronger association with stage compared to other LMEs with expression of individual modules as the response. **(K)** Patterns of lineage infidelity in LUAD suggest that the order in which AT2 markers are lost is not deterministic but can vary from tumor to tumor and that the cumulative loss of markers is more associated with tumor aggressiveness than loss of any individual marker. This infidelity can occasionally be accompanied with an increase in expression of an AT1 marker but does not result in complete transdifferentiation to AT1 cells.

To characterize which modules vary most between samples, we examined the modules with the highest percentage of variance attributable to the Sample effect in the LME (**Supplementary Table 8**). Most corresponded to biological subpopulations rather than technical artifacts. Several reflected signaling modules, including L50:Chemokine_4 (CXCL10, CXCL11, FABP7), L53:S100_3 (S100A4, S100A5), and L54:S100_4 (S100A6, S100A13), as well as the interferon-response module L61. Module L45:EGFR_amp corresponded to EGFR amplification and was detected in only 2 samples. Other modules denoted rare patient subgroups. For example, L19:Goblet (MUC5AC) marked a goblet subpopulation found in four LUADs, two of which were pathologically annotated as minimally invasive adenocarcinomas (MIAs), and L82:Fibrinogen was found in subsets of cells from 5 LUSCs. Both of these modules also varied by cluster, indicating that they distinguish subsets of cells within a tumor as well as tumors within a study. In contrast, one module reflected a likely technical artifact of tissue dissociation. L80:RBC contained several red blood cell marker genes (HBB, HBA2, HBA1), consistent with variable red blood cell lysis and the resulting ambient contamination of cancer cells across samples. Thus, while most modules with high sample-level variance denoted genuine biological subgroups, this analysis also identified the modules attributable to sample-level technical effects, such as L80.

We also performed an analysis examining the average effect size (i.e. magnitude of change) for the possible sources of variability in each module. For example, module L10 had an average log2 difference of 1.94 (corresponding to a 3.84-fold change) between early-and late-stage LUAD and an average log2 difference of 5.27 (corresponding to a 38.58-fold change) between early-stage LUADs and LUSCs (**Supplementary Figure 7A-C; Supplementary Table 9**). This outweighed the average differences between studies, samples, clusters, and cells, which all had less than a log2 difference of 1 on average (corresponding to a fold change of 2). When expanding the analysis and counting modules with an absolute log2 difference greater than 1, we found that histology affected the largest number of modules (n=79) followed by Stage (n=15) (**Supplementary Figure 7D**). Study was the third largest factor with 10 modules having an average difference greater than 1. Additionally, we could use the estimates of the LME model to subtract out the effect of Study on each module while preserving the other biological effects. A UMAP generated with all modules and the uncorrected module expression shows that the largest global difference was between Wu *et al* and other studies, which likely represents the effect of different scRNA-seq platforms (**Supplementary Figure 7E**). In contrast, when using the corrected module expression and limiting to modules with <25% variability from the Study effect, the largest global difference was between LUADs and LUSCs. Overall, these results demonstrate that the technical variability induced by study-specific differences largely affects broadly expressed housekeeping genes and that the remaining modules enriched for other processes have larger effects from biological factors such as Histology and Stage.

### Stochastic lineage infidelity is prominent in cancer cells from LUAD

Lineage infidelity in the context of lineage plasticity is where cancer cells lose their original cellular transcriptional identity and adopt a more primitive, undifferentiated state or transdifferentiate to a different cell type altogether^29^. To systematically quantify infidelity within and across tumor types, we identified modules significantly associated with histology using the LMEs (FDR q-value < 0.05; |Log2 Fold Change| > 0.6; Percentage of cells detected in LUAD or LUSC >30%) (**Figure 2D**; **Supplementary Table 7**). Modules up-regulated and more broadly expressed in cells from LUAD compared to LUSC include those with known alveolar markers such as L10 (NAPSA), L11 (RNASE1), and L12 (NKX2-1). Other modules significantly higher in the LUAD cohort but expressed to varying degrees across individual LUADs included AT2 surfactant markers such as L1 (SFTPC), L2 (SFTPA1/SFTPA2), L3 (SFTPB), and L4 (SFTPD). Some modules were broadly detected across both tumor types but were significantly higher in the LUAD cohort including those involved in ECM/EMT (L55, L58, L59), glycosylation (L72, L73, L74), and stress response (L67, L68). Recent studies have implicated AT1 cells as a potential cell-of-origin for lepidic lung cancer^30^ and plasticity to AT1-like cells as a mechanism of resistance to KRAS inhibition^31^. To explore the possible existence of an AT1 phenotype in primary, untreated LUADs, we first identified a module of genes defining normal AT1 cells using a published scRNA-seq dataset of normal lung tissue. This module contained known AT1 markers such as HOPX, AGER, RTKN2, and CAV1 (**Supplementary Figure 8**, **Supplementary Table 10**)^32^. While these markers were detected in our dataset, they were spread across different modules (L7, L9, L60, L115) and intermixed with genes unrelated to alveolar lineage specification. Additionally, these modules were detected in cells that also expressed other modules with AT2 markers. To assess AT1 and AT2 identity independently of module assignments, we examined individual AT1 and AT2 markers (**Supplementary Figure 9A**). No cancer cell cluster expressed the specific AT1 markers AGER and RTKN2 at levels approaching those of normal AT1 cells, and clusters with the lowest expression of AT2 markers showed no corresponding gain of AT1 markers (**Supplementary Figure 9B**). Heterogeneity in AT2 marker expression was also evident among clusters within individual tumors (**Supplementary Figure 9C**). We next scored each malignant cell with AT1 and AT2 signatures using AUCell, with AT1 and AT2 cells from three normal lung datasets as references (**Supplementary Figure 10A**). Cells were classified as AT1-like or AT2-like if their scores fell within the range of the corresponding normal reference cells (**Methods**). Across the 91 LUADs, only 0.3% of malignant cells were AT1-like and 0.6% were AT2-like across the atlas (**Supplementary Figure 10B**). AT1-like cells were rare in invasive tumors but constituted the predominant state in a single GGO lesion (Yanagawa Case1_GGO). Within individual tumors, clusters varied widely in AT2 score while AT1 scores remained uniformly low (**Supplementary Figure 10C**). Thus, malignant cells in LUAD largely occupy a partially differentiated alveolar state, and loss of AT2 identity is generally not accompanied by a coordinated switch to an AT1 program.

We also observed that different LUAD-related lineage modules displayed varying levels of detection across cells (**Figure 2E**, **Supplementary Table 11**). For example, the module containing SFTPC, a commonly used marker for AT2 cells, was only detected in 2,795 cells while the SFTPB module was detected much more broadly in 27,234 cells. Despite this trend, 2.7% of cells expressed the SFTPC module without the SFTPB module. In fact, many different combinations of modules could be detected in a subset of cells when examining the surfactant modules (L1-L4) and SCGB3A2 (L26). These combinations were observed among cell clusters within individual tumors and not only between tumors (**Supplementary Figure 9C**). This degree of heterogeneity did not exist in the normal lung where 74% of the alveolar cells expressed all surfactant modules (i.e. AT2 cells) without detection of the AT1 module, 10% of cells exclusively expressed the AT1 module without the surfactant modules (i.e. AT1 cells), and only 7% of cells expressed both surfactant and AT1 modules above background. The remaining ∼9% expressed some, but not all, surfactant modules (**Figure 2F, Supplementary Table 12**). Similar patterns were observed in a spatial transcriptomics dataset of a LUAD profiled with Visium HD (Methods). Cancer cells were identified with inferCNV and grouped into 12 clusters that were spatially organized within the tumor (**Figure 2H**; **Supplementary Figure 11A,B**). Module L3:Surfactant_B (SFTPB) was highly expressed in several cancer cell clusters and detected at intermediate to low levels in the others, whereas L1:Surfactant_C (SFTPC), L2:Surfactant_A, and L4:Surfactant_D were largely restricted to a single cluster (**Figure 2I**; **Supplementary Figure 11C**), demonstrating that different combinations of surfactant modules coexist within a single tumor independently of tissue dissociation and droplet-based scRNA-seq.

Lastly, the heterogeneity in AT2 module expression was also associated with clinical stage. Specifically, the expression of some AT2 and secretory modules was significantly lower in late-stage LUAD including L2:Surfactant_A, L3:Surfactant_B, and L26:Secretory_10 (LME; FDR q-value < 0.05). Beyond the expression of individual modules, the overall number of AT2 lineage modules detected was also significantly lower in cells from late-stage LUAD (p = 2.07e-08; **Figure 2G**). Interestingly, the number of lineage modules detected per cell had a stronger association with stage than the expression of any individual module alone as demonstrated by a more negative coefficient from the LMEs (**Figure 2J**). In summary, these patterns suggest that the order in which AT2 markers are lost is not deterministic but stochastic and can vary from tumor to tumor (**Figure 2K**). Stochastic lineage infidelity may also result in the occasional up-regulation of a marker for another cell type (e.g. AT1), but this does not necessarily indicate full transdifferentiation – a process in which all markers for the original cell type are lost and markers for another cell type are coordinately gained. Moreover, our data suggest the cumulative loss of markers in any order in LUAD may be more important for predicting clinical phenotypes of aggressiveness such as stage rather than the expression of individual markers.

### The landscape of gene modules associated with stage in LUAD

Our combined cohort contained LUADs along a spectrum of disease development including premalignant lesions obtained from GGOs to primary tumors from patients with late-stage disease (i.e. had a concurrent metastatic lesion in a lymph node, the other side of the lung, or another organ). In order to understand the molecular programs associated with disease progression, we examined all the gene modules that were differentially expressed between early-stage tumors (lesions found in GGOs and tumors from Stage I patients) compared to tumors from late-stage patients (stages II-IV; **Figure 3A**; **Supplementary Table 13**). Several additional lineage-related modules that distinguished LUADs from LUSCs were down-regulated in LUAD cells from late-stage patients. Beyond the previously mentioned AT2 lineage modules (L2, L3), the broader adeno modules (L10-L12) and general secretory modules (L26, L28, L29) were also down-regulated reflecting that the continued loss of cellular identity and dedifferentiation is associated with increased risk of aggressive disease (FDR q-value < 0.05; |Log2 Fold Change| > 0.55; LME)^33^. Interestingly both MHC class II modules were also down-regulated in tumors from late-stage patients suggesting a potential mechanism of immune evasion necessary for metastasis^34^. Conversely, several modules were increasing in expression among later-stage tumors. Modules related to mitosis and proliferation were increasing in cells from patients with late-stage disease (L40, L41, L43). Although larger numbers of proliferating cells can be readily observed in lung cancers^35^, our analysis demonstrates that the higher transcriptional levels of these genes can also be observed within the cancer cells of late-stage tumors. Module L71:Glycolysis (GAPDH, TPI1, NKD2), ECM modules L59:ECM_4 (COL1A1, PXDN, MMP9), L60:ECM_5 (ITGB1, LAMB3, ITGB4), and L98:ECM_6 (DSP, DST, ITGA6) as well as interferon response module L61 (IFIT1-3) also had significantly higher expression in cells from late-stage tumors. Chemokine module L50 (CXCL10, CXCL11) was specifically expressed in clusters K54 and K56 which predominantly contained cells from 7 stage II tumors across 3 studies. L50 also contained high expression of FABP7 which has been associated with lung metastasis and can modulate invasive phenotypes^36^. L82 contained fibrinogen-related genes (FGB, FGG, FGA) as well as one neuroendocrine marker, CALCA (also known as CGRP). It was highly expressed by cells in clusters K32 and K34 which predominantly contained cells from 3 patients across 2 studies. Fibrinogen is a coagulation factor normally produced by the liver and circulates in the blood. While its role as a serum biomarker for prognosis of lung cancer has been previously examined^37^, the expression of fibrinogen by lung cancer cells to promote metastasis has not been explored. L78 was expressed in 2 clusters of cells (K49 and K63) and contained the gene HMGA2 which has been associated with lung metastasis^38,39^. These clusters of cells also lacked expression of the alveolar modules L10-L12 corroborating previous evidence that loss of NKX2-1 can contribute to de-repression of HMGA2 and lead to more aggressive disease^39^.

**Figure 3.**
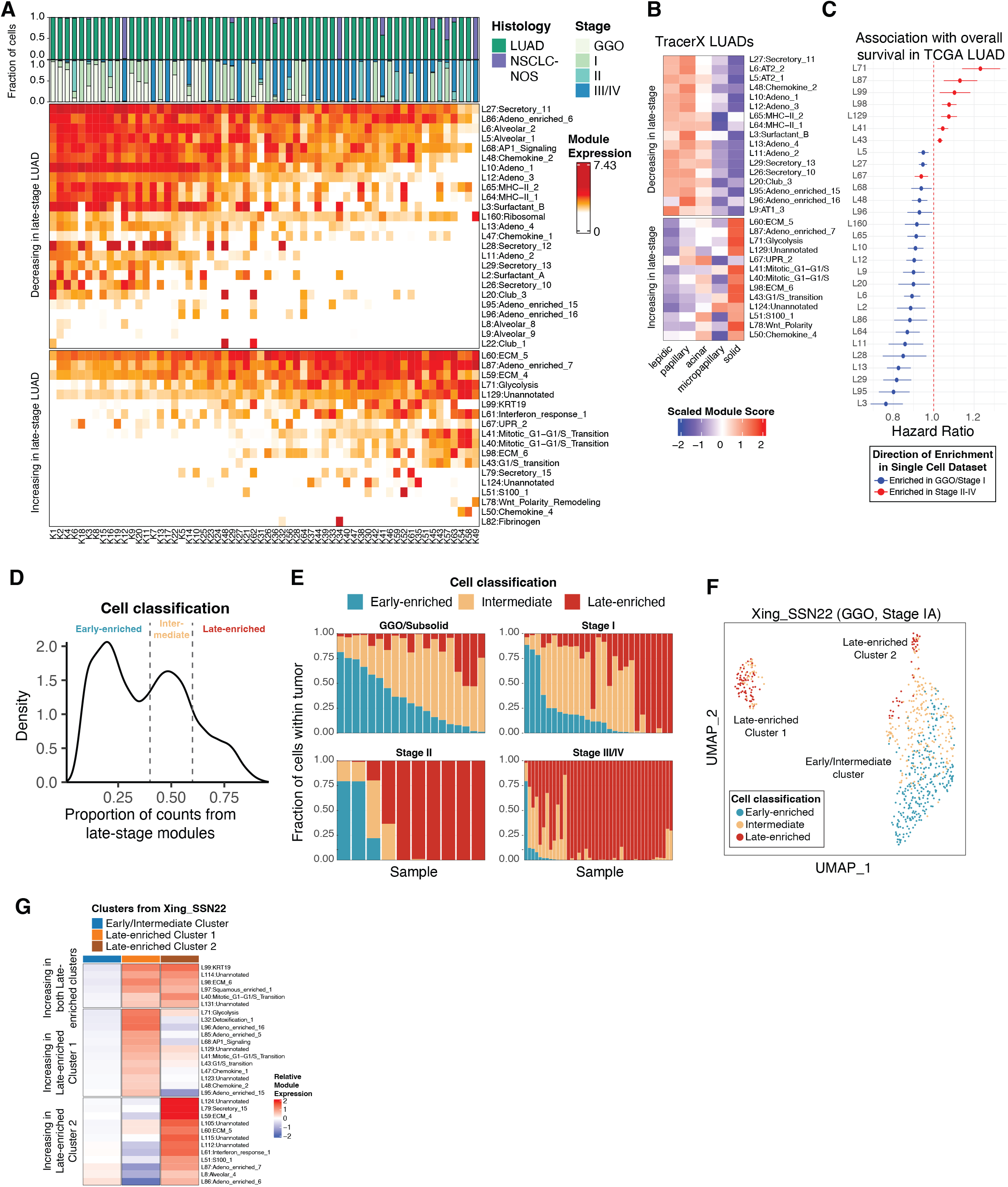
Modules associated with late-stage LUAD are heterogeneous in early-stage tumors. **(A)** LMEs identified modules associated with Stage in LUAD cells (FDR q-value < 0.05; |Log2 Fold Change| > 0.55). Each row in the heatmap corresponds to a module and each column corresponds to a cell cluster. The stacked barplot shows the fraction of cells from each histology or stage within each cluster. **(B)** Modules were scored in the TracerX bulk RNA-seq dataset and associated with LUAD subhistologies using ANOVA (FDR < 0.05). 17 of 25 early-stage associated modules were enriched in tumors with less aggressive subhistologies (i.e. lepidic and papillary) and 13 of the 18 late-stage associated modules were enriched in tumors with more aggressive subhistologies (e.g. micropapillary, solid). **(C)** Modules were scored in the TCGA LUAD dataset and associated with overall survival using a Cox proportional hazards model controlling for sex and age. 21 of 25 early-stage modules and 7 of 18 late-stage modules were concordantly associated with better or worse overall survival, respectively (FDR < 0.05). 1 late-stage module was discordantly associated with better overall survival. **(D)** The fraction of counts derived from late-stage associated modules out of the total number of counts from both early-and late-stage associated modules was calculated. Cells were divided into 3 groups based on the fraction of late-stage counts: cells containing a low fraction were deemed “early-enriched” (<0.40), those containing medium levels of late-stage counts were deemed “intermediate” (0.40-0.60), and those containing a high fraction of late-stage counts were deemed “late-enriched” (>0.60). **(E)** The distributions of early-enriched, intermediate, or late-enriched cells are shown in a stacked barplot for each LUAD. Within each stage, tumors are ordered from those with the highest fraction of “early-enriched cells” to the least. **(F)** Subsolid nodule SSN22 from Xing *et al* contained one population of early-enriched/intermediate cells and two distinct populations of late-enriched cells. **(G)** Modules such as L40 were commonly higher in both late-enriched cell populations compared to the early-enriched population. Modules such as L59:ECM_4, L60:ECM_5, and L61:Interferon_response_1 were specifically higher in the second late-enriched cell population.

To validate the relationship of these modules with aggressive phenotypes, we scored each module in the bulk RNA-seq data of TracerX LUADs and determined their association with histological subcategories of LUAD^40^. 17 of the 25 early-stage associated modules were significantly up-regulated in tumors with predominantly lepidic and papillary histologies which tend to have favorable outcomes^41^ (**Figure 3B**). Similarly, 13 of the 18 late-stage associated modules were up-regulated in micropapillary and solid tumors which have a poorer prognosis. Next, we determined the relationship between these modules and overall survival in the TCGA LUAD cohort. 21 of 25 early-stage modules were associated with better overall survival using a Cox proportional hazards model while controlling for sex and age as covariates (FDR q-value < 0.05; **Figure 3C**). 7 of the 18 late-stage modules were concordantly associated with worse survival while 1 late-stage module (L67) was discordantly associated with better survival. In general, the concordance of stage-associated modules from the single cell atlas with pathological and clinical phenotypes from independent bulk RNA-seq datasets supports the validity of the GPAS framework.

We next sought to understand the extent of heterogeneity of the early-and late-stage associated modules within and across tumors. The percentage of counts from genes in either early-or late-stage associated modules (FDR < 0.05) was calculated for each cell and cells were categorized as “early-enriched”, “intermediate”, or “late-enriched” based on trimodality observed in this distribution of percentages (**Figure 3D**). The distribution of early-enriched, intermediate, and late-enriched cells was then examined across tumors and stages (**Figure 3E**). While some tumors contained predominantly early-or late-enriched cells, other tumors contained a mix of these three categories. 3 of 19 lesions from GGOs (16%) and 11 of 27 tumors from stage I patients (41%) contained at least 30% of late-enriched cells. For example, the sample SSN22 from Xing *et al* was a subsolid nodule that contained three different clusters of cells (**Figure 3F**). One cluster contained predominantly early-enriched and intermediate cells while the other two clusters contained late-enriched cells. Differential expression analysis revealed that some late-stage associated modules such as cell cycle module L40:Mitotic_G1_G1/S_Transition_1 and L99:KRT19 were common across both sets of late-enriched cell clusters (**Figure 3G**). However, other late-stage associated modules were specific to only one late-enriched cluster. For example, cell cycle modules L41 and L43 were higher in the first cluster of late-enriched cells while L59: ECM_4 and L60:ECM_5, L61:Interferon_response_1, and L51:S100_1 were higher in the second cluster of late-enriched cells. Although this tumor was classified as Stage I, it contained many cells with high expression of modules associated with more aggressive tumors suggesting that it may be at higher risk of progression. Overall, these results demonstrate that transcriptional phenotypes associated with more aggressive LUAD can display extensive intratumoral heterogeneity and may be useful for prognostication of early-stage tumors.

### Basal-and squamous-like intermediates in mucinous LUADs

Invasive mucinous adenocarcinomas (IMAs) are a histologically defined subset of LUADs with distinct patterns of goblet cell differentiation^42^. Cell clusters K59 and K61 had high expression of module L19 which contained MUC5AC, a known marker for the goblet cell lineage (**Figure 4A**). These clusters predominantly contained cells from 4 LUADs (**Supplementary Table 4**), two of which were the only LUADs pathologically classified as IMA. 3 of these LUADs also contained other cell clusters with different lineage-related expression. Sinjab_P2 was the most complex and contained different subpopulations with high expression of alveolar modules (L2, L3, L11, L12), squamous and basal modules (L14, L16, L18), airway club modules (L20-L22), or the mucinous module (L19) (**Figure 4B, C**). Bischoff_p030t contained two clusters of cells, one with high expression of L19 (MUC5AC) and the other with significantly higher expression of modules related to EMT (L55), cell cycle and proliferation (L38, L41), basal cell expression (L17), fatty acid binding (L101)^43^, and Galectin-1 (L102) when compared to other LUAD cells from the same dataset. The intermediate cells from Bischoff_p033t also had higher levels of L101 and L102. While Zilionis_P3 only had mucinous cells with high expression of L19, these cells also had significantly higher expression of the basal module L16 compared to other LUADs in the same dataset. Goblet cells from 3 of these tumors had higher levels of the Squamous_2 module (L15). Within each of these tumors, the cancer cells shared broad copy number alterations, indicating that the alveolar-, intermediate-, and mucinous-like populations arose from a common ancestor rather than from independent clones (**Supplementary Figure 12A**). The mucinous module L19 was not enriched for genes from any single chromosomal region (**Supplementary Table 6**), indicating that its expression reflects a lineage program rather than a direct consequence of a copy number alteration. CytoTRACE analysis revealed that the intermediate cells had higher “stemness” compared to club-like and mucinous cells, consistent with the intermediate cells being the most progenitor-like state within these tumors (**Supplementary Figure 12B**). To validate these associations, we scored each module in the TracerX LUAD bulk RNA-seq dataset and determined which modules were significantly higher in IMA tumors compared to other subhistologies (**Figure 4D**). In addition to secretory-related modules L19, L21, L24, L25, and L79, we also found that the basal module L16 was significantly higher in IMAs. Within the IMA tumors (n = 35), L14, L15, L16, L17, and L79 were significantly positively correlated with L19 further supporting the association of squamous and basal-like phenotypes in IMA histology (**Figure 4E**).

**Figure 4.**
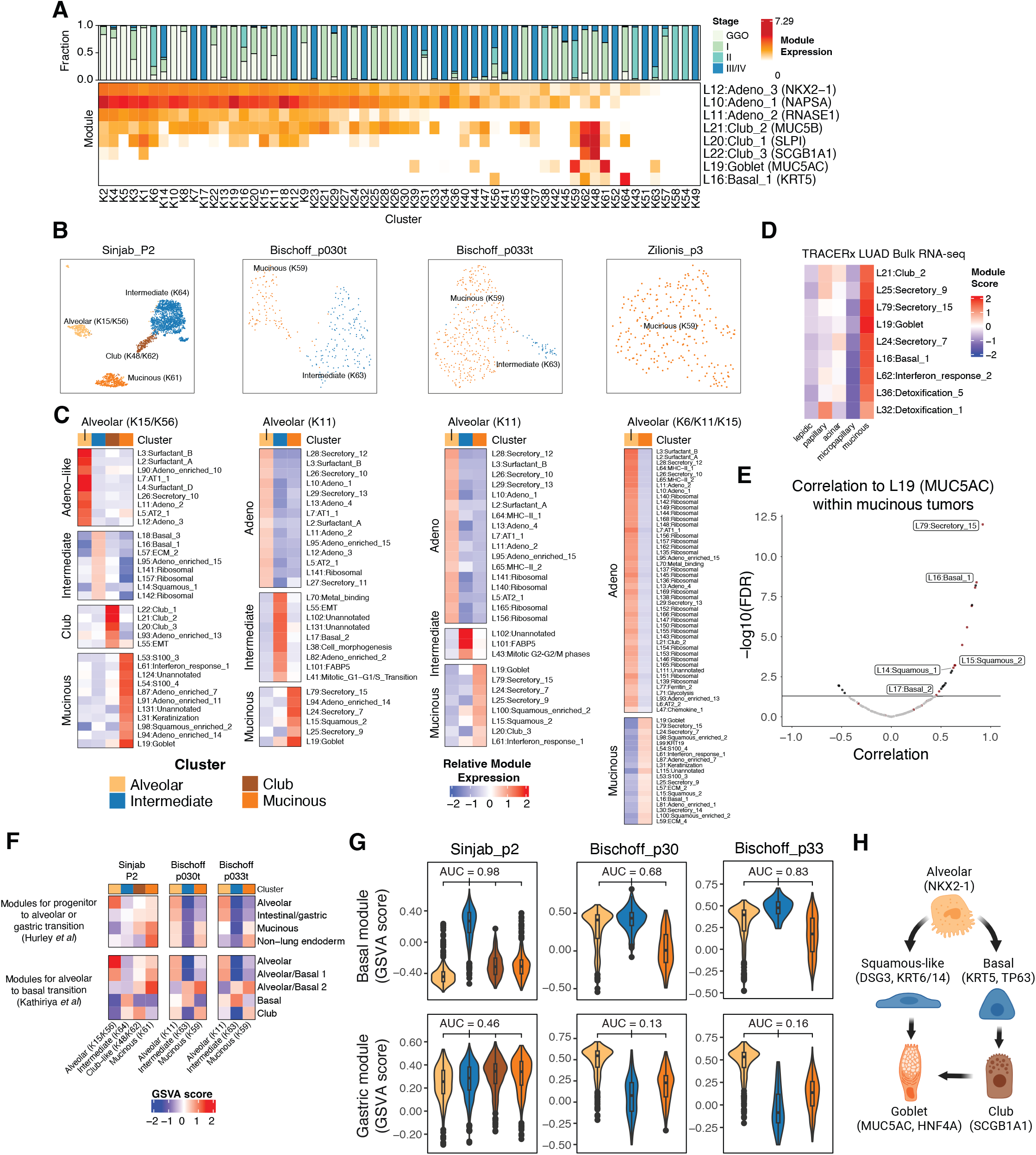
Squamous-and basal-like intermediate cell populations in invasive mucinous adenocarcinomas (IMAs). **(A)** The heatmap shows the expression of adenocarcinoma-and mucinous-related modules for LUAD cells. Each row in the heatmap corresponds to a module and each column corresponds to a cell cluster. The stacked barplot shows the fraction of cells from each stage within each cluster. Cell clusters K59 and K61 had high levels of the goblet module L19 containing MUC5AC. **(B)** Cell clusters K59 and K61 predominantly contained cells from 4 tumors, 2 of which were pathologically classified as IMA. Sinjab-P2, Bischoff_p030t, and Bischoff−p033t also contained other clusters of cells that did not have high expression of the goblet module L19. **(C)** Differentially expressed modules were identified for each cluster (FDR q-value < 0.05, |Log2 Fold Change| > 0.6). Samples are in the same order as in **(B).** As no adeno-like cells were observed within Bischoff_p030t and Bischoff−p033t tumors, adeno-like cells in cluster K11 from other tumors within the same dataset were included in the analysis. Similarly, adeno-like cells in clusters K6, K11, and K15 from other tumors within the same dataset were included in the analysis for Zilionis_p3. L19 was commonly significantly higher in the mucinous clusters for each tumor. Sinjab-P2 also contained clusters expressing basal or club modules. The intermediate clusters in Bischoff_p030t and Bischoff−p033t expressed L101 containing FABP5 and L102 containing Galectin-1. Goblet cells from 3 of these tumors had higher levels of the Squamous_2 module (L15). **(D)** Modules were scored in the TracerX LUAD bulk RNA-seq dataset and compared between tumors identified as IMA and those from other subhistologies. L19 was significantly higher in mucinous tumors as well as several secretory modules such as L21, L24, L25, and L79 in addition to the basal module L16. **(E)** The secretory module L79, the basal modules L16 and L17, and the squamous modules L14 and L15 were highly correlated with the goblet module L19 within IMAs from the TracerX LUAD bulk RNA-seq dataset. **(F)** Two *in vitro* scRNA-seq datasets were reanalyzed to derive gene modules denoting different transitions including AT2 to basal and progenitor cells to either lung or gastrointestinal cell fates. These external modules were scored in the cells from the 3 tumors using GSVA. Each row shows the average GSVA score for a module in each cluster. **(G)** The distributions of GSVA scores for 2 modules are shown for cells from 3 tumors. AUC was used to determine which modules could accurately distinguish the cell clusters from one another. The basal module derived from Kathiriya *et al* had higher AUCs between intermediate cell populations and the adeno-like or goblet-like cells in 3 tumors (top) compared to the gastric module from Hurley *et al*. (bottom). **(H)** These results suggest a novel model of IMA plasticity that includes transitions through squamous-like or basal-like intermediary cell populations before reaching a mucinous cell fate.

Previous studies have demonstrated gastrointestinal-like states in mucinous LUADs based on expression of lineage transcription factors such as HNF4A and CDX2 as well as mucinous markers such as TFF1, TFF2, and MUC2^39,44,45^. Concordantly, the mucinous clusters in all four patients had higher levels of HNF4A, TFF1, and TFF2 compared to their alveolar counterparts (**Supplementary Figure 13**). In contrast, the intermediate cells in each tumor generally lacked expression of gastrointestinal markers but displayed different combinations of squamous and basal markers. The intermediate cells from Sinjab_P2 expressed the basal marker KRT5 together with the canonical squamous markers TP63 and SOX2 and are referred to as basal-like, whereas the intermediate cells from Bischoff_p030t and Bischoff_p033t lacked TP63 and SOX2 but expressed other squamous differentiation markers (DSG3, KRT6A, and KRT6B) and are referred to as squamous-like. To further validate the presence of squamous or basal programs and lack of gastrointestinal gene programs in intermediate and mucinous cell populations, we re-analyzed two *in vitro* scRNA-seq datasets to derive gene modules of different transitions including 1) basal cells transdifferentiated from AT2 cells^46^, 2) gastrointestinal cells differentiated from endodermal progenitors^47^, and 3) alveolar cells derived from lung progenitors^47^ (**Supplementary Figures 14-15**, **Supplementary Tables 14-15**). We then assessed the ability of these modules to distinguish alveolar, intermediate, and mucinous-like clusters within each tumor using AUC (**Figure 4F; Supplementary Table 16**). As expected, alveolar cells could be distinguished from other clusters using the alveolar modules derived from either *in vitro* dataset (AUC > 0.80 in Sinjab_p2 and Bischoff_p30). Similarly, the mucinous cells could be distinguished from other clusters with the non-lung endoderm modules from Hurley et al (AUC > 0.80 in Sinjab_p2 and Bischoff_p33) or with the alveolar-to-basal-intermediate 2 (ABI2) module from Kathiriya *et al* (AUC > 0.9 in Sinjab_p2 and Bischoff_p33). Most importantly, the intermediate cells from Sinjab_p2 and Bischoff_p33 were best distinguished from the other clusters by higher levels of the basal module from Kathiriya *et al* (AUC ≥ 0.80; **Figure 4G**). In Bischoff_p30, the intermediate cells were best separated by the ABI2 module (AUC = 0.72) followed by the basal module (AUC = 0.68) from Kathiriya *et al*. In contrast, higher levels of the intestinal/gastric, mucinous, or non-lung endoderm modules from Hurley *et al* could not separate intermediate cells in any sample (AUC ≤ 0.46). Together, these results indicate that basal or squamous-like cell states are possible intermediate routes for formation of mucinous cells in IMA (**Figure 4H**). Furthermore, the identification of mucinous-like cells in 2 LUADs not classified as IMA suggests that this phenomenon is more widespread than previously captured by histopathology.

### Heterogeneity of squamous modules within LUSC

Several lineage, detoxification, and signaling modules were enriched in LUSCs compared to LUADs (**Figure 2D**). Modules significantly higher in the LUSC cohort included those with squamous markers such as L14 (SOX2) and L15 (TP63). The major modules containing basal markers and keratins included L16:Basal_1 (KRT5, KRT17), L17:Basal_2 (KRT6A, KRT6B, KRT14, KRT16, and KRT1), and L18:Basal_3 (KRT13). L18 was expressed in fewer cells than L16 or L17. Modules enriched for genes involved in detoxification and xenobiotic metabolism (L32-L37) or ferritin processing (L76, L77) were also higher in the LUSC cohort although several of these modules were still broadly detected at lower levels in LUADs. Different sets of ECM (L56, L57) and glycolysis modules (L71) were significantly higher in LUSC compared to LUAD while a module for keratinization (L31) was specific to LUSC. Many of these modules also displayed heterogeneity within the LUSC cohort (**Figure 5A**). Specifically, a subset of cells in clusters K53, K55, K60, and K65 did not contain high levels of the squamous or basal modules (**Figure 5B)**. LUSC cells were classified as “low”, “intermediate”, or “high” based on the density of module L14 (**Figure 5C**). Four LUSC tumors had greater than 10% of squamous-low cells including tumor Wu_P18 with 23.8% of squamous-low cells (**Figure 5D**). The module with the strongest up-regulation in squamous-low cells was L82:Fibrinogen (**Supplementary Table 17)**. In the two tumors with the most squamous-low cells, inferCNV showed that squamous-low and squamous-high cells shared the same copy number profile, consistent with a common clonal origin (**Supplementary Figure 16D**). Neither tumor showed the 3q gain characteristic of LUSC. To validate these findings, we examined the bulk RNA-seq data from TCGA or CPTAC and identified tumors with low levels of squamous modules L14 and basal module L16 (**Figure 5E**, **Supplementary Figure 16**). Module L82 expression was significantly higher in the squamous-low tumors in both cohorts (**Supplementary Tables 18-19**). Likewise, LUSCs classified as “fibrinogen-high” were enriched in squamous-low tumors in both TCGA (Fisher’s exact test p = 2.41 × 10^-9^) and CPTAC datasets (Fisher’s exact test p = 0.03, **Figure 5F**), confirming the presence of this novel subgroup across multiple datasets.

**Figure 5.**
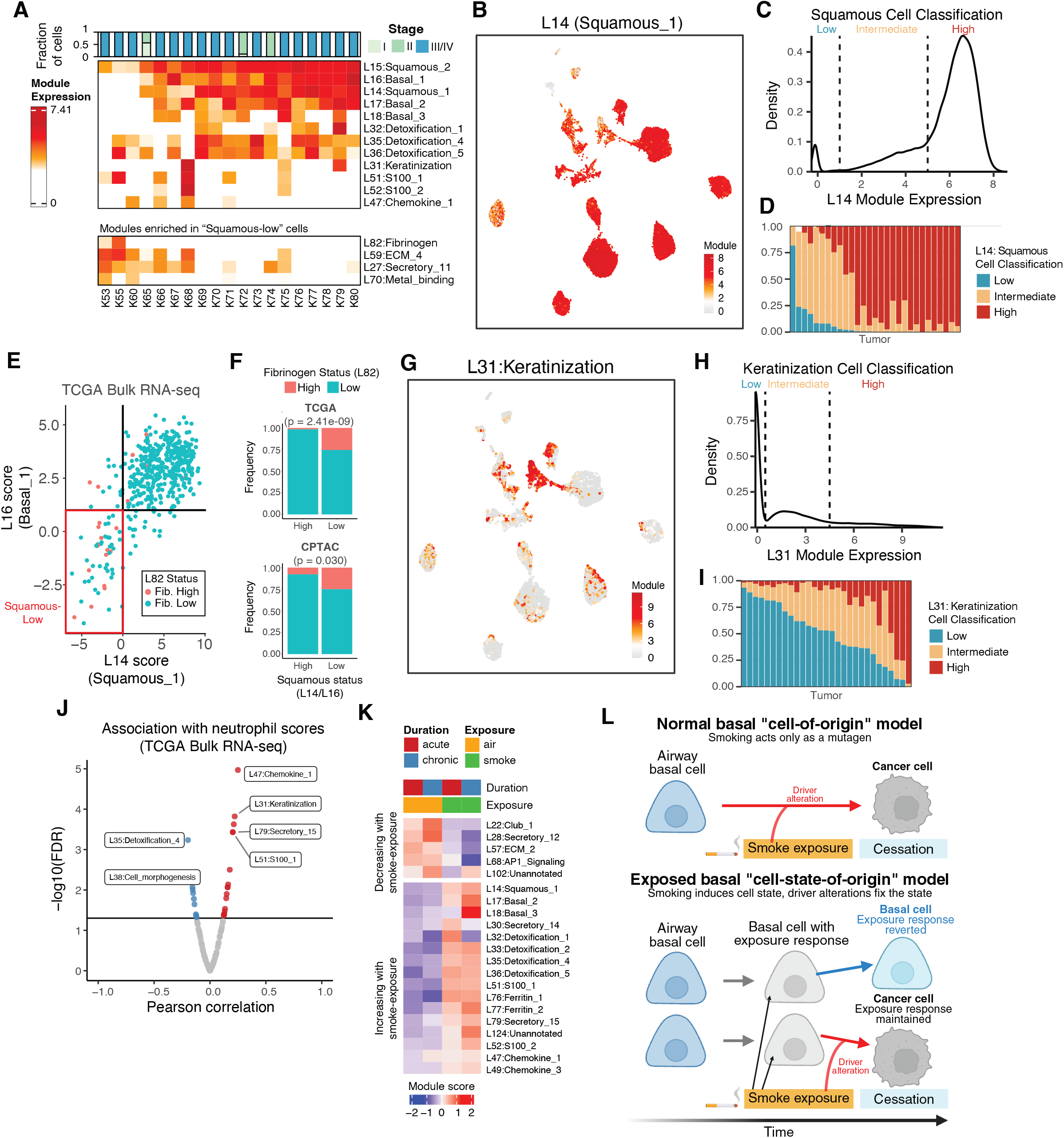
Examining heterogeneity in LUSCs elucidates subtypes and the putative cell-state-of-origin. **(A)** Each row in the heatmap corresponds to a module and each column corresponds to a cell cluster. The stacked barplot shows the fraction of cells with each stage within each cluster. Modules with significantly higher expression in the LUSCs compared to LUADs are shown. **(B)** The expression of the L14:Squamous_1 module is shown across cells from 29 LUSCs. (**C**) Cells from LUSCs were classified as squamous-low, - intermediate, or -high using the density of module L14. **(D)** Each bar shows the fraction of squamous-low, - intermediate, or -high cells in an individual tumor. The majority of LUSCs contained predominantly squamous-intermediate or -high cells. 4 LUSCs contained at least 10% of squamous-low cells. **(E)** Squamous-low LUSCs from TCGA were identified based on lower levels of L14:Squamous_1 and L16:Basal_1 in bulk RNA-seq data. **(F)** Fibrinogen-high tumors were significantly enriched in squamous-low tumors from TCGA and CPTAC LUSC cohorts validating the presence of this novel subtype. **(G)** The expression of the L31:Keratinization module is shown across cells from 29 LUSCs. **(H)** Cells from LUSCs were classified as keratinization-low, -intermediate, or -high using the density of module L31. **(I)** Each bar shows the fraction of keratinization-low, -intermediate, or -high cells in an individual tumor. The detection of cells with intermediate or high levels ranged from 6.5% to 99.8%. **(J)** The correlation between module and immune scores was assessed in TCGA LUSC bulk RNA-seq data. Module L47:Chemokine_1 containing the chemoattractant gene CXCL8 was highly correlated with neutrophil scores suggesting this module may contribute to neutrophil infiltration. **(K)** Modules were scored in a scRNA-seq dataset containing airway epithelial cells differentiated at an air-liquid interface (ALI) and exposed to air or cigarette smoke. Many of the LUSC-associated modules were significantly higher in basal cells exposed to smoke compared to unexposed basal cells. **(L)** Two models of LUSC development. Top: in the prevailing cell-of-origin model, smoke acts only as a mutagen (red curved arrow), producing driver alterations (red helix) in airway basal cells, and LUSC-associated programs (gray) emerge after transformation. Bottom: in the extended cell-state-of-origin model, smoke first induces exposure-response programs in basal cells (black arrows; gray fill) and also produces driver alterations in these cells (red curved arrow). Driver alterations fix the programs so that they are maintained in cancer cells after smoking cessation, whereas in exposed cells without such alterations the programs revert after cessation (blue arrow). Shaded bars indicate periods of smoke exposure and cessation.

The 2021 WHO Classification of Lung Tumors defines LUSC subtypes as keratinizing, non-keratinizing, or basaloid^48^. L31:Keratinization contained small proline-rich genes (SPRR1B, SPRR2D, and SPRR2A) and was enriched for “keratinization” and “Formation of the cornified envelope” (**Figure 5G**), processes that normally occur in the terminal differentiation of epidermal keratinocytes^49^. Each cell was classified as having low, intermediate, or high levels of L31 (**Figure 5H**). The detection of cells with intermediate or high levels ranged from 6.5% to 99.8%, suggesting that this molecular process is activated to varying degrees across most LUSCs (**Figure 5I**). Several signaling modules displayed heterogeneity across LUSC including those containing chemokine and S100 proteins. Neutrophils are among the most abundant immune cell types enriched in LUSC and associated with poor prognosis^50,51^. CXCL8 is a potent chemokine known to attract neutrophils^52^. L47:Chemokine_1 contained CXCL1, CXCL3, and CXCL8. Both L47 and L31 were significantly correlated with neutrophil scores when projected in the TCGA bulk RNA-seq data^53^ (**Figure 5J**) suggesting that chemokine signaling accompanying keratinization may contribute to neutrophil infiltration.

To examine whether LUSC-associated modules can be induced by smoke exposure in the absence of somatic alterations, we quantified these modules in an in vitro scRNA-seq dataset of airway epithelial cells differentiated at an air–liquid interface and exposed to cigarette smoke or air^54^ (**Supplementary Figure 17, Supplementary Table 20**). Many LUSC-associated lineage, signaling, and detoxification modules were significantly higher in smoke-exposed than air-exposed basal cells (**Figure 5K**). Because exposure was experimentally controlled in this dataset, these results indicate that smoke is sufficient to induce these programs in non-malignant basal cells. The strong epidemiological association between smoking and LUSC, together with the abundance of genotoxic carcinogens in cigarette smoke, has led to the prevailing model in which smoke acts as a mutagen, producing somatic driver alterations in airway basal cells, and LUSC-associated cell states emerge after transformation^55,56^ (**Figure 5L, top**). The presence of many of these states in non-malignant smoke-exposed basal cells suggests an extended model in which smoke first induces these programs (e.g., KRT6/13, detoxification) in basal cells, and somatic alterations arising in these cells subsequently render the programs cell-autonomous, allowing them to persist after smoking cessation (**Figure 5L, bottom**). In this extended model, the smoke-exposed basal cell, rather than the unexposed basal cell, is the more proximal “cell-state-of-origin” for LUSC.

### Combinations of genetic alterations associated with gene modules

We next sought to determine which modules could potentially be driven by somatic alterations including nonsynonymous mutations and focal CNAs. As mutational data was not available for most scRNA-seq samples, we scored each module in the LUAD or LUSC TCGA bulk RNA-seq datasets and used CaDrA to identify the best combination of known driver alterations that were associated with each module^57^ (**Methods; Supplementary Table 21**). Modules L32-L37 were enriched for genes involved in detoxification, oxidative stress, and xenobiotic metabolism (**Figure 6A**). L33, L35, and L36 were associated with alterations in the Nrf2 pathway including *NFE2L2* and *KEAP1* (**Figure 6B**). As a protein product of the *NFE2L2* locus, Nrf2 is a transcription factor that induces the expression of antioxidant genes while Keap1 is the substrate adaptor of a CUL3-based E3 ubiquitin ligase complex that targets Nrf2 for degradation^58^. *KEAP1*, *NFE2L2*, and *CUL3* alterations were also associated with ferritin processing modules L76 and L77 in LUSC (**Figure 6C**). These modules were also strongly associated with *KEAP1* alterations in LUAD. Cell cycle and proliferation modules such as L39 and L41 were another biological class which could be explained by combinations of genetic alterations (**Figure 6D**). In LUAD, amplifications of *CCND3* and *CCNE1* were associated with these modules along with alterations in other genes such as *MET* amplifications, *SMARCA4* mutations, *FGFR1* amplifications, and *KAT6A* amplifications (**Figure 6E**). In LUSC, module L41 was associated with alterations in the cell cycle regulator *RB1* (**Figure 6F**). This module along with other cell cycle related modules L43 and L46 were enriched for E2F1, E2F4, and E2F6 transcription factor (TF) binding sites (**Supplementary Table 22**). *RB1* is a tumor suppressor which normally interacts with E2F TFs to inhibit expression of cell cycle genes^59^. In LUAD patients, the *STK11* mutations were associated with several modules including L66, L25, L126, L94, and L82. L66 is a stress response module enriched for genes involved in the unfolded protein response (UPR) which has been shown to be a vulnerability of *STK11/LKB1* knockout mouse models^60^. Scissor was also used to integrate bulk TCGA LUAD RNA-seq data with malignant cells from the atlas using *KEAP1* mutation status as the phenotype. Modules L33, L35, and L36 were significantly higher in Scissor+ than Scissor− cells (p < 0.001, Wilcoxon rank-sum test; **Supplementary Figure 18**). Overall, these results demonstrate the diversity of genetic alterations that are candidates for driving the detoxification, cell cycle, and stress response gene expression programs active in lung cancer cells. Interestingly, a general lack of associations was observed for other categories of modules including the majority of lineage and signaling related pathways suggesting that other types of mechanisms may be important for driving these changes (e.g. epigenetic).

**Figure 6.**
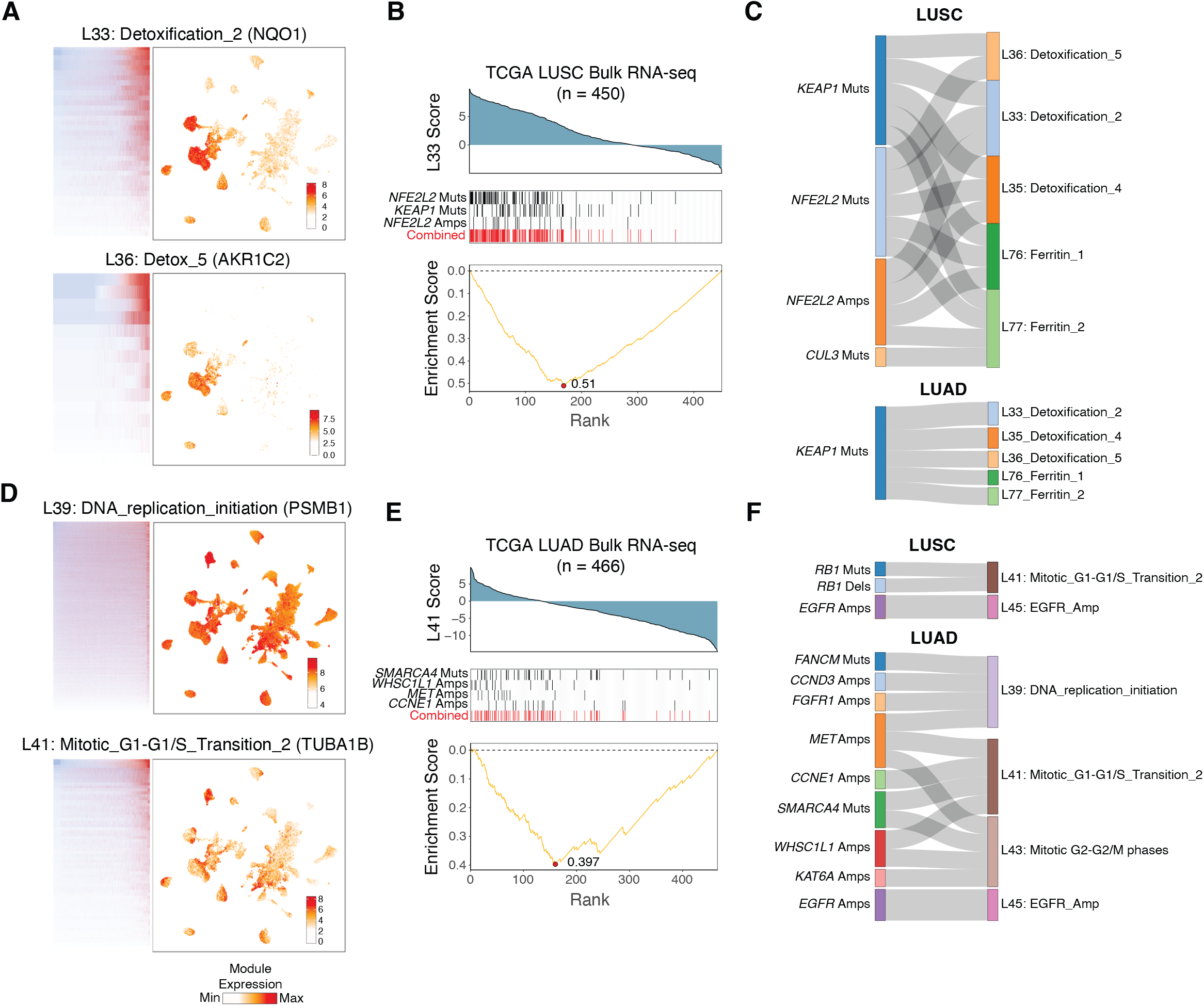
Genetic drivers of module expression. **(A)** 6 modules were enriched in detoxification genes. Co-expression heatmaps and UMAPs with corrected module expression are shown for modules L33 and L36. Cells in the co-expression heatmaps are ordered from those with the lowest expression of the module to the highest. **(B)** Modules were scored in TCGA bulk LUSC or LUAD RNA-seq data and CaDrA was used to identify combinations of mutations and focal CNAs enriched in higher module expression. An example CaDrA result is shown for module L33 in LUSC. Top: Each vertical bar indicates the expression score for that module in a tumor and tumors are rank ordered from highest to lowest score. Middle: CaDrA selected combinations of somatic alterations that are enriched among tumors with higher module expression. Vertical black bars indicate that the specific alteration is present in that tumor. The red vertical bars indicate that any of the selected features are present in that tumor. Bottom: Running enrichment score for KS test. The point farthest from zero indicates the maximal enrichment. **(C)** The Sankey diagram summarizes several CaDrA results and shows that several alterations in the Nrf2 pathway are associated with detoxification and ferritin processing modules. **(D)** 9 modules were enriched in proliferation and cell cycle related genes including modules L39 and L41 which are shown as examples. **(E)** CaDrA result demonstrating alterations associated with module L41 in LUAD including *CCNE1* amplifications. **(F)** Different sets of alterations were associated with different cell cycle modules including *RB1* mutations and deletions in LUSC or *CCNE1* amplifications in LUAD for L41.

### Benchmarking the ability of different factorization tools to perform GPAS

Co-expression module discovery followed by association testing with clinical variables is widely used in cancer; however, the accuracy with which different factorization methods recover gene modules and patient-level effects has not, to our knowledge, been systematically evaluated in cancer cells, where cells are nested within tumors and studies and much of the variation is tumor-specific. Because the true modules and effect sizes are unknown in real data, we generated a simulated dataset where each cell was composed of 175 synthetic modules with the same technical and biological effects observed in our NSCLC atlas including study, sample, cluster, histology, and stage (**Methods**). The UMAP generated using these module scores demonstrates that many tumors form their own cluster, global variation can be observed between studies or different tumor histologies, and more subtle variation can be observed between low and high stage patients within studies (**Supplementary Figure 19A**). Rather than using the true expression counts for each gene, counts for 25 synthetic genes were generated for each module by sampling from a Dirichlet distribution (25 genes * 175 modules = 4,375 genes total). We applied two module detection tools, GeneNMF and hdWGCNA^83^, as well as RcppML, which contains a fast implementation of NMF. For standard NMF, three different approaches were used to assign genes to modules (**Methods**). We examined several benchmarking metrics including the number of genes assigned to a module, the number of modules with at least one assigned gene, Adjusted Rand Index (ARI) of the module assignments, and the root mean square error (RMSE) for estimating the recovery of the two biological coefficients, histology and stage. These metrics capture complementary properties: the number of genes assigned measures coverage, since unassigned genes cannot be tested for associations; the number of modules with assigned genes measures whether the full set of programs is recovered; ARI measures the accuracy of gene-to-module assignments relative to the ground truth and penalizes both missed and incorrect assignments; and RMSE measures how accurately patient-level effects are recovered from the module scores, which is the goal of GPAS. Celda was the only approach that clustered all genes into a module and all modules had at least one gene (**Supplementary Figure 19B**). For ARI, Celda was ranked 6th behind some implementations of NMF and hdWGCNA although the ARI was still relatively high at 0.94 (**Supplementary Figure 19C**). Of note, hdWGCNA only assigned a small fraction of genes to a module (23%) and discovered fewer modules with default parameters. Celda also had the lowest RMSE values when using LMEs to estimate coefficients for histology and stage (**Supplementary Figure 19D**). Celda performed better than GeneNMF for recovery of patient-level coefficients and number of genes assigned to a module (**Supplementary Figure 19E**). In summary, in simulated data with the hierarchical structure of a multi-study cancer cell atlas, Celda assigned all genes to modules while recovering module membership with high accuracy and patient-level effects with the lowest error among the methods tested.

### Comparison of different factorization or co-expression approaches using the NSCLC cancer cell atlas

To complement the simulation benchmarking, we compared module discovery methods directly on the NSCLC cancer cell atlas, evaluating how well each approach recovered known modules, produced coherent gene-to-module assignments, corresponded to annotated biology, and associated with clinical phenotypes. geneNMF failed in all 81 runs to recover lineage-specific modules (L5, L8–L10, L14–L17, L19, L31, L82, L84, L86–L88), xenobiotic metabolism modules L35–L36, proliferation modules L38–L39, and many housekeeping modules (nine representative runs shown in **Supplementary Figure 20A**). hdWGCNA failed in all 9 runs to recover lineage-specific modules L7–L8 and L86, cellular process modules L62, L74, L76, and L103–L104, proliferation modules L39 and L44–L46, and many housekeeping modules (**Supplementary Figure 20A**; **Supplementary Table 23**). Of the 45 histology-associated and 35 stage-associated modules, 22 and 15 were not recovered by geneNMF in any run, and 8 and 3 were not recovered by hdWGCNA. Conversely, Celda recovered a median of 97% of geneNMF metaprograms (range 40–100% across 81 runs) and 100% of hdWGCNA modules, indicating that the Celda module set largely subsumes those of the other methods (**Supplementary Figure 20B**). Standard NMF implemented in RcppML showed greater overlap with Celda modules when genes were assigned to factors by the “Maximum” or “Correlation” methods than by the top-unique-genes method. Across all runs, RcppML recovered 140 of 141 Celda modules, including 43 modules recovered by all three methods and 35 modules recovered by neither geneNMF nor hdWGCNA (**Supplementary Figure 21A**). The run with 200 factors and correlation-based gene assignment (RcppML_k200_Corr) captured the largest share of Celda modules and was used for all subsequent comparisons. However, the correspondence between RcppML factors and Celda modules was not one-to-one: 54 of the 200 factors did not significantly overlap any Celda module, and over half (108 of 200) overlapped two or more distinct Celda modules, indicating that individual factors were less specific than Celda modules (**Supplementary Figure 21B**). Consistent with this, the average correlation between each gene’s expression and the activity of its assigned module or factor was significantly higher for Celda modules than for RcppML_k200_Corr factors (Wilcoxon rank-sum p = 0.0008; **Supplementary Figure 21C**). For example, factor NMF_80 was active in a single squamous-enriched cluster, whereas its top-loading genes (SPP1, AKR1C2, AKR1C3) were expressed across several additional clusters that were captured by the corresponding Celda module L36 (**Supplementary Figure 21D**). The same discordance was observed between factor NMF_99 and Celda module L14, which share the squamous lineage marker SOX2 (**Supplementary Figure 21E,F**). Reactome over-representation analysis identified significant enrichment (q < 0.05) for 122 of 175 Celda modules (70%) compared with 116 of 200 NMF factors (58%), indicating that a higher proportion of Celda modules corresponded to annotated biological processes (Fisher’s exact test p = 0.024). Finally, we fit the same LME used for Celda modules to the RcppML_k200_Corr factor activities and compared the strength of clinical associations. Celda modules had a significantly higher percentage of variance explained by the histology and stage fixed effects than NMF factors (Wilcoxon rank-sum p = 1.08 × 10⁻; **Supplementary Figure 21G**), and the association significance (−log10 FDR) for histology and stage showed the same pattern (Wilcoxon rank-sum p = 8.83 × 10^-10^ and p = 5.8 × 10^-7^, respectively; **Supplementary Figure 21G**). Together, these analyses show that no competing method matched Celda across the properties most relevant to this study. geneNMF and hdWGCNA failed to recover many lineage-specific, proliferation, and housekeeping modules, and while standard NMF recovered most Celda modules, it did so through broad, less coherent factors that each aligned with multiple Celda modules. Celda modules were also more often enriched for annotated biological processes and more strongly associated with clinical phenotypes.

### Sensitivity analysis to compare module detection between “pooled”, “non-pooled”, and “down-sampled cell” multi-study designs

Our primary analysis used a “pooled” design in which Celda was run on cancer cells from all studies and LMEs were subsequently used to account for inter-study differences. An alternative “non-pooled” design runs Celda on each study separately and then reconciles recurrent modules across studies. To evaluate the sensitivity of our results to this choice, we ran Celda independently on each of the 12 studies and compared the resulting study-specific modules to the 175 modules from the pooled analysis (Methods). Rather than merging study-specific modules into a consensus set, which would require arbitrary similarity thresholds, we directly assessed how often each pooled module was recovered within individual studies. Of the 141 pooled modules with at least five genes, 138 significantly overlapped a module in at least one study-specific run (Fisher’s combined p < 1 × 10^-5^; **Supplementary Figure 22A**). In the reverse direction, an average of 81% of study-specific modules (range: 66–87%) significantly overlapped a pooled module, indicating that the pooled analysis captured most modules identifiable within any single study.

We next asked whether the histology and stage associations identified in the pooled analysis were preserved in the study-specific runs. For each pooled module associated with histology, we identified its best-matching module in each of the four studies containing both LUAD and LUSC tumors and tested that module for association with histology within the study. All 45 histology-associated pooled modules were concordantly associated with histology (nominal p < 0.05) in at least one study (**Supplementary Figure 22B**). For stage, all 35 stage-associated pooled modules were concordantly associated in Xing et al. (**Supplementary Figure 22C**), the study with the most LUAD tumors (14 early-stage, 7 late-stage), and 30 were also concordant in at least one additional study. A small number of modules showed opposite-direction associations in individual studies with few tumors, illustrating why the statistical power of a multi-study meta-analysis is needed to establish associations robustly. Overall, these results indicate that the majority of associations identified in the pooled analysis are reproducible within individual studies.

Lastly, because the number of cancer cells per tumor ranged from 51 to 7,806, we assessed whether tumors with many cells disproportionately influenced module discovery. Within Wu et al., the largest study, 99 of 100 modules learned from all cells were recovered after capping each tumor at 500 cells (**Supplementary Figure 22D**). We then capped each tumor in every study at 500 cells and reran Celda separately within each study, so that module discovery was insulated from both cross-study pooling and cell-number imbalance. Under these conditions, 136 of 141 original modules were recovered in at least one study (**Supplementary Figure 22E**), nearly identical to the result without downsampling. Together, these results indicate that cell-number imbalance across tumors has little effect on module discovery.

## DISCUSSION

Previous scRNA-seq studies have only been able to investigate the transcriptome of cancer cells from a limited number of NSCLCs. In this study, we combined scRNA-seq data for 127 NSCLCs from 12 studies and determined the combination of co-expression modules that could reconstruct each tumor. Differences in cell dissociation procedures and single cell platforms can influence gene expression and produce batch effects. Standard single-cell integration approaches assume that the same cell types or states are observed across studies and can thus be aligned in a reduced dimensional space where clustering and 2D visualization are performed. This may be reasonable for shared cell types such as immune cells in the tumor microenvironment. However, cancer cells present a unique challenge in that each tumor may have a unique combination of genetic alterations, epigenetic changes, and interactions with other cells in the TME that impact the cancer cell transcriptome. The majority of integration methods do not have capabilities for dissecting variation from different technical and biological sources. One major advantage of a module-based approach is that the overall sparsity is greatly reduced compared to individual genes due to the presence of multiple genes in most modules. This allowed us to utilize LMEs to quantify the variability in each module attributed to differences between studies and samples. While some highly expressed housekeeping modules were largely driven by study-to-study effects, the majority of modules with biologically diverse functions were more strongly associated with biological and clinical variables such as histology or stage. We also observed that Histology and Stage had large effect sizes on more modules compared to Study. These observations demonstrate that cross-study analyses of heterogeneous tumors are possible because the magnitude of biological signal can outweigh the technical noise. We refer to our approach as a Gene Program Association Study (GPAS) as it is analogous to a Genome-Wide Association Study (GWAS) which seeks to understand how inherited genetic variation is associated with traits or disease risk.

In general, dedifferentiation and lineage plasticity have been associated with worse clinical outcomes, metastasis, and resistance to treatment^17,61,62^. In this atlas, several types of lineage-related heterogeneity could be observed. First, complete dedifferentiation was observed in some late-stage LUADs that did not have detectable levels of modules containing specific AT2 markers and reduced or absent expression of modules containing general broader alveolar markers such as NKX2-1, NAPSA, and RNASE1. Additional stochastic lineage infidelity was also evident in cells from other LUADs which displayed different combinations of detected AT2 and AT1 markers. No module exclusively contained human AT1 markers. Instead, markers such as AGER and RTKN2 were spread across multiple modules, and examination of the individual markers confirmed that AGER and RTKN2 were low or absent from most cancer cell clusters. Furthermore, cells with a coordinated AT1 program in the absence of AT2 expression were rare in invasive LUAD, suggesting that full AT2-to-AT1 transitions are uncommon in primary untreated tumors. The notable exception was a single GGO lesion in which most malignant cells were AT1-like. Because AT1 cells have been proposed as a cell of origin for lepidic LUAD in mouse models^30,63^, this lesion may represent a tumor arising from AT1 cells rather than one that has transdifferentiated from an AT2 state, although we cannot distinguish these possibilities from expression data alone. These observations do suggest that once tumorigenesis has begun and an AT2 state has been obtained, the expression of individual AT2 and AT1 genes can be independently modulated resulting in disordered expression across tumors. Lastly, lineage plasticity or transdifferentiation is the ability of cells to transition from one committed developmental pathway to another^62^. We observed lineage plasticity in mucinous adenocarcinomas where alveolar markers were completely lost and programs for mucinous/goblet cell types were gained in a subset of cells. Surprisingly, other subpopulations were identified in the IMAs including basal and club cells in one tumor and squamous-like cells in two other tumors. This observation suggests that basal or squamous-like intermediates may precede the formation of mucinous cells. This finding is supported by recent *in vitro* experiments demonstrating that basal and club cells can be transdifferentiated from AT2 cells with TGFβ1 and anti-bone morphogenetic protein signaling (BMP) likely derived from fibroblasts^46^.

Lineage loss was also observed in LUSC, where a subset of cancer cells in several tumors lacked squamous and basal modules and instead expressed fibrinogen (L82) and ECM (L59) genes. Several interpretations are possible. In Wu_P34, the squamous-low cells shared the copy number profile of the squamous cells, indicating that they arose within the same clone and represent a dedifferentiated, EMT-like state of the squamous tumor, analogous to the loss of alveolar identity in late-stage LUAD. The presence of the neuroendocrine gene CALCA in L82 raises the additional possibility of partial neuroendocrine differentiation, as has been described in lineage transitions between squamous and neuroendocrine states in other contexts. Alternatively, these tumors could represent a metastasis from an extrapulmonary primary that does not depend on 3q gain such as cutaneous basal cell carcinoma metastasizing to the lung^9,64^. The reproducible detection of fibrinogen-high, squamous-low tumors in TCGA and CPTAC indicates that this subgroup is not restricted to the single-cell cohort, but distinguishing these possibilities will require matched pathology and genomic profiling.

The observation that many LUSC-associated modules are induced in non-malignant basal cells by cigarette smoke suggests an extension of the cell-of-origin model for LUSC, in which smoke acts not only as a mutagen but also as an inducer of the cell state in which driver alterations subsequently arise. Several lines of evidence indicate that somatic alterations alone are not sufficient to produce LUSC in the human airway. Driver mutations are abundant in the histologically normal bronchial epithelium of smokers^56^, preinvasive squamous lesions carrying driver mutations frequently regress^65^, and although never-smokers develop lung cancer, they rarely develop LUSC^66^. In the extended model, the exposure-induced state contributes to the fitness of basal cells under continued smoke exposure, and alterations that render the same programs cell-autonomous, such as *KEAP1* and *NFE2L2* mutations for the NRF2-dependent detoxification modules, are favored because they maintain an adaptive phenotype after exposure ends. This is analogous to genetic assimilation, in which an environmentally induced phenotype becomes genetically encoded through selection^67^. Because many smoke-induced genes revert rapidly after cessation in normal airway epithelium^68^, this model predicts that their persistence in LUSC from former smokers depends on genetic or epigenetic fixation, a prediction that can be tested as matched exposure histories and genomic profiles become available for single-cell cohorts.

Using the GPAS framework, we were able to identify gene modules associated with late-stage disease. Stage III tumors have regional (mediastinal or contralateral) nodal involvement or locally invasive disease, while Stage IV tumors have intrathoracic or distant metastatic disease. As our cohort only contained the primary tumor samples, we were not able to ascertain whether these modules were also enriched in metastatic cells or if metastasis-specific modules also exist. Generating cohorts of primary and metastatic samples will be necessary to determine if the late-stage associated modules are indeed enriched in metastatic cells. Future work will be needed in a clinical setting to determine if the gene modules associated with late-stage disease are predictive of worse outcome when active in early-stage tumors or can predict responses to specific therapies. Ultimately, characterizing the combinations of gene programs that contribute to tumor aggressiveness in NSCLCs will aid in the development of novel biomarkers for patient prognostication and reveal novel avenues for therapeutic development.

## MATERIALS and METHODS

### Single cell RNA-seq dataset preprocessing

The original gene count matrices and cell type annotations were downloaded from the GEO repository or transferred from corresponding authors upon request. Quality control analysis was performed using SCTK-QC pipeline^69^. Cells were excluded if they had fewer than 500 genes detected or contained more than 20% of counts from mitochondrial genes. Genes that were detected in less than 10 cells were excluded from downstream analysis. For studies that did not publish or deposit their final cell type annotations, we performed initial annotation analysis with SingleR package using the Human Primary Cell Atlas^70^ as the reference to identify epithelial cells. Clustering with the Seurat workflow^71^ was also performed and expression of EPCAM was examined to distinguish epithelial populations from other immune and stromal populations.

### Identification of cancer cells with copy number variations (CNVs)

To obtain CNV estimates for each tumor, we utilized the *inferCNV* package^72^ with the following parameters: using ‘i3 hidden Markov model (HMM)’ model to infer CNV events and setting the “cutoff” as 0.1 to determine which genes to use in inferCNV analysis. Stromal cells and immune cells from the same tumor samples were considered as putative nonmalignant cells and used as the reference. Low-probability CNVs were filtered using the threshold of *BayesMaxPNormal* = 0.5. *analysis_mode* = ‘subcluster’ was used to identify different subclones in each tumor sample. Malignant cells were detected per sample and filtered using the following procedure: 1) The average CNV signal for each cell was calculated as the mean square of all CNV estimates across all gene regions; 2) The top 5% of cells with the highest CNV signals within each tumor were designated as reference malignant cells, and their average CNV signal was treated as the reference CNV signal for malignant cells; 3) For each cell within the tumor sample, the Pearson correlation coefficient between its CNV signal and the reference malignant CNV signal was computed; 4) Cells with a correlation coefficient greater than 0.2 were classified as malignant and included in the final analysis. Finally, tumors with less than 50 malignant cells detected were excluded from downstream analysis.

### Identification of gene modules with Celda

The gene count matrices containing all malignant cells from all studies were concatenated. The top 10,000 genes with highest average expression were obtained for each tumor. The union of these genes (n=13,979) were used for module detection. The Celda package was used to simultaneously cluster genes into co-expression modules and cells into clusters. To choose a reasonable solution for the number of modules L and cell clusters K, the function *recursiveSplitModule()* was used to examine the range of L from 80 to 200. The function *recursiveSplitCell()* was used to examine the range of K from 70 to 90. After initial application of Celda, we observed that some cell clusters were enriched for modules containing genes for ciliated cells or B cells. These cell clusters and genes in those modules were removed from the dataset and the Celda analysis was repeated. The rate of perplexity change (RPC) was used to select an initial solution of L = 165 and K = 80 as previously described^27^. 10 gene modules were split into two modules with the *splitModule()* function resulting in 175 gene modules and 80 cell clusters for downstream analysis. The counts of each gene module in each cell were retrieved using the *factorizeMatrix()* function. The log2 normalized expression for each gene module was calculated by dividing the module count by the total number of counts in that cell, multiplying by 10,000, and applying a log2 transformation after adding a pseudocount of 1. For the lineage infidelity analysis, gene modules that had at least 0.5% of the total counts in a cell were considered detected in that cell. Enrichment of each module for Reactome^73^ functional categories and for chromosomal cytobands was assessed with Enrichr^74^.

### Using linear mixed-effect (LME) modeling to dissect biological and technical variability

We applied a linear mixed-effects (LME) regression model to analyze the variability in module expression across cells. The model treated each cell as a repeated measure by including fully nested random effects for Study, Sample, and Cluster. Additionally, fixed effects for histology and tumor stage were incorporated to account for biological heterogeneity stemming from histological and clinical phenotypes. The LME model was formulated as follows:

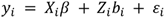

Where:

- 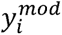 : Module expression for cell i.
- *X_i_*: A design matrix of fixed effects (Histology + Stage).
- *β*: A vector of fixed-effect coefficients.
- *Z_i_*: A design matrix of random effects for cell i (Study/Sample/Cluster).
- *b_i_*: Random effects for cell i.
- *ϵ_i_*: Residual error term for cell i.

The variance explained by each random effect was calculated using the *get_variance()* function from the *insight* package in R. The significance of fixed effects was determined using a Type III ANOVA test performed with the *Anova()* function from the *car* package in R. P-values were corrected for multiple hypothesis testing using the false discovery rate (FDR). A similar LME model was applied to LUADs to identify modules associated with stage with the only difference being the omission of the histology term. Tumors with a histological diagnosis of NSCLC-NOS (n = 7, all from Chen et al.) were excluded from the LME models, as the Histology fixed effect was specified as a contrast between LUAD and LUSC.

### Correcting module expression for study-level effects

The full LME model was used to remove the effects of the Study variable on each module while preserving other biological effects. The corrected module score was calculated using the following formula:

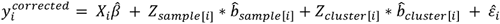

Where:

- *X_i_* and *β*: Design matrix and coefficients for fixed effects fitted by the LME model for cell i.
- *Z_sample[i]_* and *b̂*_*sample[i]*_: Design matrix and random effect estimates corresponding to Sample of cell i.
- *Z_cluster[i]_* and *b̂*_*cluster[i]*_: Design matrix and random effect estimates corresponding to Cluster of cell i.
- *ϵ̂*_*i*_ : Estimated residual error term for cell i.

Corrected log2-normalized module expression was used in all downstream analyses unless otherwise noted.

### Creating 2D embeddings with UMAP

The corrected module expression for gene modules that had less than 25% of variability explained by the Study random effect was used to generate a 2D embedding with the UMAP algorithm. The UMAP was created with the *getUMAP()* function from the SCTK package using the first 25 principal components (PCs) from a principal component analysis (PCA). To generate a LUSC-specific UMAP, cells were limited to those originating from LUSC samples and that were grouped into clusters containing more than 60% of cells from LUSC samples. The UMAP was generated using the same procedure for the full dataset.

### Scoring modules in bulk RNA-seq datasets

RSEM-normalized gene expression values, DNA copy number (GISTIC2), somatic mutation, tumor purity score (ABSOLUTE), and patient metadata of TCGA-LUAD cohort were obtained from FireBrowse (version 28 January 2016). The RSEM-normalized expression matrix was log2 transformed after adding a pseudocount of 1 for downstream analysis. For each gene module, principal component analysis (PCA) was performed on the z-scored log2 expression values of its gene members using the *prcomp()* function in R, similar to the eigengene calculation in WGCNA^75^. The first principal component (PC1) was used as the module score. Since the direction of the principal component is arbitrary, the direction for each module score was reversed by multiplying the scores by -1 if the scores were negatively correlated with the average expression of the genes in the module. To associate module scores with subhistology in the TRACERx cohort, we used an LME model that included subhistology as a fixed effect and Patient ID as a random effect.

### Defining gene modules and AT1/AT2 reference cells in normal lung alveolar cells

To characterize detection of lineage genes in normal alveolar cells from non-cancerous lung tissue, scRNA-seq data was downloaded and analyzed for 10 donors from Basil *et al*^32^ from the Gene Expression Omnibus (GSE168191). Quality control was performed with the SCTK-QC pipeline^69^. Cells with at least 500 genes and 750 counts were included while cells with greater than 10% mitochondrial counts were excluded. Cells were initially clustered with Celda with K=50 and L=150 using the top 10,000 most variable genes after excluding those with fewer than 5 counts in 5 cells. Clusters containing lung epithelial cells were identified with canonical markers including basal (KRT5), secretory (MUC5B), club (SCGB1A1), goblet (MUC5AC), ciliary (FOXJ1), transitional (SCGB3A2), AT1 (AGER), and AT2 (SFTPC) cells. These cells were reclustered using Celda with K=50 and L=125 using the decontaminated counts from decontX^76^. One module contained AGER, CAV1, and RTKN2 and was used to define AT1 cells. One module contained SCGB3A2 and was used to define alveolar transitional cells. Four modules contained one or more surfactant genes and were used to define AT2 cells. For the lineage infidelity analysis, gene modules that had at least 0.5% of the total counts in a cell were considered detected in that cell.

Two additional normal lung datasets were processed with the same general workflow to serve as independent AT1 and AT2 reference populations. For Travaglini *et al*^77^, 10x Genomics UMI counts were downloaded from Synapse (syn21041850), quality control was performed with SCTK-QC, and cells with at least 500 genes and 750 counts were retained. Cells were initially clustered with Celda (K=50, L=150) using the top 5,000 genes, lung epithelial clusters were selected with the canonical markers listed above, and these cells were reclustered with Celda (K=30, L=100) using the top 15,000 genes after excluding those with fewer than 3 counts in 3 cells. For Murthy *et al*^78^, filtered count matrices for three donors (GSM5388411–GSM5388413) were downloaded from GEO, quality control was performed with SCTK-QC, and cells with at least 500 genes, 750 counts, and less than 25% mitochondrial counts were retained. Cells were clustered with Seurat (2,000 highly variable genes, 75 principal components, Louvain algorithm), lung epithelial clusters were selected with the canonical markers, and these cells were clustered with Celda (K=25, L=100) using the top 10,000 genes after excluding those with fewer than 5 counts in 5 cells. In each dataset, cell clusters were labeled as AT1 or AT2 if they were defined by modules containing the same AT1 (AGER, CAV1, RTKN2) or AT2 (surfactant) marker genes identified in Basil et al. To assess lineage identity independently of the Celda module assignments, cancer cells and normal AT1 and AT2 reference cells were scored with AUCell using curated AT1 and AT2 marker sets, and Celda clusters were ordered by the median of the AT1 minus AT2 AUCell score. Malignant cells were classified as AT1-like if their AT1 score was at or above the 25th percentile and their AT2 score at or below the 75th percentile of the corresponding scores in normal AT1 cells, and as AT2-like if their AT2 score was at or above the 25th percentile and their AT1 score at or below the 75th percentile of the corresponding scores in normal AT2 cells; all other cells were considered partial/dedifferentiated. Because AUCell scores depend on gene set size and expression level, thresholds were defined separately for each signature from the reference cells rather than at a common value.

### Examining lineage infidelity in spatial transcriptomics data

We analyzed a publicly available spatial dataset of a human LUAD FFPE sample published by 10X Genomics (Visium HD Spatial Gene Expression Libraries, Post-Xenium, Human Lung Cancer [FFPE], Experiment 1, HD only [control]). Space Ranger v4.0.1 was used to map fastq files to the transcriptome to get the cell segmentations. Cells that passed the threshold of (a) UMI counts > 150, (b) detected features > 50, and (c) percentage of mitochondrial genes < 10 were retained for downstream analysis. inferCNV was used to identify cells with CNV events with stromal cells and immune cells as a reference group and epithelial cells were used as an observation group. Malignant cells were classified using the same CNV-based criteria as the single-cell atlas. Module scores were calculated as the sum of raw counts across genes in each module, normalized by per-cell size factor computed using all genes in the dataset, followed by log2 transformation.

### Generating basal and gastric modules using published in vitro datasets

Gene count matrices were downloaded from GEO for the Hurley *et al*^47^ and the Kathiriya *et al*^46^ datasets. The authors of Hurley *et al*.,^47^ supplied the original cell annotations as well as UMAP coordinates. A subset of cells in the Non-lung Endoderm (intestine/liver/stomach) cluster expressed mucinous genes. We split this cluster of cells into 2 distinct clusters for further analysis. Celda_CG was used to extract gene modules with K = 10 and L = 100. Modules were identified that could distinguish the non-lung endoderm, mucinous, and alveolar fates. For the Kathiriya *et al* dataset^46^, QC parameters and the preprocessing workflow from the original paper were followed. Cells were included if they had less than 6,000 counts, more than 200 features detected, and less than 10% mitochondrial reads. For batch correction, we used fastMNN with the subset of the 2000 genes with the highest variance. After batch correction, SingleR was used to filter out non-epithelial cells and cells were clustered using the Louvain algorithm on an SNN graph according to the Bioconductor OSCA best practices^79^. Celda_CG was used to extract gene modules with K = 10 and L = 100. Modules were identified that could distinguish the original AT2-like cells, alveolar-to-basal intermediate clusters 1 and 2 (ABI1 and ABI2), as well as basal cells. Modules of interest from both datasets were scored in 3 tumors with mucinous and intermediate cells using GSVA^80^. AUC was used to determine which modules could best distinguish alveolar, intermediate, mucinous, and/or club cells within each tumor. CytoTRACE2^81^ was applied to the cancer cells from each tumor containing mucinous cells, and scores were compared between the alveolar-like, intermediate, club-like, and mucinous clusters.

### Determining association of modules with smoke exposure

To determine the association of LUSC-related modules with smoke exposure, scRNA-seq data from basal cells exposed to cigarette smoke and differentiated at an air-liquid-interface (ALI) was reanalyzed from Wohnhaas *et al*^54^. Fastq files were obtained from the European Nucleotide Archive (PRJEB44878) and processed with CellRanger v6.1.2 to generate count matrices. Quality control was performed with the SCTK-QC pipeline^69^. Cells with at least 500 genes and 750 counts were included while cells with greater than 10% mitochondrial counts or greater than 25% of counts from long non-coding RNAs (lncRNAs), MALAT1 and NEAT1, were excluded. Cells were initially clustered with Seurat v4^71^ using 2,000 highly variable genes, 50 dimensions from PCA, and the Louvain clustering algorithm with resolution of 0.8. Basal and secretory cells were identified using known markers and reclustered with Celda. Clusters containing basal cells were identified using the module containing KRT5. The lung cancer cell modules were scored in this scRNA-seq dataset as follows: 1) Module counts were calculated as the sum of all gene counts contained within each module; 2) The log2 normalized expression for each gene module was calculated by dividing the module count by the total number of counts in that cell, multiplying by 10,000, and applying a log2 transformation after adding a pseudocount of 1. To assess the association between lung cancer module scores and exposure to smoke or air in basal cells, we used an LME model that included exposure (smoke or air) and duration of exposure as fixed effects, and Patient ID as a random effect. The significance of fixed effects was determined using a Type III ANOVA test performed with the Anova() function from the car package in R. P-values were corrected for multiple hypothesis testing using the false discovery rate (FDR). Lung cancer module scores that were differentially expressed following smoke exposure were identified based on FDR < 0.05 and |log2 fold change| > 0.5.

### Determining mutational drivers of modules with CaDrA

DNA copy number (GISTIC2), somatic mutation, tumor purity score (ABSOLUTE), and patient metadata of TCGA cohorts were obtained from FireBrowse (version 28 January 2016). CaDrA was run separately for LUAD (466 samples) and LUSC (450 samples). Mutations with a frequency between 4% and 60%, as well as CNVs with a frequency between 2% and 60% were included as potential features. For the CaDrA candidate search, the top 10 starting points were evaluated and a maximum size of the meta feature of 4 individual features was chosen. Meta features were scored using the “ks_pval” function and the top-scoring meta feature of each module that had an enrichment score above 0.35 and a significant FDR-corrected p-value was further investigated. Transcription factor binding site enrichment was performed using enrichR and the “ENCODE_and_ChEA_Consensus_TFs_from_ChIP-X” database with a cutoff of 0.05 on the adjusted p-values. Scissor v2.0^82^ was also used to identify malignant cells associated with mutation status. Samples were labeled as mutant or wild-type for each driver gene. This binary phenotype was supplied to Scissor together with the log-normalized expression matrix of LUAD malignant cells from the atlas, using family = “binomial” and α = 0.5. Significance of the resulting cell selection was confirmed with Scissor’s reliability significance test. This procedure was repeated for the TCGA LUSC cohort and the LUSC malignant cells.

### Benchmarking the ability of different tools to perform GPAS

We performed a comprehensive benchmarking comparison of various factorization and clustering methods to identify associations of genes with patient-level phenotypes in the setting of complex hierarchical structures observed in cancer cells. To simulate a complex dataset with hierarchical technical and biological effects, we utilized the estimated coefficients from the linear mixed effect models for 175 modules estimated on our NSCLC atlas. The coefficients for a module in a cell were added together, including the intercept, the random effects (study, sample, cluster), and two fixed biological effects (histology, stage). The true residuals from the models were discarded to remove additional biases not captured by the original coefficients. To add relatively simple noise, the residual for each module in each cell was sampled from a Gaussian distribution,ϵ_i, j_∼ *N*(0,σ_i_), where σ_i_ was the estimated standard deviation of the true residuals for module. To create gene-level counts, we first reversed the log2-transformed module scores to return them to their original scale. Then for each module, the counts for 25 genes were generated by sampling the total module count from a Dirichlet distribution with a concentration parameter of 0.5 (25 genes * 175 modules = 4,375 genes total). We applied several NMF or factorization algorithms including GeneNMF which implements the procedure outlined in Gavish *et al*^26^ and represents “state-of-the-art” in terms of module detection across tumors as well as hdWGCNA another co-expression module detection tool^83^. Standard NMF was also tested using the implementation from the RcppML package. We used the suggested metrics from each package for choosing the number of modules if available. The LME model was applied to the module scores to test for associations with histology and stage while controlling for study. The “ground truth” effect sizes for histology and stage for each gene were assumed to be the same as the overall module. One major challenge with interpretation of results for NMF-related tools is that each gene has membership in all modules to some degree, and it is not always clear how to assign genes to an individual module. Therefore, we utilized 3 different approaches to assign genes to NMF modules: 1) *<u>Max</u>*: each gene is assigned to the factor with the maximum weight across all factors; 2) *<u>Corr</u>*: Each gene is assigned to a single factor based on the best Pearson correlation between the expression for that individual gene and the estimated module activities across cells; and 3) *<u>Top</u>*: The top 25 genes in each factor are assigned to that factor and all other genes are ignored.

### Comparison of modules from different tools in the NSCLC cancer cell atlas

We compared Celda modules against the gene programs identified by alternative methods and parameterizations to assess recovery and coherence. The methods included geneNMF, hdWGCNA, and standard NMF implemented in the RcppML package. geneNMF metaprograms were identified using different combinations of nMP, the number of metaprograms; specificity.weight, which controls overlap between metaprograms; min.confidence, the minimum fraction of individual programs in which a gene must recur to be retained; weight.explained, which controls metaprogram size by keeping the genes that explain a set fraction of loadings in the recurrent gene set. hdWGCNA modules were detected with different values of minModuleSize, which is the minimum number of genes retained in a module during dynamic tree cutting, and mergeCutHeight, which is the eigengene dissimilarity threshold below which modules are merged. NMF was run at ranks k = 10 to 200 in increments of 10, and three ranks were selected to cover a range of factor numbers. To determine whether a module recovered by another method corresponded to a Celda module, we tested for significant overlap in gene membership between them. For this comparison, we retained Celda modules containing 5 or more genes and assessed overlap significance using a one-sided Fisher’s exact test. For standard NMF, we used the three gene-to-factor assignment approaches described above. To evaluate gene-to-module (or gene-to-factor) coherence for Celda and standard NMF, we computed the correlation between each gene’s expression and the activity of its corresponding module/factor, averaged these correlations within each module or factor, and compared the resulting distributions between methods. We assessed the biological interpretability of Celda modules and NMF factors by Reactome over-representation analysis using ReactomePA^73^. We then applied the same LME model to the NMF factor activities across all cells. Because RcppML NMF, unlike Celda, does not inherently cluster cells into populations, cell-cluster labels were omitted from the random effects. The percentage of variability in module or factor activity explained by the fixed effects was then compared between Celda modules and NMF factors. P-values for histology and stage were adjusted within each method using the Benjamini-Hochberg procedure.

### Comparing pooled and non-pooled study designs for module detection

The primary analysis used for biological discovery was to run Celda on cells from all studies (i.e., a pooled multi-study design) and then use LMEs to estimate the contribution of biological and technical effects. A complementary approach is to run Celda on cells from each individual study and then merge recurrent modules on the basis of overlap and/or clustering statistics. To compare results from “pooled” and “non-pooled” designs, we ran Celda on cells from each study cohort separately and compared the modules from the pooled approach to those from each study-specific run. Each of the 175 pooled modules was compared to the top 30 genes, ranked by probability, of every study-specific module using a one-sided hypergeometric test, requiring a minimum overlap of two genes. Because a single pooled module may be split across several study-specific modules, the hypergeometric p-values for a given pooled module were combined across all modules within a study using Fisher’s combined probability test, yielding one score per pooled module per study. A pooled module was considered recovered in a study if the combined p-value was < 1 × 10^-5^. The reverse comparison was performed identically with each study-specific module as the query and the pooled modules as the reference. To assess preservation of clinical associations, for each histology-or stage-associated pooled module we selected the study-specific module with the smallest overlap p-value within each study and tested its association with histology (in the four studies containing both LUAD and LUSC tumors) or stage (in the five LUAD-containing studies with both early-and late-stage tumors) using an LME with Sample and Cluster as random effects. The Study term was omitted because each model was fit within a single study. P-values from these within-study tests are reported unadjusted. Module specificity across cell clusters in the pooled analysis was summarized with the *τ*index^84^, 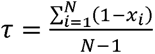, *N* where represents the total number of cell clusters and *x*_i_ represents the mean module expression in cluster *i* normalized to the maximum mean expression across all clusters. To assess the influence of cell-number imbalance across tumors, cells from each tumor were randomly downsampled to a maximum of 500 cells; tumors with fewer than 500 cells retained all cells. Celda was rerun on the downsampled cells from Wu et al. and separately within each study, and overlap with the original modules was assessed as described above.

## Acknowledgements

This work was supported by the National Library of Medicine R01LM013154 (J.D.C. and M.Y.), the Informatics Technology for Cancer Research (ITCR) 1U01CA220413 (J.D.C.), and the NCI Human Tumor Atlas Network (HTAN) Lung Pre-Cancer Atlas (PCA) 1U2CCA233238 (J.D.C., S.M., J.B.).

## Contributions

R.H., P.K., and N.Z. performed the statistical analyses. S.M., A.E.S., J.B., and M.Y. contributed to the study design and methodology. R.H., P.K. and J.D.C. wrote the manuscript. All authors read and approved the final manuscript.

## Data availability

Data for Kim *et al* was obtained from the Gene Expression Omnibus (GEO) database under accession number GSE131907 and is available at the following URL: https://www.ncbi.nlm.nih.gov/geo/query/acc.cgi?acc=GSE131907. Data for Laughney *et al* was obtained from the GEO database under accession number GSE123904 and is available at the following URL: https://www.ncbi.nlm.nih.gov/geo/query/acc.cgi?acc=GSE123904. Data for Wu *et al* was obtained from the GEO database under accession number GSE148071 and is available at the following URL: https://www.ncbi.nlm.nih.gov/geo/query/acc.cgi?acc=GSE148071. Data for Dost *et al* was obtained from the GEO database under accession number GSE149655 and is available at the following URL: https://www.ncbi.nlm.nih.gov/geo/query/acc.cgi?acc=GSE149655. Data from Xing *et al* was shared by the original authors and is available in the Genome Sequence Archive (GSA) database under accession number HRA000154 at the following URL: https://ngdc.cncb.ac.cn/gsa-human/browse/HRA000154. Data from Yanagawa *et al* was shared by the original authors and is available in the Human Tumor Atlas Network (HTAN) data portal under the Lung Pre-Cancer Atlas at the following URL: https://humantumoratlas.org/data-access. Data from Sinjab *et al* was shared by the original authors and is available in the GEO database under accession number GSE222901 at the following URL: https://www.ncbi.nlm.nih.gov/geo/query/acc.cgi?acc=GSE222901. Data from Qian *et al*, Zilionis *et al*, Goveia *et al*, and Chen *et al* are available from Salcher *et al* under the following URL: https://lambrechtslab.sites.vib.be/en/pan-cancer-blueprint-tumour-microenvironment-0. Data from Bischoff *et al* was obtained from the following URL: https://codeocean.com/capsule/8321305/tree/v1.

## Code availability

Code to reproduce the analyses related to gene module detection and characterization is available at https://github.com/campbio-manuscripts/NSCLC_GeneModules.

## Supplementary Figure Overview

**Supplementary Figure 1.**
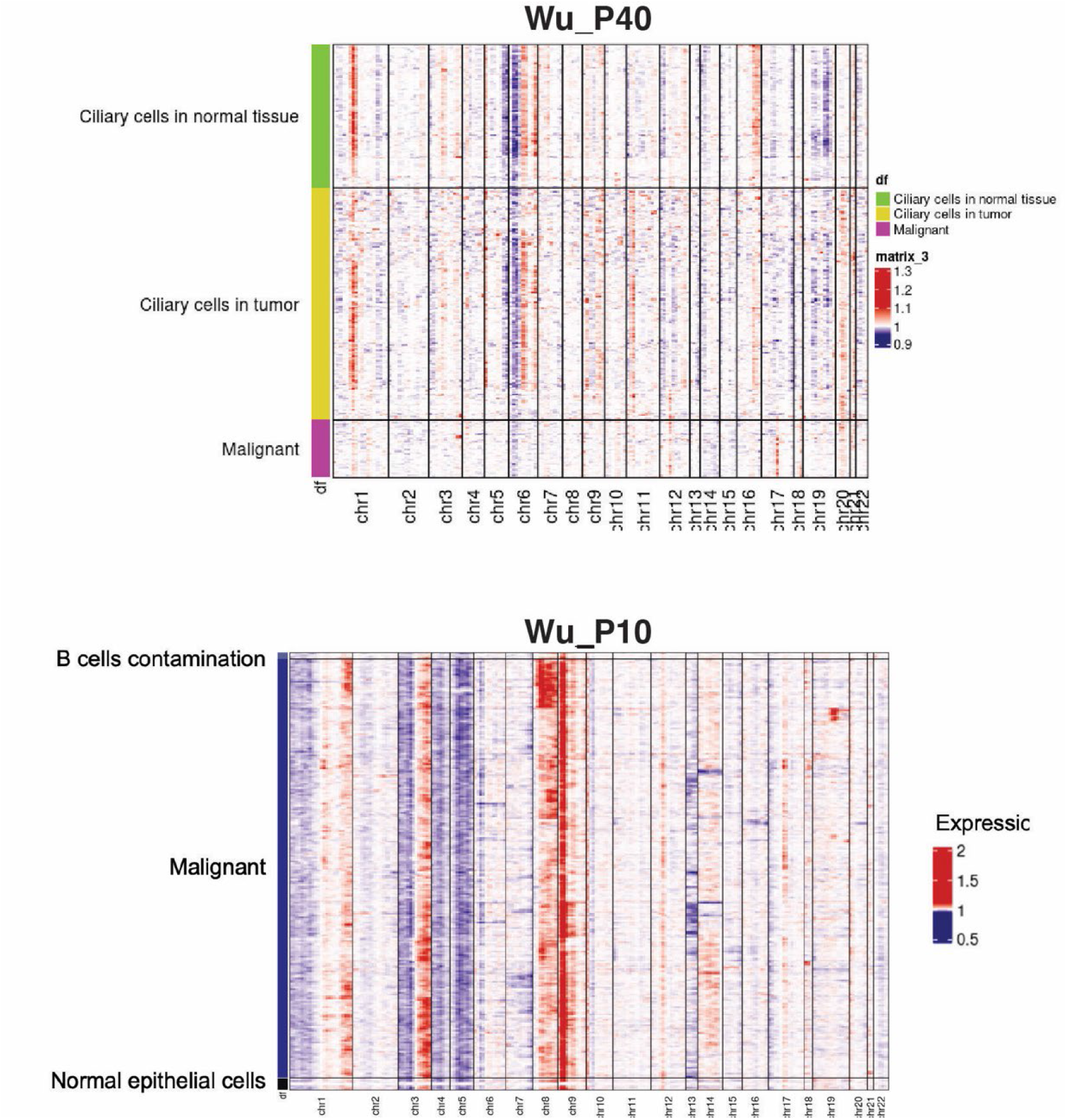
Identification of cancer cells and removal of misclassified normal ciliated cells and B-cell doublets. **(A)** inferCNV profiles for Wu_P40. Two groups of epithelial cells were initially classified as malignant based on apparent copy number alterations. When ciliated cells from a normal lung sample (Yanagawa et al., Case 4-Normal) were included in the observation set, one group (Ciliated cells in tumor) showed the same profile as the normal ciliated cells and a different profile from the malignant cells, indicating that the apparent alterations reflect the ciliated expression program rather than copy number changes. These cells were removed from the atlas. **(B)** inferCNV profiles for Wu_P10. Cells with high expression of a B-cell module (top) shared the copy number profile of the malignant cells, consistent with malignant–B-cell doublets, and were removed. Normal epithelial cells from the same sample are shown at the bottom for comparison. Rows are cells, columns are genes ordered by chromosome, and color indicates inferCNV-modified expression relative to the reference. After removal of both groups, Celda was rerun and the ciliary and B-cell modules were no longer detected.

**Supplementary Figure 2.**
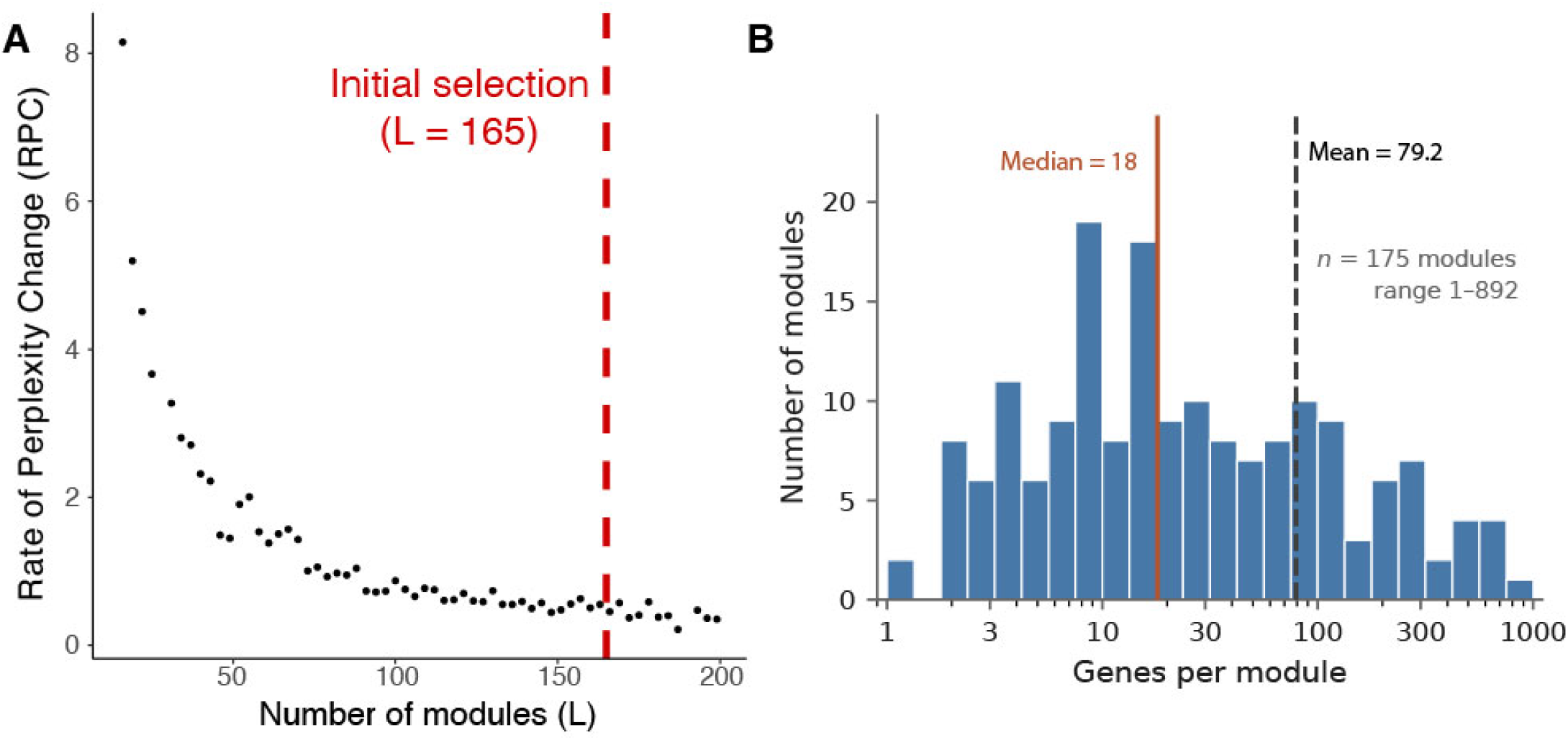
Rate of Perplexity Change (RPC) to detect numbers of modules and the distribution of genes per module. **(A)** Perplexity is a measure of uncertainty in the value of a sample from a discrete probability distribution. Like other topic models, Celda uses perplexity to determine key parameters such as the number of cell clusters (K) or number of modules (L). This plot shows the RPC for each choice of L as previously described^1^. An initial L of 165 was selected. 10 modules were further split using the *splitModule()* function in the celda package. **(B)** The number of genes in modules ranged from 1 to 892 (median = 18, mean = 79.2). 55 modules contained fewer than 10 genes, while 36 modules contained 100 or more genes.

**Supplementary Figure 3.**
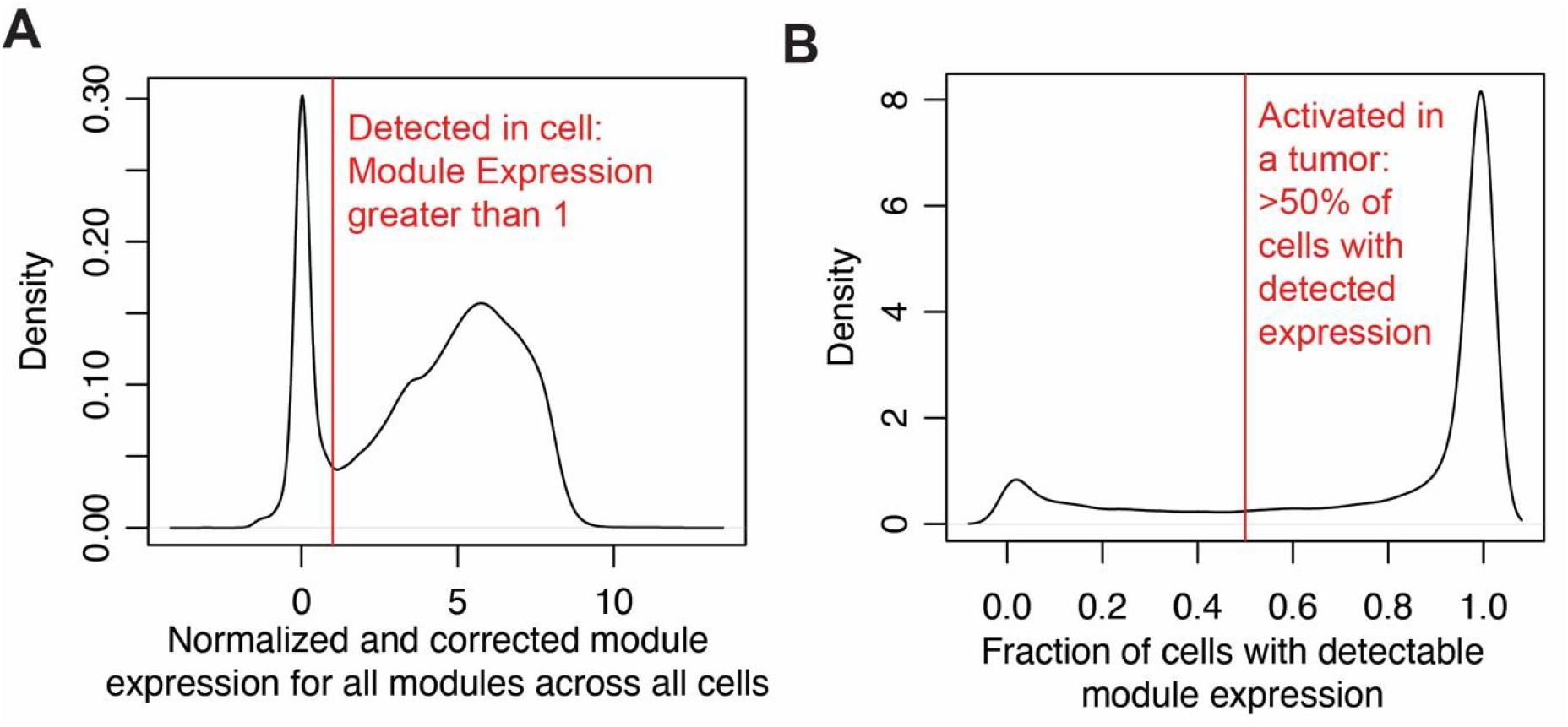
Detection of modules in cells and tumors. **(A)** The density of normalized and corrected module expression was plotted for all cells and all modules. A module was considered “detected” in an individual cell if it had an expression value of at least 1. **(B)** The density of the fraction of detected cells was plotted for all tumors and all modules. A module was considered active in an individual tumor if it was detected in at least 50% of cells.

**Supplementary Figure 4.**
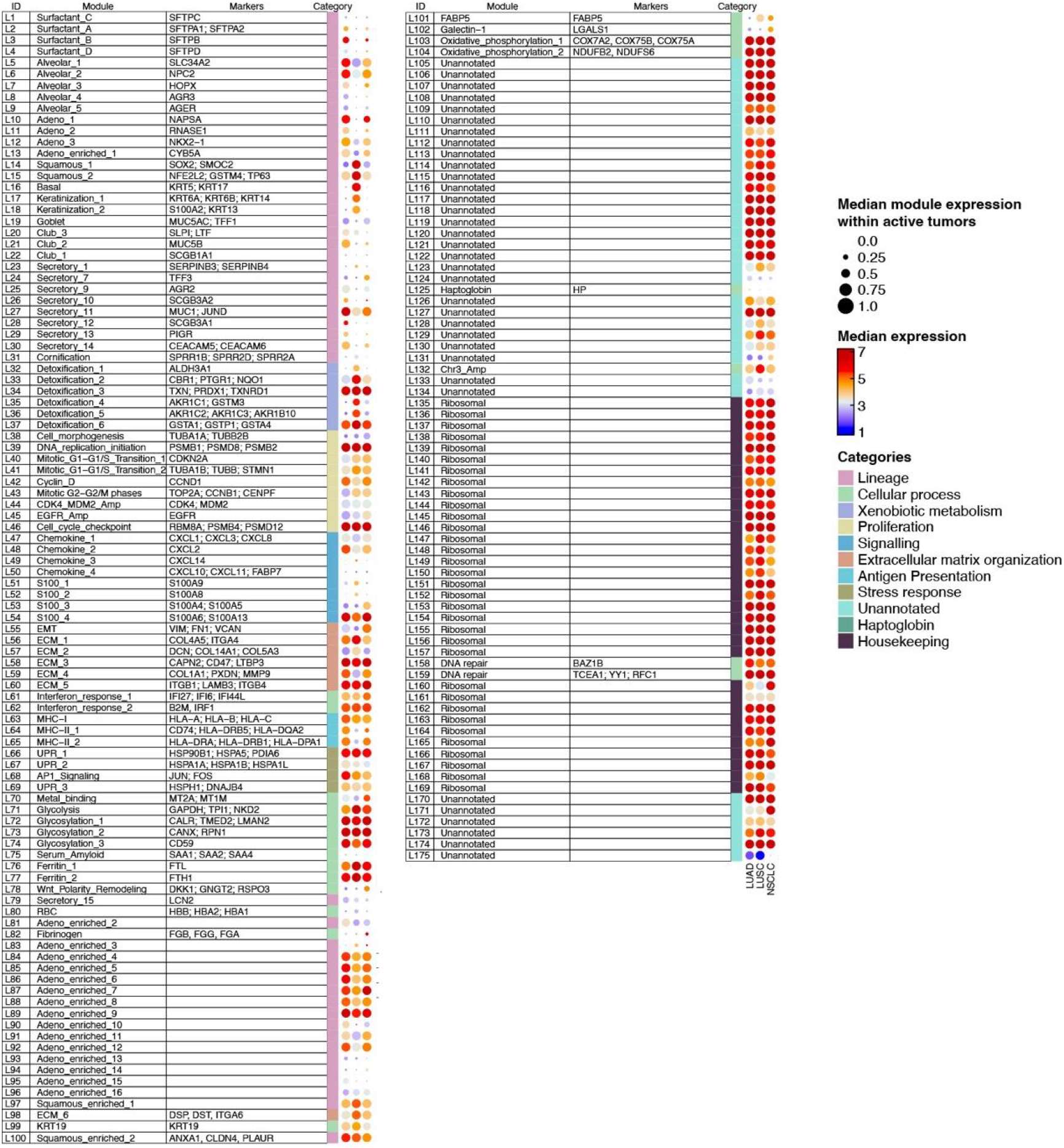
The landscape of gene co-expression modules in non-small cell lung cancer. The table lists the curated modules with corresponding key genes used for annotation. A module was considered “detected” in an individual cell if it had a log2 normalized expression value of at least 1. A module was considered active in an individual tumor if it was detected in 50% of cells. The size of the dot for each module indicates the proportion of activated tumors within LUAD, LUSC, or unspecified NSCLCs. The color of the dot indicates the median expression of the module within the activated tumors. All modules are shown regardless of biological category.

**Supplementary Figure 5.**
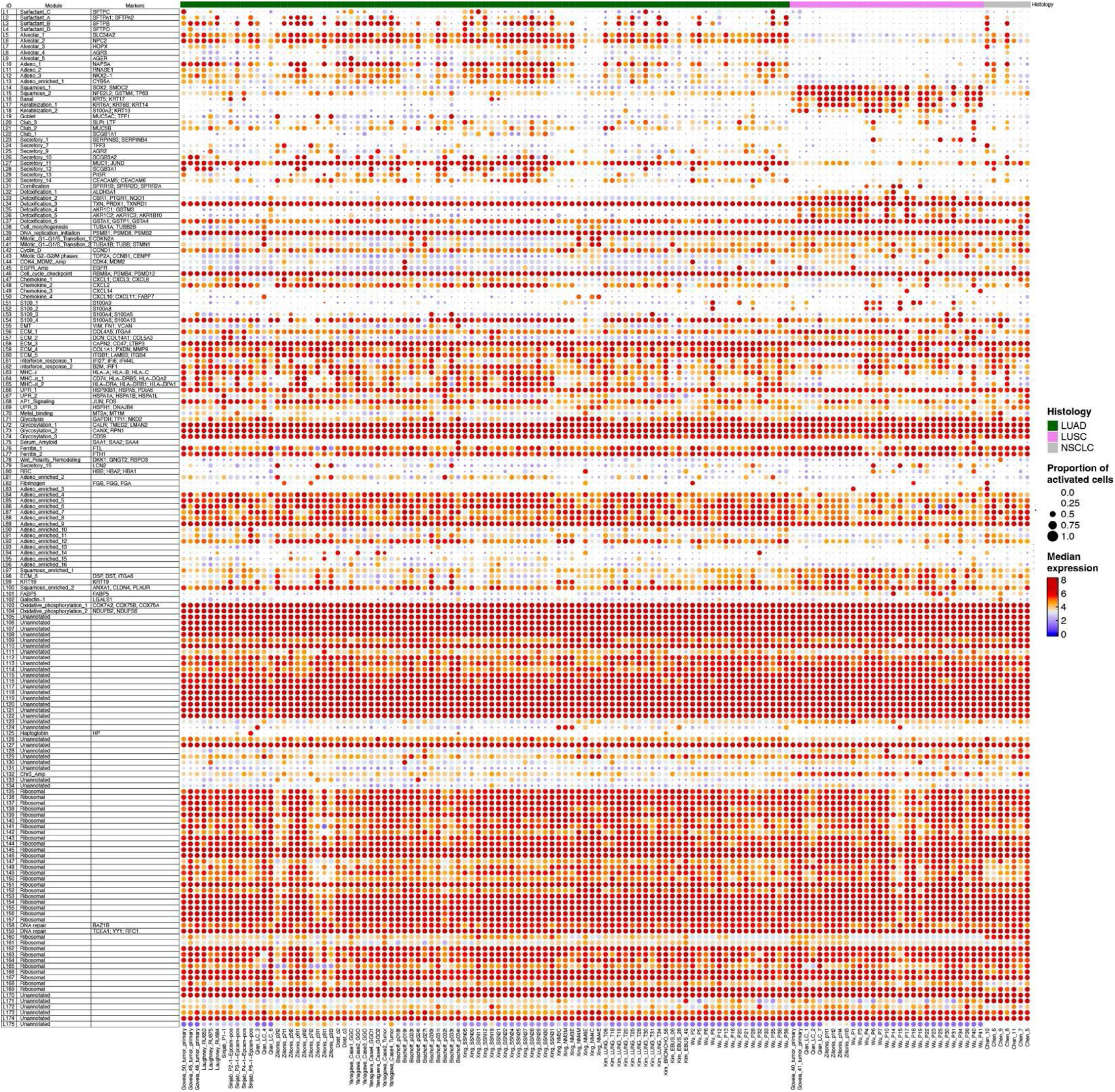
Landscape of module detection rates and expression in individual tumors. A module was considered “detected/activated” in an individual cell if it had a log2 normalized expression value of at least 1. Each row is a module, and each column is a tumor. The size of the dot for each module indicates the proportion of activated cells within each tumor. The color of the dot indicates the median expression of the module across cells within that tumor.

**Supplementary Figure 6.**
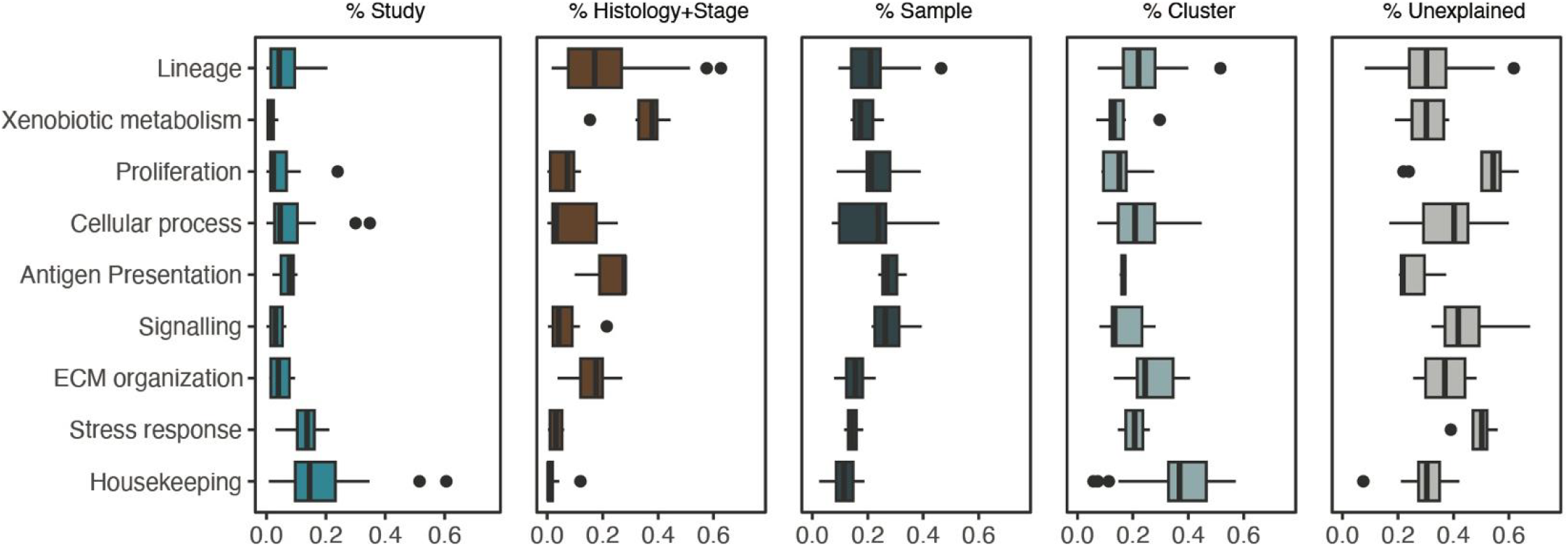
Percent variation explained by fixed and random effects for different biological categories. Linear mixed effect (LME) modeling was used to quantify the variability and association of each module with biological and technical covariates. Each cell was treated as a repeated measure using Study, Sample, and Cluster as fully nested random effects while Histology and Stage were included as fixed effects. Each point represents the percentage of variability for a module for that term. Modules are grouped by overall biological category.

**Supplementary Figure 7.**
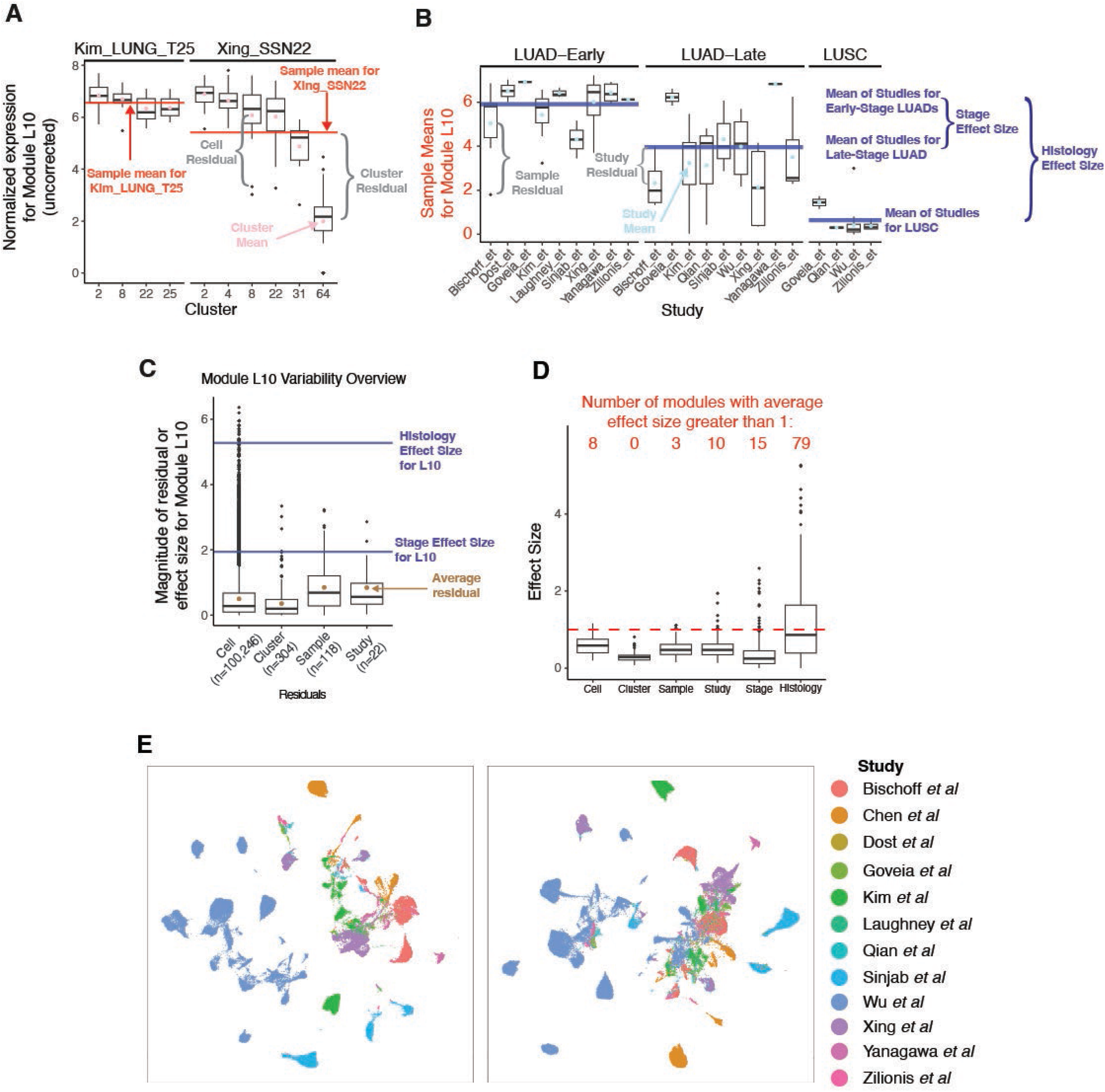
Determining effect sizes of each module for cell, cluster, sample, study, stage, and histology. The goal of this analysis was to compare the average effect size or “magnitude of change” for different variables in the LME model. To illustrate how this was measured, we show how the effect sizes were calculated for module L10. The analysis was limited to cells coming from clusters with at least 20 cells. This included 100,246 cells from 304 clusters from 118 tumors. Here the “Cluster” term refers to a combination of the tumor and cluster label. For example, Celda cluster K2 in Kim_LUNG_T25 will be considered a different cluster than K2 in Xing_SSN22. **(A)** Each point represents the module expression levels of L10 in cells. Two tumors with multiple clusters are shown as examples, Kim_LUNG_T25 and Xing_SSN22. For each cell cluster, the cluster mean was calculated and the “cell residual” was determined by taking the absolute value of the difference between each cell and its respective cluster mean. The “sample mean” for each tumor was calculated as the mean of the cluster means. This ensures that each cluster will be treated equally when calculating the sample means and will not be more heavily influenced by clusters with more cells. **(B)** Each point represents the sample mean for each tumor for module L10. Samples are first grouped by three biological categories, LUAD Early-Stage, LUAD Late-Stage, and LUSC, and then by study within each major group. The “study” means (light blue) were calculated by averaging the sample means for each study within each biological group. The “sample residual” is the absolute difference between a sample mean and its respective study mean within each biological group. The biological group means (dark blue) were calculated by averaging the study means within each biological group. The Stage effect size was calculated by taking the absolute difference between the LUAD Early-Stage group mean and the LUAD Late-Stage group mean. The Histology effect size was calculated by taking the absolute difference between the LUAD Early-Stage group mean and the LUSC group mean. **(C)** The magnitude of average residuals (brown points) was compared to effect sizes for module L10. The effect sizes for Histology (5.27) and Stage (1.94) are larger than the means of residuals for the Study (0.84), Sample (0.85), Cluster (0.35), and Cell (0.50) variables demonstrating that the magnitude of the biological effects of Histology and Stage are greater than the average differences between cells, cluster, samples, and studies. The effect sizes and mean residuals for each term for all modules are listed in **Supplementary Table 9**. **(D)** The average effect size (i.e. magnitude of change) for the possible sources of variability in each module was also examined. When counting modules with at least an absolute log2 difference greater than 1, we find that histology affected the largest number of modules (n=79) followed by Stage (n=15). Study was the third largest factor with 10 modules having an average difference greater than 1. **(E)** When using all modules and uncorrected module expression, the global variation in the UMAP was driven by differences between the *Wu et* al study, which used a different single cell assay, and other studies. In contrast, when using the corrected module expression and limiting to modules with <25% variability from the Study effect, the largest global difference was between LUADs and LUSCs. In general, the technical variability induced by study-specific differences largely affect broadly expressed housekeeping and unannotated modules while the remaining modules enriched for other processes have larger effects from biological factors such as Histology and Stage.

**Supplementary Figure 8.**
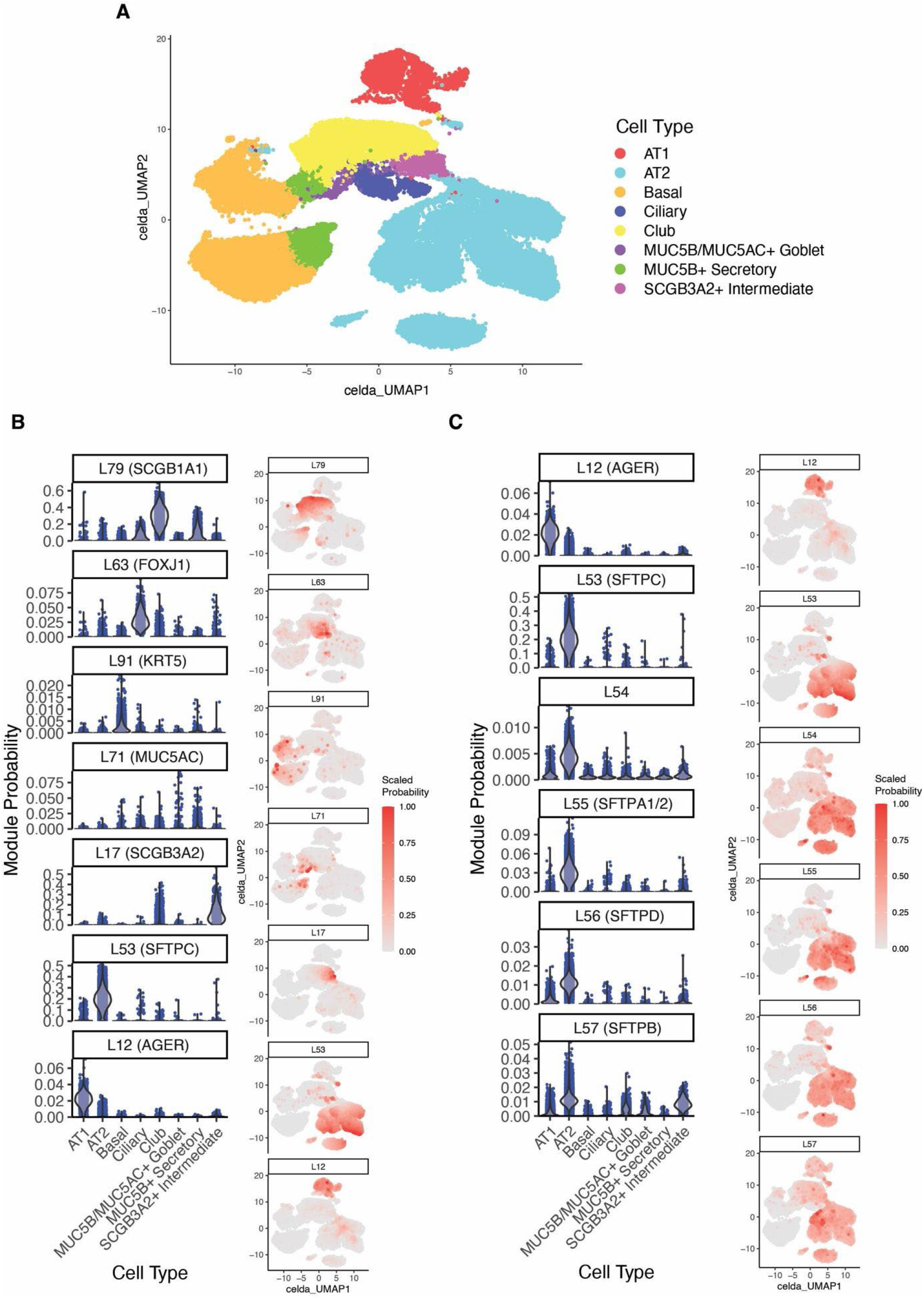
Detection of alveolar modules in normal lung epithelial cells. Single-cell RNA-seq data from non-cancerous lung tissue specimens was obtained from Basil *et al*^2^ from 10 subjects without lung cancer. After initial clustering with Celda, lung epithelial cells were isolated and reclustered. **(A)** 125 gene modules were identified among the basal, secretory MUC5B+, club, goblet, ciliary, intermediate SCGB3A2+, and alveolar AT1/2 cells with Celda. **(B)** The distribution of probabilities for modules of each of the major cell types are shown. Full lists of genes in these modules can be found in **Supplementary Table 10**. Module probabilities are defined by the fraction of counts for the genes in that module out of the total number of counts in each cell. **(C)** Modules denoting AT1 and AT2 cells and that contained previously characterized marker genes were identified. L12 was used for AT1 cells and contained AGER, CAV1, and RTKN2. Several modules containing one or more pulmonary surfactant genes were enriched in AT2 cells including L53 (SFTPC), L55 (SFTPA1 and SFTPA2), L56 (SFTPD) and L57 (SFTPB). Note that L57 was also expressed in the SCGB3A2+ alveolar transitional cells. These modules were used for the lineage infidelity analysis of normal lung cells in Figure 2. Gene modules that had at least 0.5% of the total counts in a cell were considered detected in that cell.

**Supplementary Figure 9.**
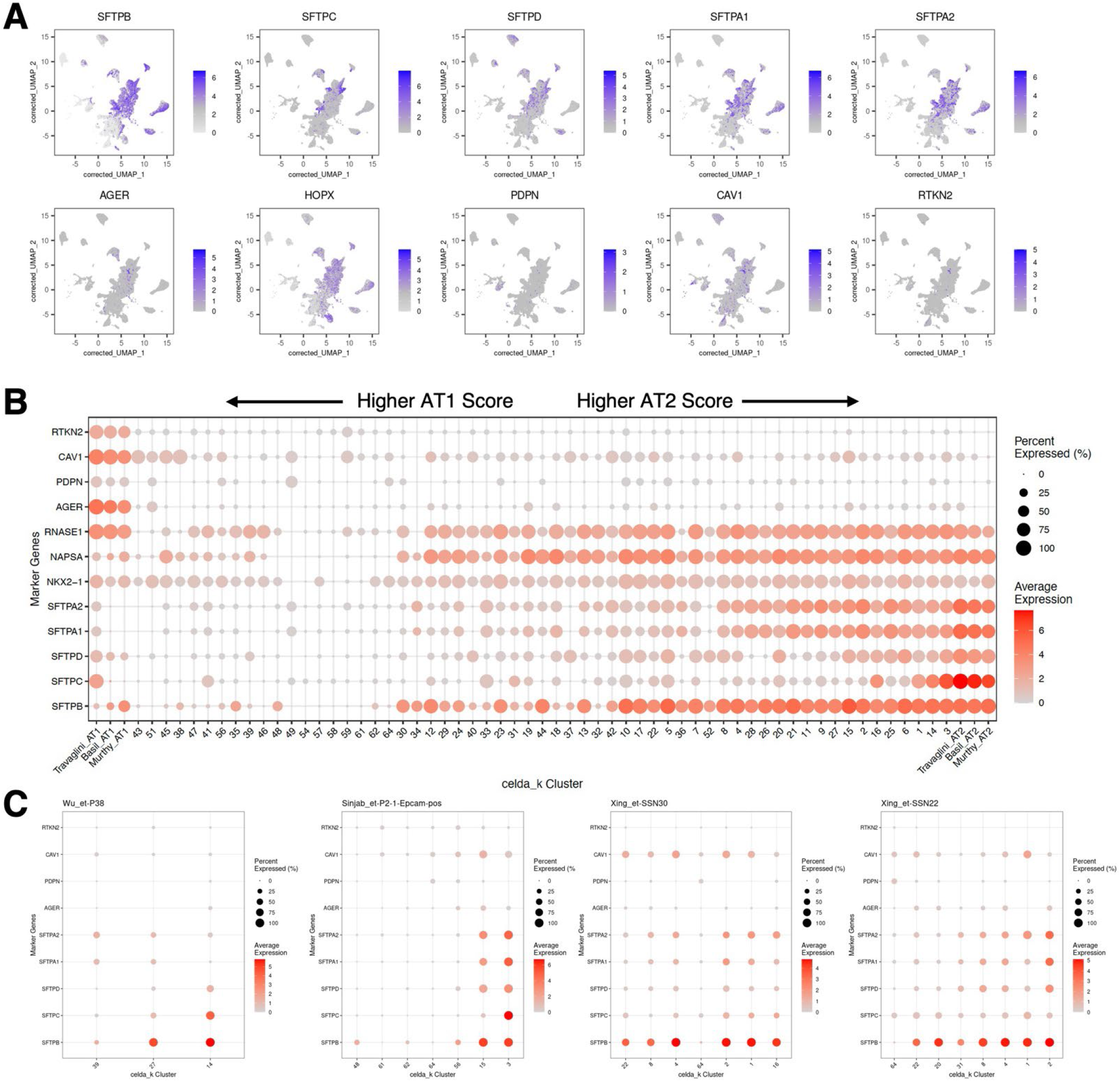
Marker-level analysis of AT1 and AT2 lineage identity in LUAD cancer cells. **(A)** Expression of AT2 markers (SFTPB, SFTPC, SFTPD, SFTPA1, SFTPA2; top row) and AT1 markers (AGER, HOPX, PDPN, CAV1, RTKN2; bottom row) on the UMAP of all cancer cells. The surfactant genes show distinct spatial distributions, consistent with their assignment to different Celda modules, whereas SFTPA1 and SFTPA2 share a similar distribution, consistent with their assignment to the same module. AT1 markers are sparsely expressed, with the exception of HOPX, which is broadly expressed across early-stage LUAD cells and also shows moderate expression in normal human AT2 cells. **(B)** Dot plot of AT1 markers (RTKN2, CAV1, PDPN, AGER), general alveolar markers (RNASE1, NAPSA, NKX2-1), and AT2 surfactant markers (SFTPA2, SFTPA1, SFTPD, SFTPC, SFTPB) across Celda cell clusters. Cells were scored with AT1 and AT2 gene signatures, and clusters were ordered by the median of the AT1 score minus the AT2 score, with the highest relative AT1 score on the left. AT1 and AT2 cells from three normal lung datasets (Travaglini et al., Basil et al., and Murthy et al.) were scored and included as reference populations at the left and right ends. Dot size indicates the percentage of cells in the cluster expressing the gene and color indicates average expression. Normal AT1 cells express high levels of AGER, RTKN2, and CAV1, whereas normal AT2 cells express high levels of all five surfactant genes. No cancer cell cluster expresses AGER or RTKN2 above trace levels. The cancer clusters with the highest relative AT1 scores are those that have lost surfactant and general alveolar markers rather than gained AT1 markers. Across cancer clusters, SFTPB, NAPSA, and RNASE1 are the most broadly retained, SFTPA1, SFTPA2, and SFTPD are retained in a subset of clusters, and SFTPC is retained at appreciable levels only in the clusters most similar to normal AT2 cells. **(C)** Dot plots of the same AT1 and AT2 markers across Celda cell clusters within four representative LUADs: Wu et al. P38, Sinjab et al. P2, and two subsolid nodules from Xing et al. (SSN30 and SSN22). Within each tumor, clusters differ in which surfactant genes are detected (for example, SFTPB with SFTPC and SFTPD versus SFTPB with SFTPA1/SFTPA2 versus no surfactant expression within Wu et al. P38), demonstrating that multiple surfactant-marker combinations coexist within individual tumors rather than reflecting only tumor-to-tumor differences.

**Supplementary Figure 10.**
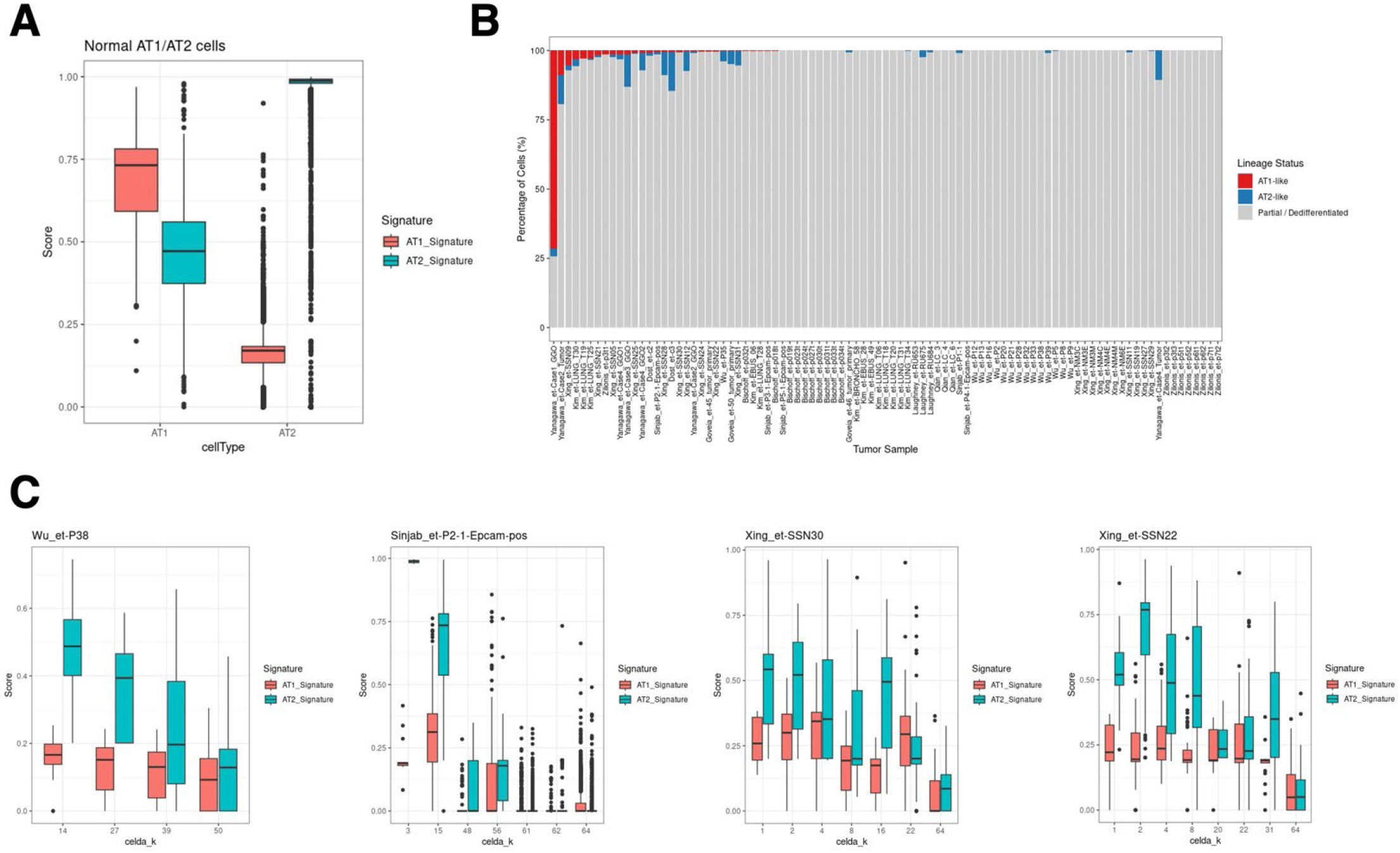
Quantification of AT1 and AT2 lineage identity in LUAD cancer cells with signature scores. **(A)** AUCell scores for the AT1 and AT2 signatures in normal AT1 and AT2 reference cells from three normal lung datasets (Travaglini et al., Basil et al., and Murthy et al.). Normal AT1 cells have high AT1 scores and intermediate AT2 scores, whereas normal AT2 cells have near-maximal AT2 scores and low AT1 scores. These distributions were used to define classification thresholds: a cell was called AT1-like if its AT1 score was at or above the 25th percentile of normal AT1 cells and its AT2 score at or below the 75th percentile of normal AT1 cells, and AT2-like if its AT2 score was at or above the 25th percentile of normal AT2 cells and its AT1 score at or below the 75th percentile of normal AT2 cells. All other cells were classified as partial/dedifferentiated. **(B)** Percentage of malignant cells in each of the three categories for every LUAD, ordered by the fraction of AT1-like cells. AT1-like cells were rare across the atlas and were only the predominant state in one premalignant sample, Yanagawa Case1_GGO. Cells meeting the full AT2-like criteria were also uncommon, and the large majority of malignant cells in nearly every tumor showed partial or absent expression of both programs. **(C)** AT1 and AT2 signature scores by Celda cell cluster within four representative LUADs (Wu et al. P38, Sinjab et al. P2, and subsolid nodules SSN30 and SSN22 from Xing et al.). Clusters within a tumor differ substantially in AT2 score, whereas AT1 scores remain low in all clusters and do not increase in clusters with reduced AT2 scores, demonstrating that loss of the AT2 program within a tumor is not accompanied by gain of an AT1 program in these samples.

**Supplementary Figure 11.**
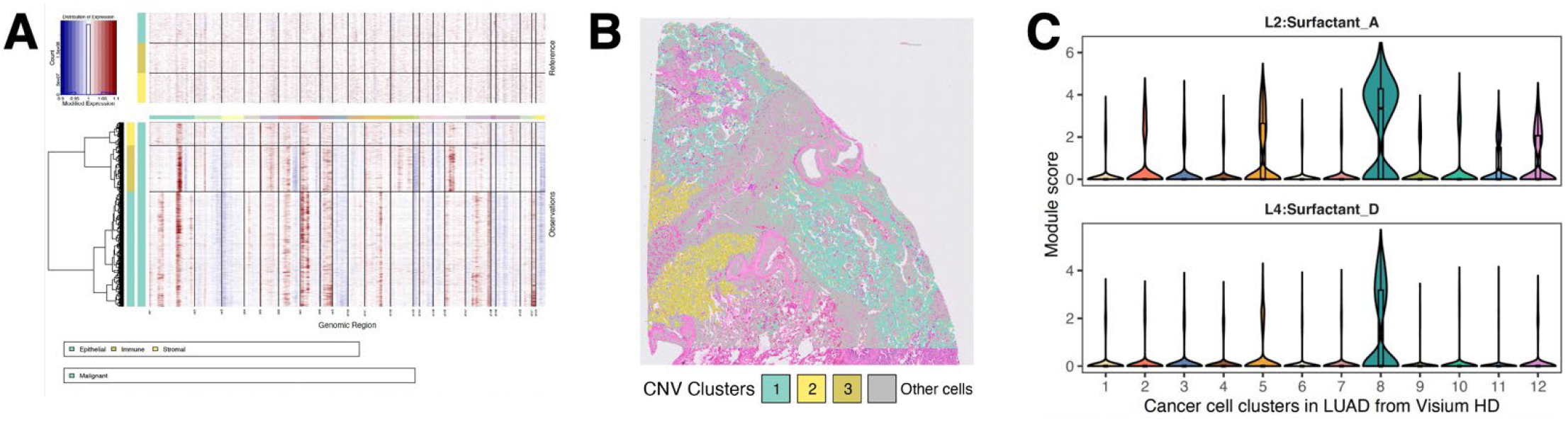
Examining intratumoral lineage infidelity using spatial transcriptomics. (**A**) inferCNV heatmap of epithelial cells from a LUAD profiled with Visium HD (10x Genomics). Stromal and immune cells from the same section were used as the reference. Rows are cells ordered by chromosomal region (columns), and the row annotation indicates the three CNV-defined clusters of cancer cells. (**B**) Spatial location of the three CNV-defined cancer cell clusters within the tissue section, with non-cancer cells shown in gray. Each CNV cluster occupied a distinct region of the tumor. (**C**) Violin plots of module expression for L2:Surfactant_A (SFTPA1/SFTPA2) and L4:Surfactant_D (SFTPD) across the 12 cancer cell clusters shown in Figure 2H. Both modules were largely restricted to the same cluster as L1:Surfactant_C (Figure 2I).

**Supplementary Figure 12.**
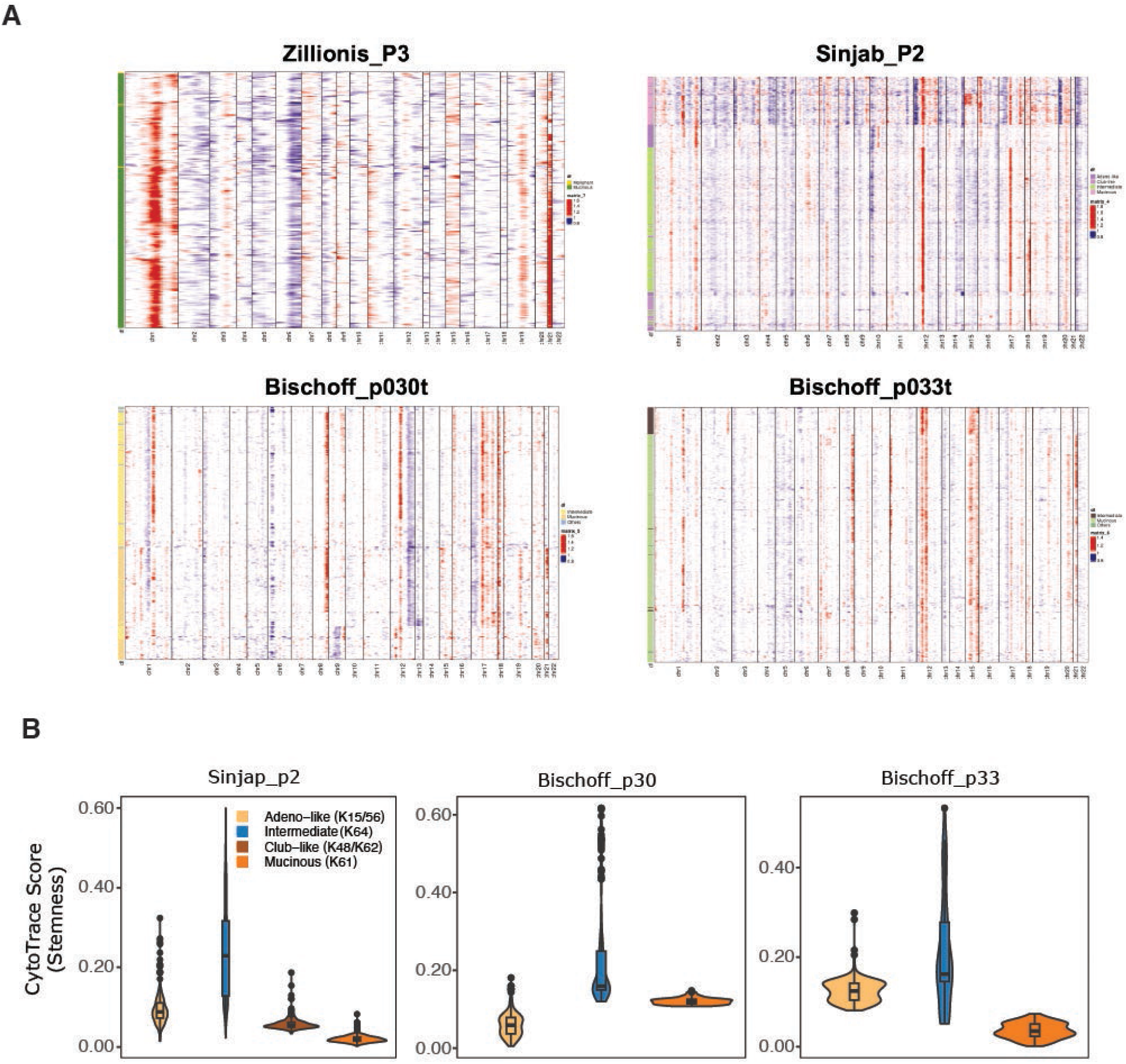
Copy number patterns and CytoTRACE scores in LUADs with high L19 mucinous module activity. Four tumors had at least 10 mucinous cells denoted by high expression of module L19 containing MUC5AC. **(A)** Cells from each tumor shared broad copy number alterations, indicating that the alveolar-, basal/squamous-like intermediate, club-like, and mucinous populations arose from a common malignant ancestor rather than from independent clones. Subclonal differences in copy number are visible within some tumors (e.g., Sinjab_P2). **(B)** CytoTRACE scores for the three tumors containing both mucinous and non-mucinous cancer cells. Intermediate basal-or squamous-like cells had the highest scores in each tumor, indicating the highest predicted developmental potential.

**Supplementary Figure 13.**
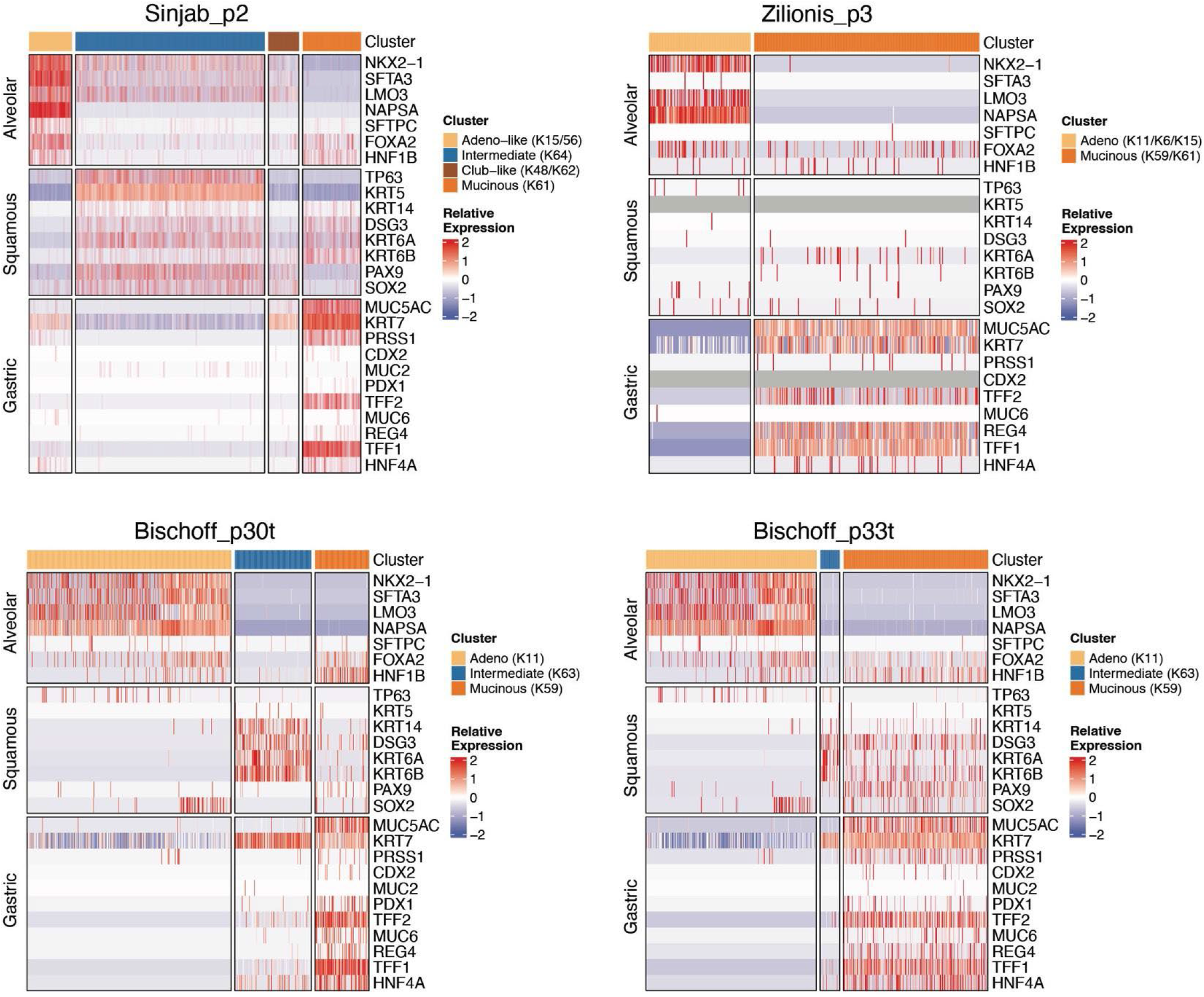
Expression of alveolar, squamous, and gastric markers in LUADs with high L19 mucinous module activity. Expression of markers for lung adenocarcinomas and alveolar cells, lung squamous cell carcinoma and basal cells, and gastric mucinous cells are shown for 4 lung adenocarcinomas (LUADs). These LUADs had a substantial number of mucinous-like cells with high levels of module L19 containing MUC5AC. Adeno-like cells were found within the same tumor as the mucinous cells in Sinjab_P2. For the other three tumors, intermediate and mucinous cells were compared to adeno-like cells from other tumors in the same dataset. The mucinous clusters in all four patients had higher levels of HNF4A, TFF1, and TFF2 compared to their alveolar counterparts. In contrast, the intermediate cells in Sinjab_P2, Bischoff_p030t, and Bischoff_p033t generally lacked expression of gastrointestinal markers but displayed different combinations of squamous and basal markers. The intermediate cells from Sinjab_P2 had higher expression of SOX2, TP63, and KRT5 while the intermediate cells from Bischoff_p030t and Bischoff_p033t had higher expression of DSG3, KRT6A, KRT6B. Zilionis_p3 only had mucinous-like cells. Markers for the alveolar, squamous, and gastric states were derived from Tata *et al*^3^.

**Supplementary Figure 14.**
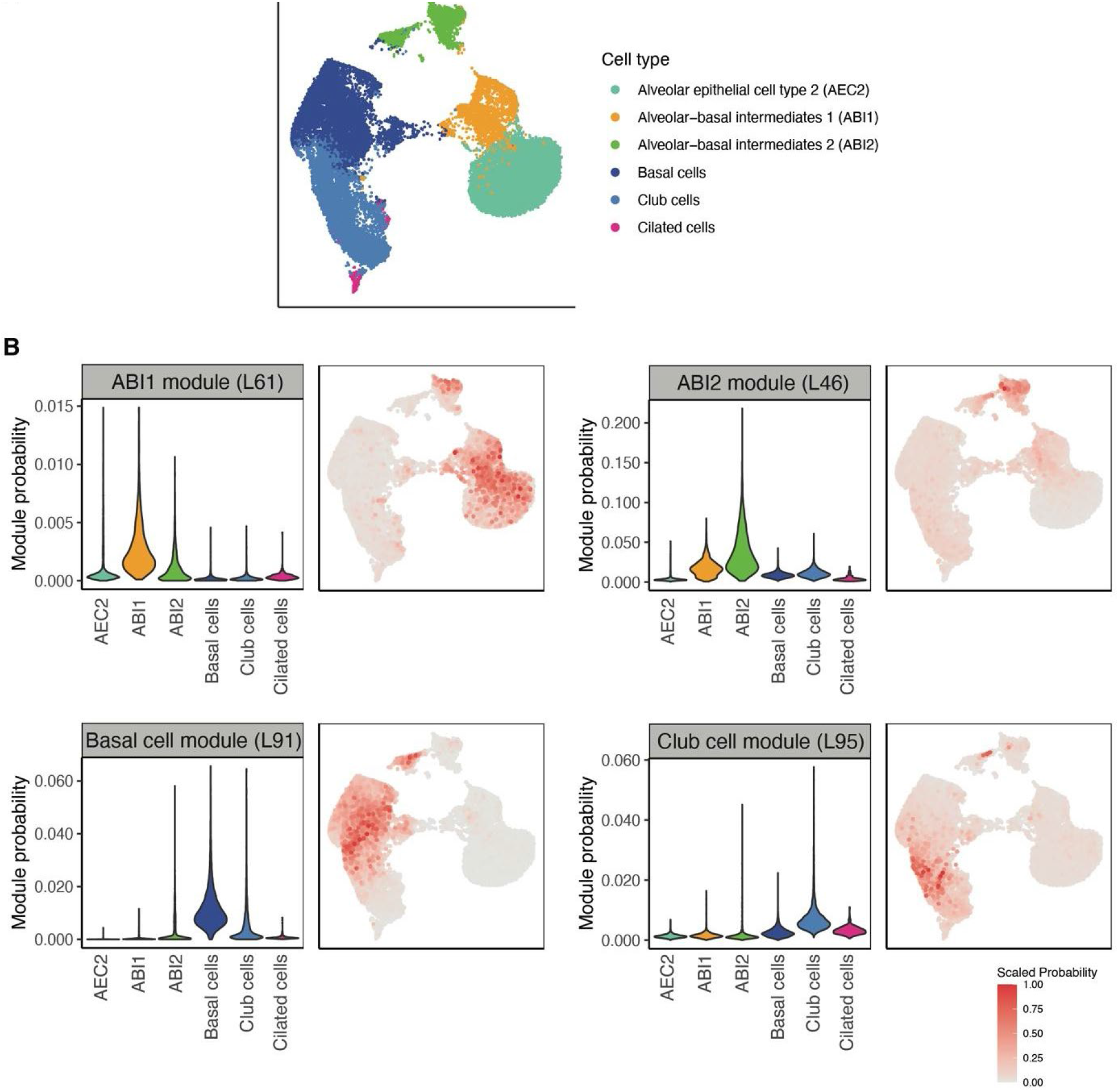
Defining modules associated with alveolar-to-basal transitions. Cells from the *in vitro* dataset from Kathiriya *et al*^4^ were reanalyzed to identify modules associated with alveolar-to-basal transitions. **(A)** Epithelial cells were identified with SingleR and cells were clustered using the Louvain algorithm on an SNN graph after batch correction with fastMNN. Cells were annotated with the same labels used in the original publication by examining marker genes and performing differential expression analysis between clusters. **(B)** Celda was also run on the dataset to identify 100 gene co-expression modules. Modules that could distinguish each major cell type were identified including module L11 containing NKX2-1 for the alveolar epithelial type 2 (AEC2) cells, module L61 for alveolar-to-basal intermediates 1 (ABI1), module L46 for the alveolar-to-basal intermediates 2 (ABI2), L91 for the basal cells, and L95 for the club-like secretory cells. The UMAPs show the probability of the module in each cell derived from Celda which has been “min/max” scaled to range from 0 to 1. These modules were scored in the LUAD tumors with mucinous cells to determine the contribution of transitional and basal-like modules to intermediate and mucinous cells in these LUADs.

**Supplementary Figure 15.**
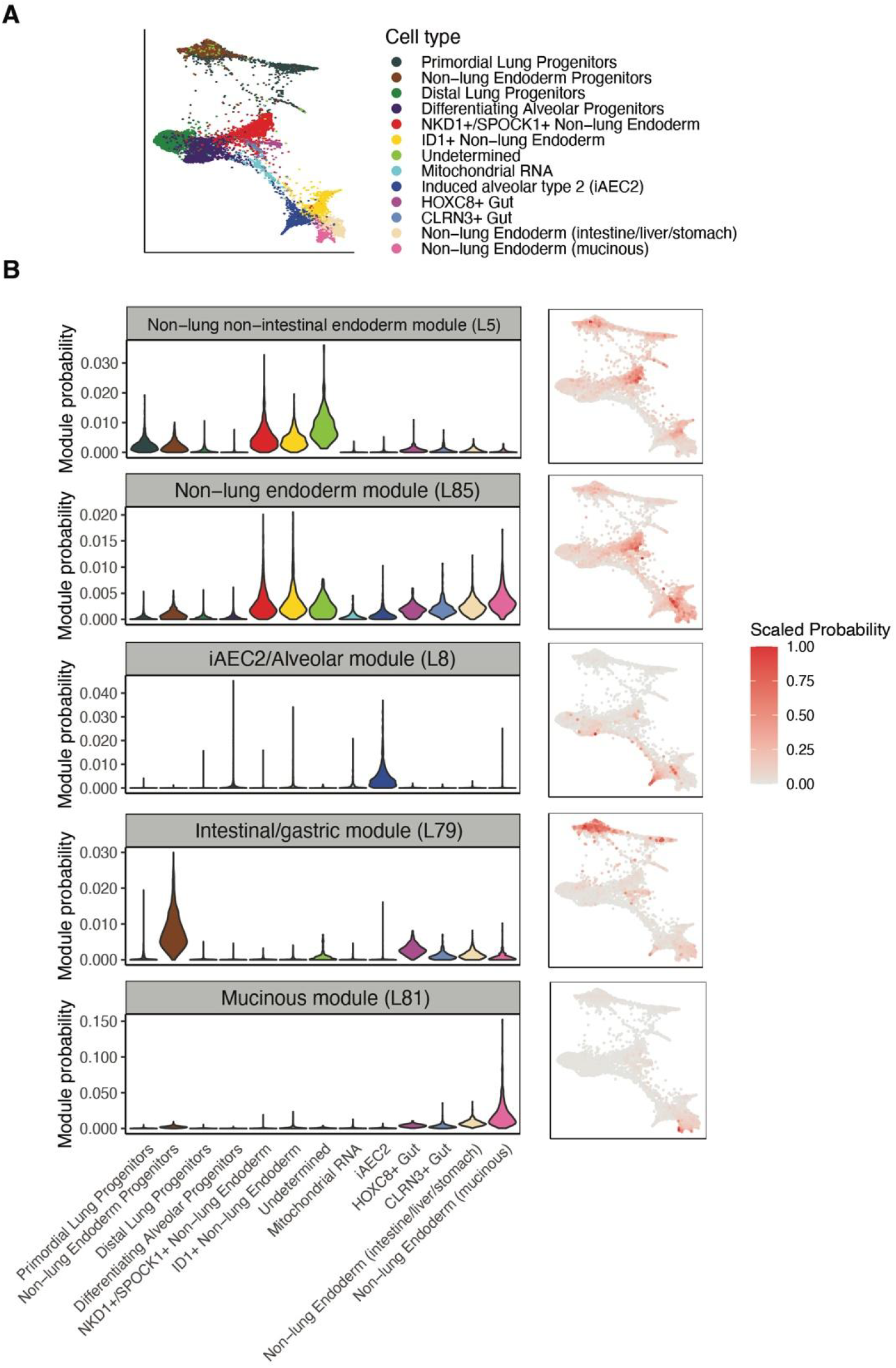
Defining modules associated with progenitor-to-alveolar and progenitor-to-gastric differentiation. Cells from the *in vitro* dataset from Hurley *et al*^5^ were reanalyzed to identify modules associated with progenitor-to-alveolar and progenitor-to-gastric/intestinal transitions. **(A)** The gene expression matrix, cell annotations, and UMAP coordinates were obtained from the original authors. **(B)** Celda was also run on the dataset to identify 100 gene co-expression modules. Modules that could distinguish each major cell type and transition were identified including module L5 expressed across all non-lung non-intestinal endoderm cells, L85 expressed across all non-lung endoderm cells (including gastric cells), L8 containing SFTPB expressed in alveolar type II cells, L79 containing CDX2 expressed in progenitor cells and across intestinal/gastric cells, and L81 containing MUC13 which is predominantly expressed in the mucinous cells. The UMAPs show the probability of the module in each cell derived from Celda which has been “min/max” scaled to range from 0 to 1. These modules were scored in the LUAD tumors with mucinous cells to determine the contribution of gastric modules to intermediate and mucinous cells in these LUADs.

**Supplementary Figure 16.**
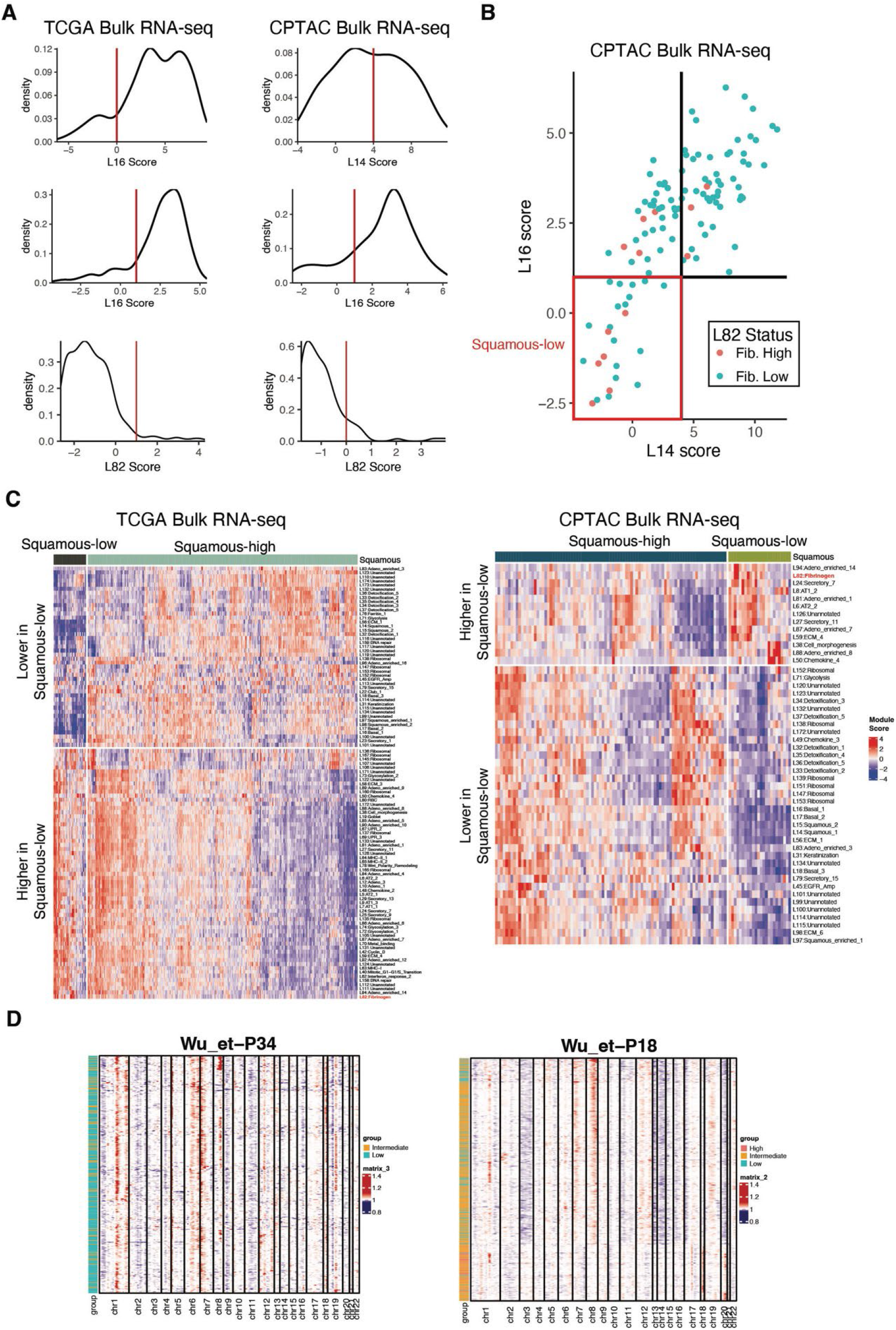
Validation of novel fibrinogen-high subgroup in LUSC using independent bulk RNA-seq datasets. **(A)** Each gene module was scored in TCGA and CPTAC LUSC bulk RNA-seq datasets. The density of the modules L14, L16, and L82 was examined to identify tumors with high or low levels of each module. **(B)** Squamous-low tumors were defined as those with low levels of both L14 and L16. **(C)** Modules that were significantly different between squamous-low and squamous-high tumors were identified in each cohort using a *t*-test (FDR q-value < 0.05 & absolute Log2 Fold Change > 1.5). **(D)** inferCNV profiles of squamous-low and squamous-high cancer cells from Wu_P34 and Wu_P18. Rows are cells, columns are genes ordered by chromosome, and color indicates inferCNV-modified expression relative to the reference. For both tumors, squamous-low and squamous-high cells largely shared the same copy number profile exemplified by intermixing in the clustering diagram. Neither tumor showed the 3q gain characteristic of LUSC.

**Supplementary Figure 17.**
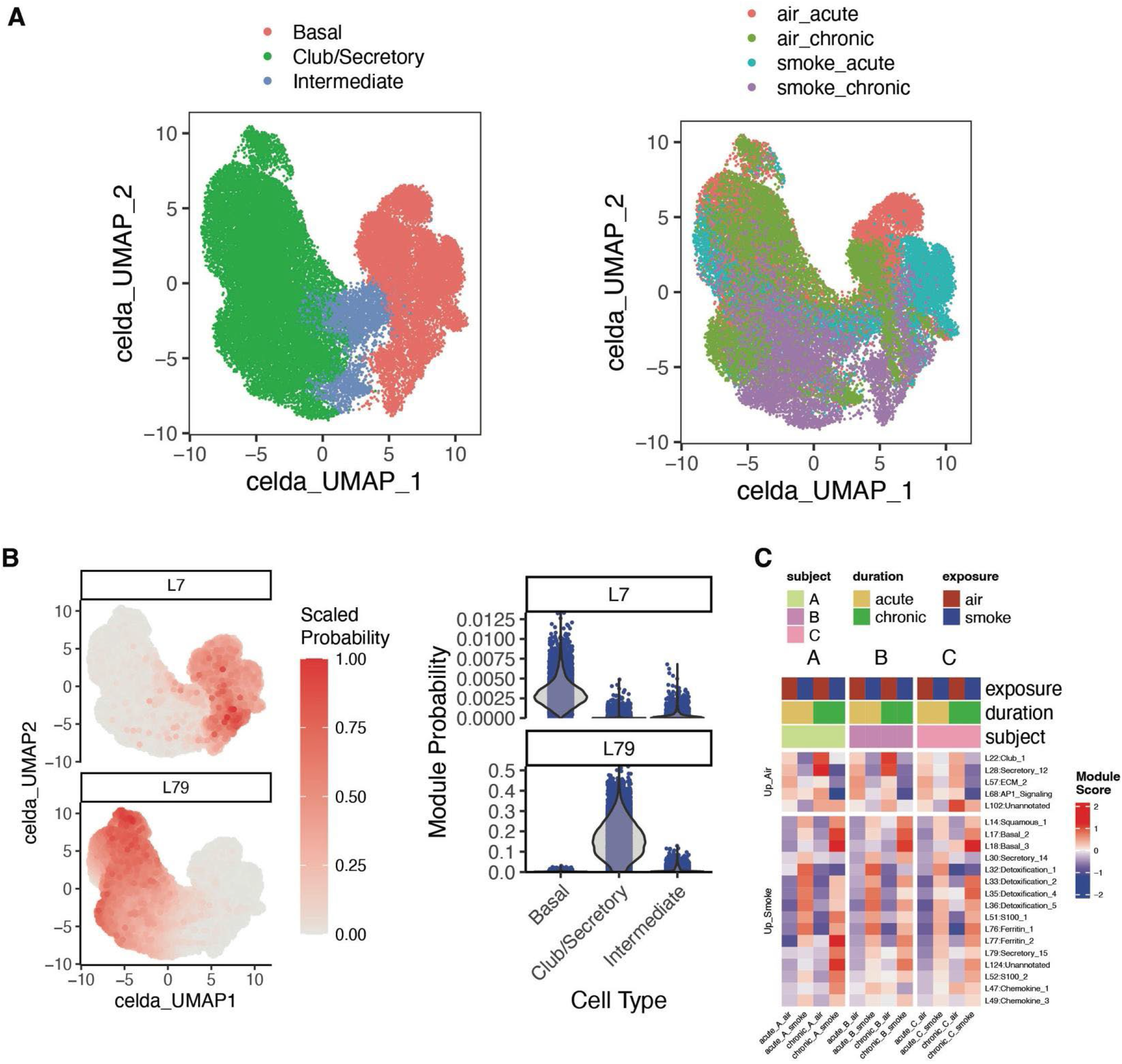
Characterization of epithelial cell states exposed to cigarette smoke after differentiation at an air-liquid interface (ALI). Single cell RNA-seq data was obtained from Wohnhaas et al^6^ which included basal cells differentiated at an air-liquid-interface (ALI) and exposed to cigarette smoke for 4 days (acute) and basal cells exposed to cigarette smoke during the differentiation for 28 days (chronic). Basal cells were derived from 6 donors including 3 healthy non-smokers and 3 patients with COPD. **(A)** Cells were initially clustered with Seurat and then basal (KRT5), transitional, and secretory/club cells (SCGB1A1) were isolated and reclustered with Celda. The UMAPs from Celda show the assigned cell types and the four exposure conditions for each cell. **(B)** Clusters containing basal cells were identified as those expressing module L7 which contained KRT5 (**Supplementary Table 20**). **(C)** Subsequently, modules discovered in lung cancer cells were scored in this single-cell dataset by summing and normalizing the counts of genes in the modules. Modules differentially expressed between basal cells exposed to cigarette smoke versus those exposed to air were identified using LMEs that included exposure and duration as fixed effects and subject ID as a random effect (FDR < 0.05; |Log2 Fold Change| > 0.5). Only healthy donors without COPD were used for this analysis. Many detoxification and lineage-related modules associated with LUSC were significantly higher in basal cells exposed to cigarette smoke compared to unexposed basal cells. Each column in this heatmap represents the average expression of basal cells in each condition for each subject.

**Supplementary Figure 18.**
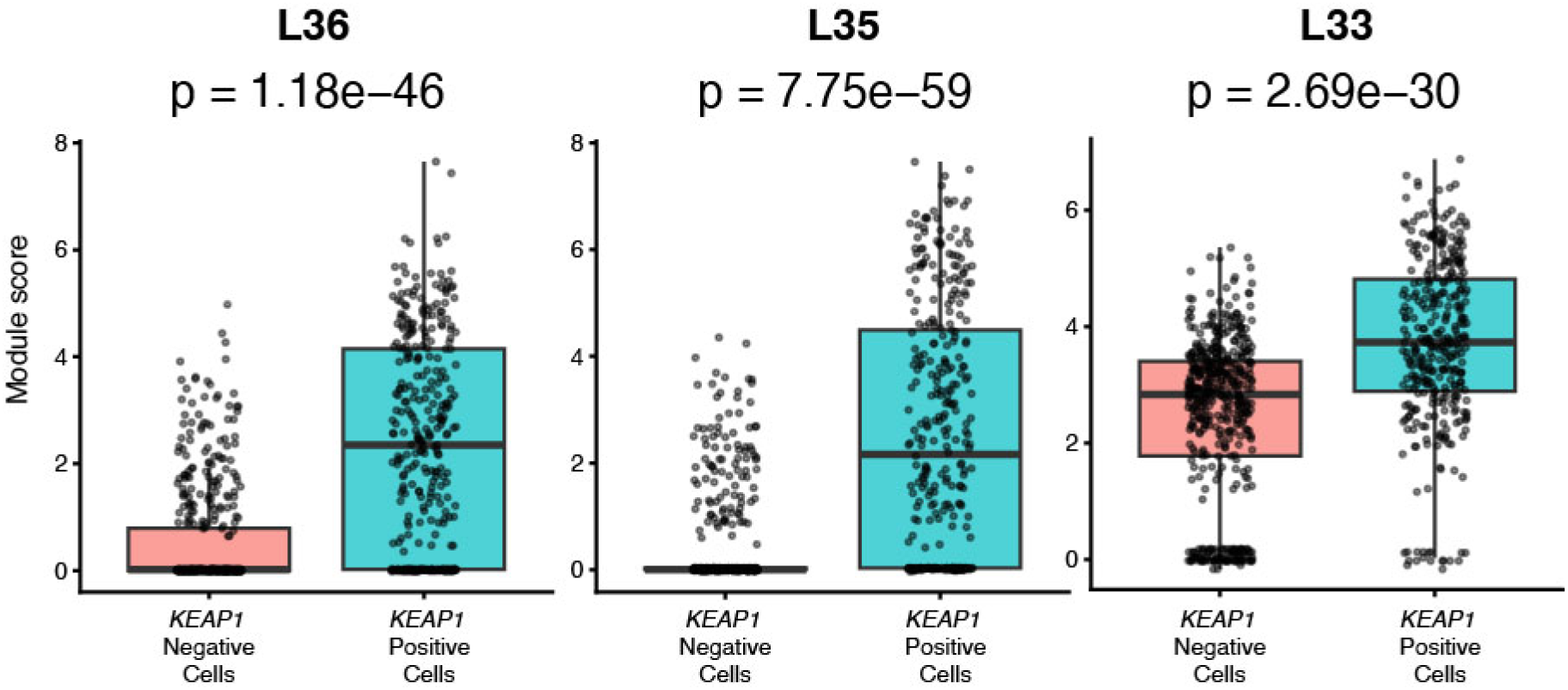
Scissor analysis of *KEAP1* mutations in LUAD. Scissor was used to integrate the LUAD TCGA bulk RNA-seq data and the LUAD cells from our atlas. Modules L33, L35, and L36 were significantly higher in Scissor+ cells than Scissor-cells (Wilcoxon rank-sum test; p < 0.001), further supporting the putative role for this gene in regulating the expression of detoxification and xenobiotic response programs.

**Supplementary Figure 19.**
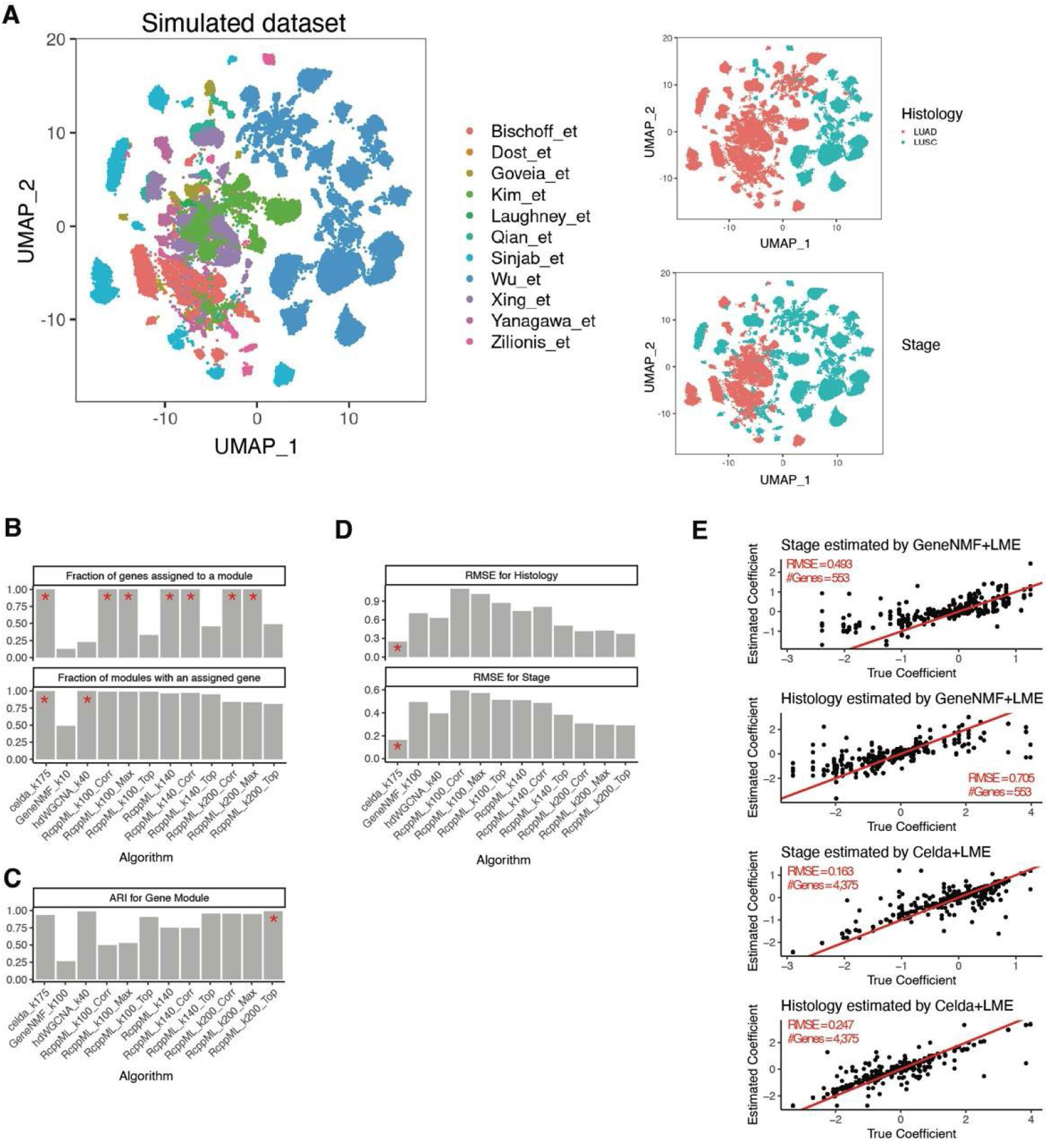
Benchmarking the ability of different tools to perform GPAS. **(A)** A synthetic dataset was created using the coefficients for 175 modules from the LMEs derived from the lung single cell atlas. For each module, the counts for 25 genes were generated by sampling the total module count from a Dirichlet distribution (25 genes x 175 modules = 4,375 genes total). UMAPs of the simulated cells generated by celda are shown and colored by study, tumor type, or stage. This structure is similar to the original real-world data in which the cells from each tumor tend to cluster separately but there are also larger global effects of study, histology, and stage. Performance metrics are shown for 4 different gene module detection tools with various parameters including: **(B)** the total number of genes assigned to a module and number of modules with an assigned gene; **(C)** the ARI of module assignments; **(D)** the RMSE assessing the accuracy of recovered coefficients from the LME model. Red asterisks indicate which tool(s) ranked the highest in each metric. **(E)** Examples of recovered coefficients for GeneNMF and Celda.

**Supplementary Figure 20.**
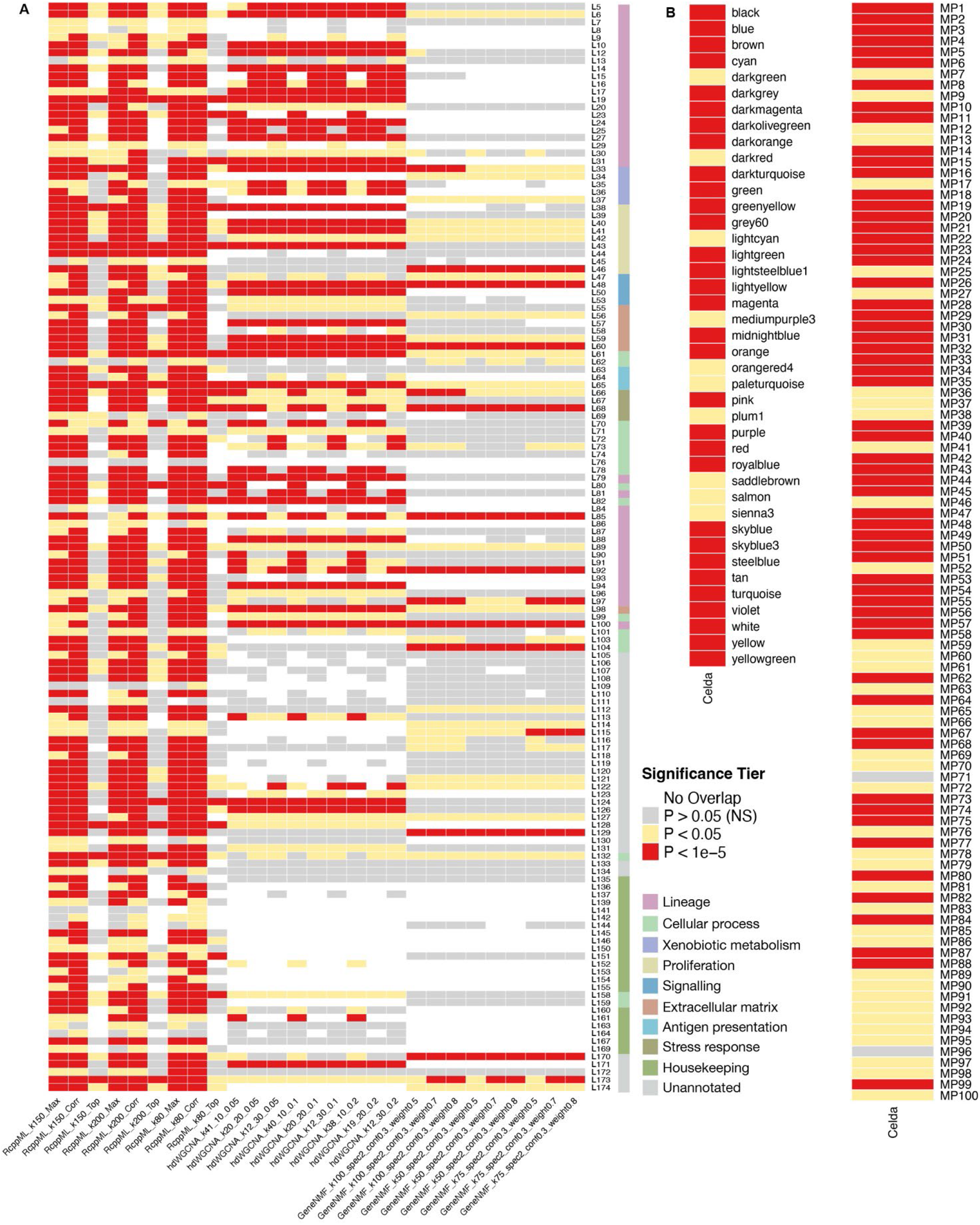
Celda modules recovered by geneNMF, hdWGCNA, and RcppML NMF. **(A)** Heatmap showing overlap between celda modules and modules recovered across all runs of the three methods. Where a celda module overlapped multiple programs within a run, the program with the highest significance was retained. Rows represent celda modules; columns represent nine runs per method. Nine out of 81 geneNMF runs are shown. Column names include the parameter set used. For RcppML NMF: number of factors {80, 150, 200} and gene-assignment method {Max, Corr, Top}. For hdWGCNA: number of modules detected {41, 20, 12, 40, 19, 38}, minModuleSize {10, 20, 30}, and mergeCutHeight {0.05, 0.1, 0.2}. For geneNMF: number of metaprograms {50, 75, 100}, specificity.weight {2}, min.confidence {0.3}, and weight.explained {0.5, 0.7, 0.8}. Cells are colored by significance of gene overlap: white, no overlap; grey, non-significant overlap; yellow, moderately significant (P < 0.05); red, highly significant (P < 1e−5). The annotation bar indicates the functional category of each celda module. **(B)** Left: celda recovered 100% of modules detected by a single hdWGCNA run (minModuleSize = 10, mergeCutHeight = 0.05, 41 modules). Right: celda recovered 98% of programs identified by a single geneNMF run (nMP = 100, specificity.weight = 2, min.confidence = 0.3, weight.explained = 0.8).

**Supplementary Figure 21.**
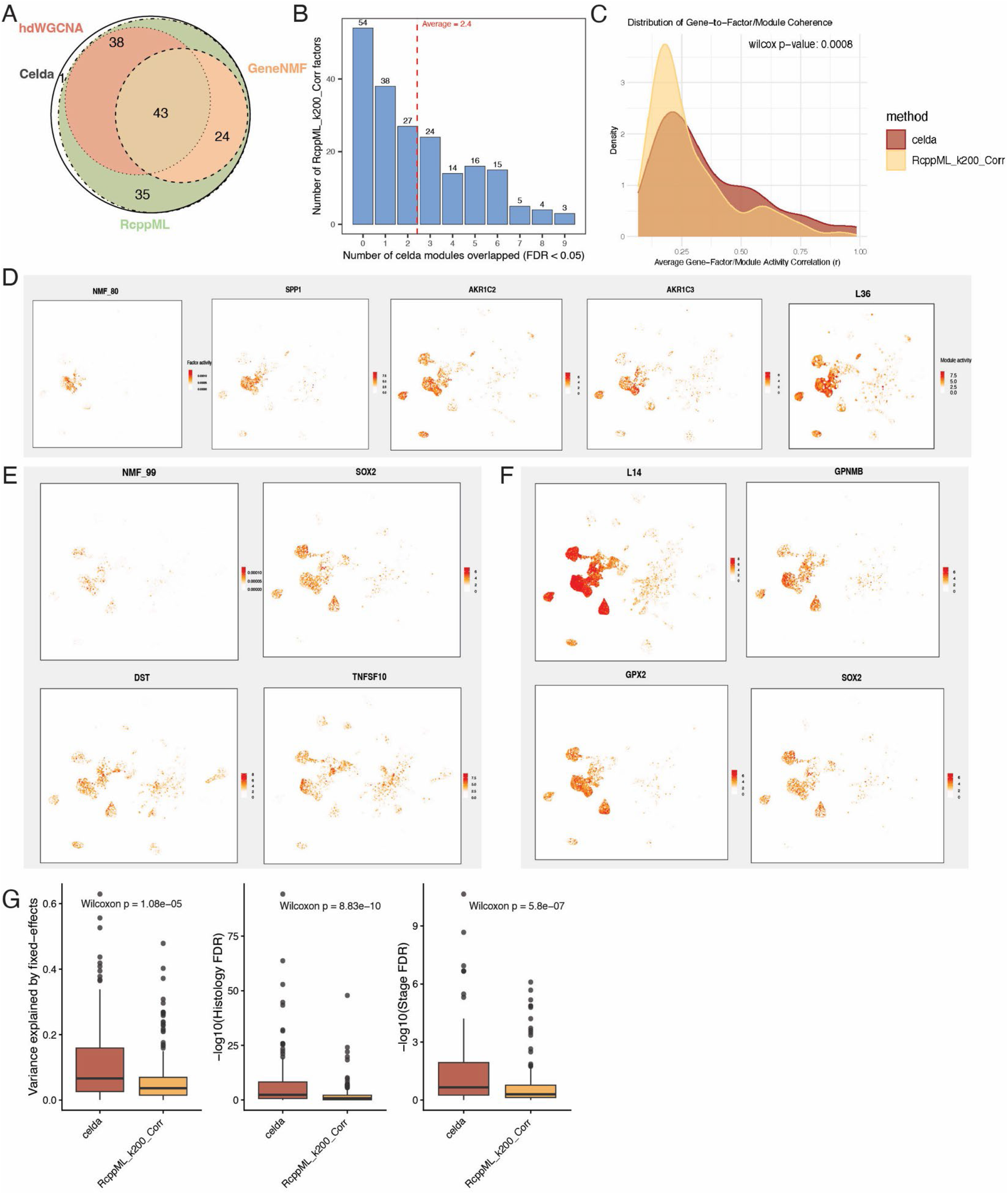
Celda modules show stronger coherence and phenotype association compared to RcppML_k200_Corr factors. **(A)** Euler diagram showing the celda modules (n=141) significantly overlapped by programs from geneNMF (runs=81), hdWGCNA (runs=9), and RcppML NMF (runs=9). Each celda module was tested against every program from any run of a given method and counted as recovered if at least one program reached statistical significance (FDR < 0.05). **(B)** Distribution of the number of celda modules overlapping with each of the 200 RcppML_k200_Corr factors. The red line represents average number of modules each factor overlapped with. **(C)** Density distribution of average gene-to-module (celda; n=171) or gene-to-factor (RcppML_k200_Corr; n=183) activity correlations, compared by Wilcoxon rank-sum test. Modules/factors with single gene were excluded. **(D)** UMAP showing activity of factor NMF_80 and its corresponding celda module L36, alongside expression of the top three genes shared between them (SPP1, AKR1C2, AKR1C3). **(E)** UMAP representation of NMF_99 alongside expression of its top three genes (SOX2, DST, TNFSF10). **(F)** UMAP representation of the corresponding celda module L14 alongside expression of its top three genes (GPNMB, GPX2, SOX2). **(G)** Boxplots comparing linear mixed effects model results for celda modules (n=175) and RcppML_k200_Corr factors (n=200). Left: Fraction of variance explained by fixed-effects (histology and stage). Middle: Significance of the histology effect. Right: Significance of the stage effect.

**Supplementary Figure 22.**
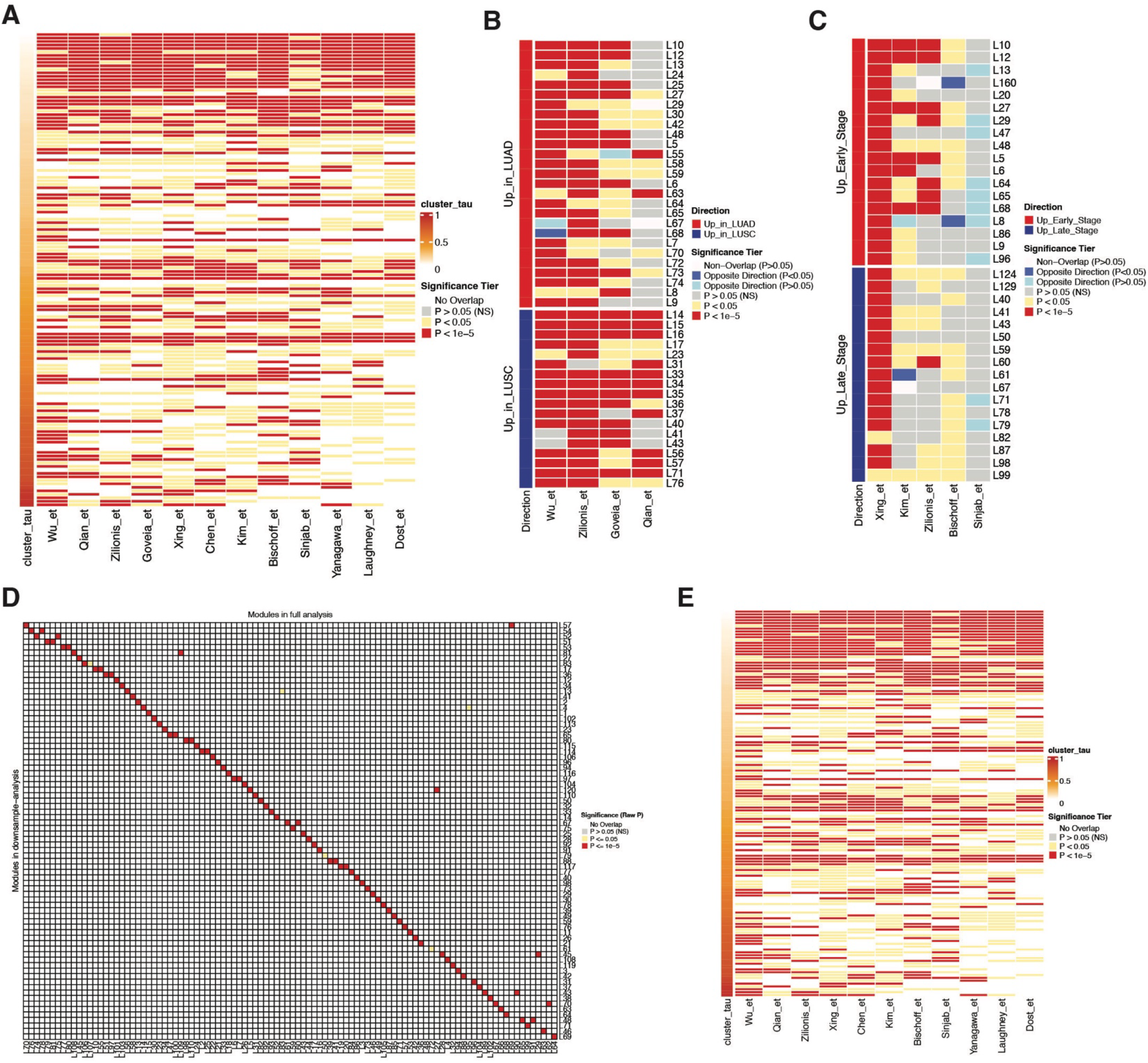
Sensitivity analyses comparing module recovery and clinical associations between pooled and non-pooled study designs and after downsampling cells per tumor. **(A)** Overlap between modules from the pooled analysis (rows; modules with at least 5 genes) and modules learned independently within each study (columns). For each pooled module, one-sided hypergeometric p-values against the top 30 genes of every study-specific module (minimum overlap of two genes) were combined using Fisher’s method to yield one value per study. The left annotation shows the cell-cluster specificity (*τ*) of each pooled module. Modules recovered in only a subset of studies were typically those restricted to specific cell clusters in the pooled analysis (i.e., high *τ*), as expected when the corresponding subpopulations are present in only some samples. **(B–C)** Preservation of **(B)** histology (LUAD vs. LUSC) and **(C)** stage (early vs. late, LUAD only) associations within individual studies. For each pooled module associated with histology or stage, the best-overlapping study-specific module was tested for the same association within that study using an LME. Colors indicate the significance and direction of the within-study association relative to the pooled result; white indicates that no overlapping module was detected. Histology associations were highly reproducible across the four studies containing both LUAD and LUSC. Stage associations were concordant in every module in Xing et al., the study with the most LUAD tumors, and in the majority of modules in at least one additional study, with attenuated significance and occasional opposite directions in studies with few tumors per stage group, consistent with the limited power of individual cohorts. **(D)** Overlap between modules learned from all cells in Wu et al. (columns) and from the same tumors after capping each at 500 cells (rows). 99 of 100 modules were recovered along the diagonal, indicating that module structure within a single study is stable to cell-number normalization. **(E)** As in (A), for study-specific modules learned after capping each tumor at 500 cells. Recovery of pooled modules was nearly identical to that observed without downsampling, demonstrating that module discovery is robust to cell-number imbalance across tumors even in the absence of cross-study pooling.

